# Multi-year drought strengthens positive and negative functional diversity effects on tree growth response

**DOI:** 10.1101/2024.11.21.622593

**Authors:** Hernán Serrano-León, Haben Blondeel, Paula Glenz, Johannes Steurer, Florian Schnabel, Lander Baeten, Joannès Guillemot, Nicolas Martin-StPaul, Georgios Skiadaresis, Michael Scherer-Lorenzen, Damien Bonal, Matthieu Boone, Renaud Decarsin, Arsène Druel, Douglas L. Godbold, Jialiang Gong, Peter Hajek, Hervé Jactel, Julia Koricheva, Simone Mereu, Quentin Ponette, Boris Rewald, Hans Sandén, Jan Van den Bulcke, Kris Verheyen, Ramona Werner, Jürgen Bauhus

## Abstract

- Mixed-species forests are proposed as strategy to increase the resistance and resilience of forests to drought stress. However, evidence suggest that increasing tree species richness does not consistently enhance tree growth responses to drought. Moreover, tree diversity effects under unprecedented multiyear droughts remain uncertain, calling for a better understanding of the underlying processes.
- Here, we used a network of planted tree diversity experiments to investigate how drought-induced growth responses of individual trees are influenced by neighborhood tree diversity and the functional traits of the focal tree species. We analyzed tree cores (948 trees across 16 species) from nine experiments across Europe featuring gradients of tree species richness (1–6 species), which experienced severe droughts in recent years. Radial growth response to drought was quantified as tree-ring biomass increment using X-ray computed tomography. We applied hydraulic trait-based growth models to analyze single-year drought responses across all sites and site-specific responses during consecutive drought years for six sites as a function of neighborhood tree diversity.
- The large variability in tree growth responses to a single-year drought was partially explained by the focal species’ hydraulic safety margin (representing species’ drought tolerance) and drought intensity, but independent of neighborhood species richness or functional trait diversity. However, tree diversity effects on growth responses strengthened during consecutive drought years and were site-specific, with contrasting direction (both positive and negative). This indicated opposing pathways of diversity effects under consecutive drought events, possibly resulting from competitive release or greater water consumption in diverse mixtures.
- We conclude that tree diversity effects on growth responses to single-year droughts may differ considerably from responses to consecutive drought years. Our study highlights the need to consider trait-based approaches (specifically, hydraulic traits) and tree neighborhood scale processes to understand the multifaceted growth responses of tree mixtures under prolonged drought stress.

## Introduction

Drought events of increased frequency, intensity, and duration are globally causing large-scale forest dieback and mortality (Schuldt et al. 2020; Senf et al. 2020; Hartmann et al. 2022). Under intensifying climate change, multi-year extreme drought events are expected to become more frequent in the future, as foreshadowed by the record-breaking 2018-2020 drought in Central Europe (Hari et al. 2020; Rakovec et al. 2022; Zscheischler and Fischer 2020). Consecutive drought years can exacerbate initial drought impacts (Anderegg et al. 2020; Schnabel et al. 2022). Moreover, drought impacts can persist for several years following a drought event, so-called drought legacy effects (Kannenberg et al. 2020; Anderegg et al. 2015; Wu et al. 2018). Abiotic (accumulated water deficit) and biotic (vegetation response) legacy effects can lead to increased vulnerability to subsequent droughts (Kannenberg et al. 2019b; Müller and Bahn 2022; Bastos et al. 2021). Tree and ecosystem responses to prolonged drought effects can differ largely depending on the drought characteristics (Anderegg et al. 2013; Huang et al. 2018; Guisset et al. 2024), site conditions (Bose et al. 2024; Gazol et al. 2020b; Kannenberg et al. 2019a), and the drought-tolerance of tree species (Gazol et al. 2020a). Given the unprecedented nature of multiyear drought events, there is a large uncertainty about the efficiency of adaptive forest management strategies to face these events (Himes et al. 2023).

Increasing tree diversity in forests has been suggested to foster the resistance, resilience and adaptive capacity of forests to cope with drought impacts (Schnabel et al. 2021; Jucker et al. 2014; Messier et al. 2022). However, increasing tree species richness alone might not necessarily improve tree ability to face increasing drought stress. Studies reported that the effect of tree diversity can vary from positive or neutral effects under mild drought stress, to negative under severe conditions droughts (Haberstroh and Werner 2022; Forrester et al. 2016). The impacts of tree diversity on the drought response might also differ between single-year droughts and consecutive droughts, characterized by cumulative soil water deprivation and legacy effects. However, since most research has focused on single-year drought events, the role of tree diversity in buffering consecutive drought impacts remains unclear.

To gain a better understanding, we propose two contrasting conceptual pathways of tree diversity effects under consecutive drought conditions. In the first pathway (Figure 1a), increased functional diversity buffers the impacts of the initial drought and this positive diversity effect becomes more pronounced under consecutive droughts. This aligns with the stress-gradient hypothesis, where facilitative interactions outweigh competition during increased stress (Bertness and Callaway 1994). Functional diversity can reduce competition for soil water and mitigate drought stress through multiple mechanisms related to resource partitioning and facilitation, e.g. complementary stomatal regulation or root stratification strategies (Loreau and Hector 2001; Trogisch et al. 2021; Mas et al. 2024b), hydraulic redistribution (Forrester and Bauhus 2016; Bauhus et al. 2017), or improved microclimate through shading and evapotranspirative cooling (Beugnon et al. 2024; Zhang et al. 2022; Schnabel et al. 2024b). These mechanisms allow trees in diverse mixtures to maintain hydraulic function, growth, and carbon reserves, reducing vulnerability to both initial and subsequent droughts (McDowell et al. 2022; Mas et al. 2024a). Functionally diverse forests may also maintain their buffering capacity and ecosystem function over time by stabilizing the community (Mahecha et al. 2024; Schnabel et al. 2021; Loreau et al. 2021). In the second pathway (Figure 1b), increased functional diversity may intensify drought stress, specifically under consecutive drought years. This pathway considers that diversity effects depend on the stress tolerance and competitive ability of interacting species, becoming negative at the extremes of resource-driven stress gradients (Maestre et al. 2009; Soliveres et al. 2015). Under increased drought stress, complementary resource-use strategies in diverse mixtures might lead to higher exploitation of limited soil water and increased interspecific competition (Haberstroh and Werner 2022; Forrester et al. 2016). Increased water consumption in mixtures may also result from overyielding and higher leaf area during favorable conditions preceding drought (Jump et al. 2017; Jacobs et al. 2021) and selection effects (Grossiord 2020; Forrester and Bauhus 2016). Negative tree diversity effects might become more pronounced over time as drought stress is amplified by abiotic and biotic legacy effects (Kannenberg et al. 2020; Mahecha et al. 2024; Shovon et al. 2024). Ultimately, this could lead to the performance loss of drought-sensitive species in mixtures as competition for limited water intensifies (Jacobs et al. 2021; Sachsenmaier et al. 2024). Whether tree diversity mechanisms positively or negatively influence the growth responses under consecutive drought conditions might strongly depend on tree species identity, mixture composition and site context (Ratcliffe et al. 2017; Grossiord et al. 2014b; Pardos et al. 2021).

**Figure 1.**
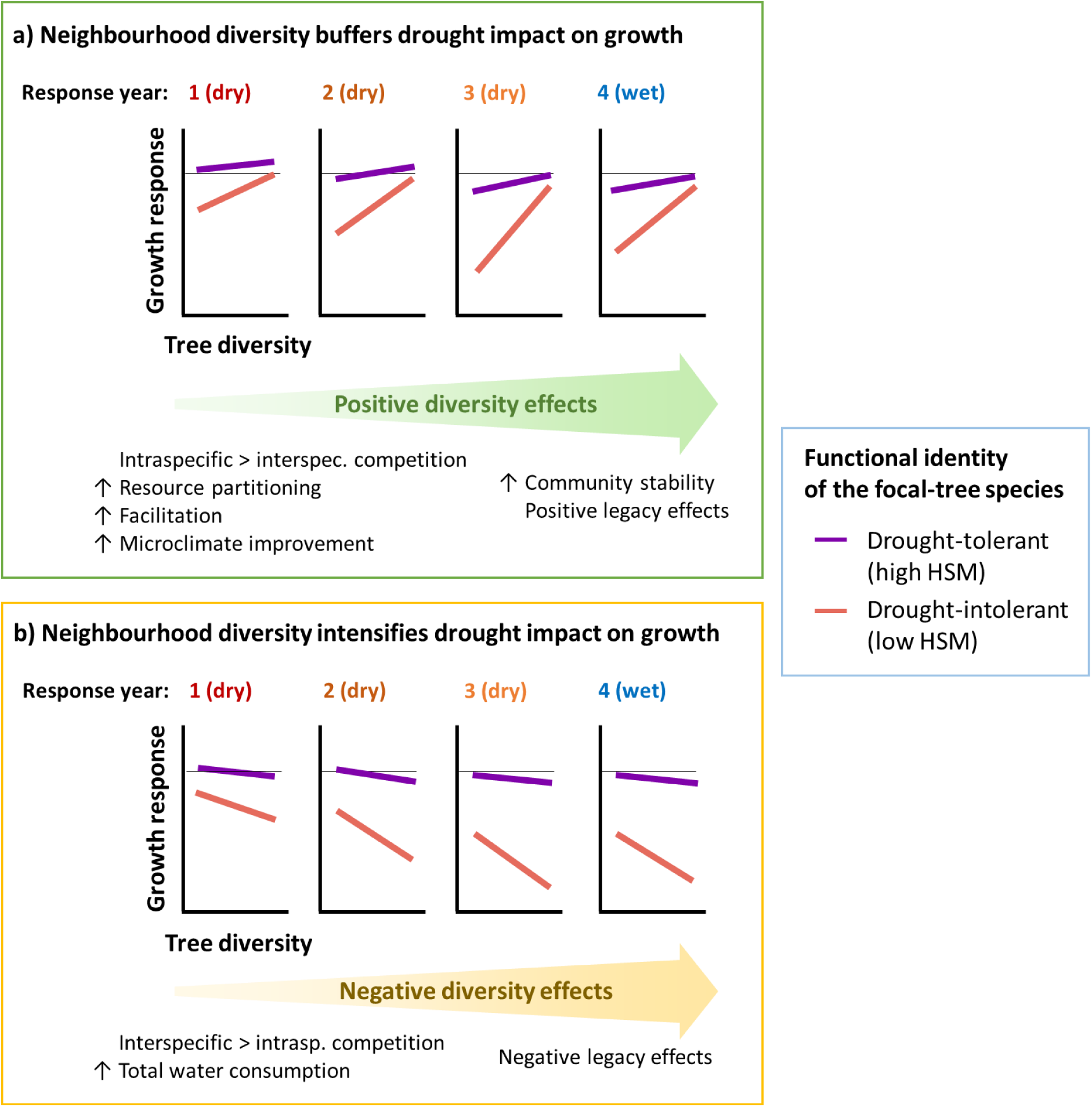
Theoretical pathways for the role of neighborhood tree diversity and drought tolerance of the focal-tree species (quantified as hydraulic safety margin HSM) for determining growth responses under consecutive drought years. In the example, an initial drought is followed by two consecutive drought years and a post-drought year with wet conditions, similar to the drought conditions analyzed in this study. Growth responses values below the horizontal line correspond to growth reductions relative to the reference pre-drought year. In the first pathway (a), functional diversity has an increasingly positive effect buffering the drought impacts on the growth response under consecutive drought stress. In the alternative pathway (b), functional diversity causes an intensification of water stress under consecutive drought years. In both cases, drought-intolerant species are particularly sensitive to drought and tree diversity effect.

Contrasting reports of tree diversity effects on drought-induced growth responses indicate that the underlying biological mechanisms remain unclear. Most studies assess tree diversity effects solely by species richness at the stand level, overlooking the functional diversity at the local neighborhood scale, where facilitative and competitive interactions actually emerge (Fichtner et al. 2017; Trogisch et al. 2021). Recent advances in plant physiology highlight the role of plant hydraulic traits in mediating these responses (Torres-Ruiz et al. 2024). Among these traits, hydraulic safety margin (HSM) is a strong predictor of drought tolerance (Martínez-Vilalta and Garcia-Forner 2017; Choat et al. 2012), with higher HSM associated with lower risk of tree hydraulic failure under drought (Anderegg et al. 2016; Martin-StPaul et al. 2017). Higher hydraulic diversity in forest communities could enhance resilience and stabilizes ecosystem productivity during droughts (Anderegg et al. 2018; Schnabel et al. 2021). Functional traits of focal tree species can also influence how tree diversity affects drought responses. Drought-vulnerable species have been reported to benefit from species-rich neighborhoods, while drought-tolerant species can be less sensitive to diversity effects (Fichtner et al. 2020; Schnabel et al. 2024a; Sachsenmaier et al. 2024; Göransson et al. 2016). Yet, these single-site studies from few experimental sites cannot elucidate the potential context dependency of neighborhood tree diversity effects on tree responses to drought. A comprehensive analysis following a trait-based neighborhood approach across multiple environmental conditions is still missing to understand how tree diversity and functional trait identity modulate tree susceptibility to drought.

Here, we present the first study to evaluate the effect of tree functional diversity on tree growth responses to single-year and multi-year droughts across nine planted tree diversity experiments in Europe. The examination of these experiments allowed us to study tree diversity effects in young tree communities growing under controlled experimental conditions while simultaneously covering varied environmental conditions. We quantified radial growth responses in terms of biomass increment using dendrochronological analysis. We used trait-based models to test whether the drought-induced growth responses were determined by the tree diversity of its neighborhood and the functional identity of the focal tree. We captured the effect of functional identity in terms of drought-tolerance represented by the HSM of the focal-tree species, and the effect of tree diversity of the neighborhood in terms of tree species richness and functional diversity of HSM. Specifically, we aim to answer the following questions:

(Q1) Does tree diversity have a consistently positive, negative or neutral effect on the growth response to a single-year drought across tree diversity experiments in contrasting environmental conditions?

(Q2) Are tree diversity effects maintained, intensified, or reduced under consecutive drought years compared with the initial drought response?

(Q3) Does drought tolerance of focal trees modulate the effect of tree diversity on growth responses to single-year droughts and consecutive drought years?

## Methods

### Study sites and sample collection

We studied nine tree diversity experiments across Europe that are part of the global TreeDivNet network of forest biodiversity experiments (https://treedivnet.ugent.be/; (Verheyen et al. 2016; Paquette et al. 2018). The studied experiments included B-Tree (Austria), BIOTREE-Kaltenborn (Germany), FORBIO-Gedinne, FORBIO-Hechtel-Eksel, FORBIO-Zedelgem (Belgium), IDENT-Freiburg (Germany), IDENT-Macomer (Italy), ORPHEE (France), and Satakunta (Finland) (Figure 2, Table S1). The experiments cover a wide range of climatic conditions comprising Mediterranean, continental, temperate oceanic, and boreal climates. All experiments use a site-specific pool of tree species adapted to local climate and soil conditions. At each site, all the species were planted in monocultures and in mixtures with varying degrees of species richness in a replicated randomized design that allows separating effects of tree identify from tree diversity on forest functioning and controls for confounding effects of environmental variation (Verheyen et al. 2016; Scherer-Lorenzen et al. 2007). At the time of sampling, all species combinations in all experiments had developed beyond canopy closure at least for several years. The tree age ranged between eight (IDENT-Macomer) and 23 years (Satakunta). Within each experiment, we selected a subset of species and mixtures compositions with contrasting functional diversity exhibiting different ecological strategies (Table S2). Within each experiment, each species was sampled in different species compositions that differ in tree species diversity: monocultures (1 species), simple (2-species) and more diverse mixtures (3- to 6-species, depending on the site). For each experiment, 10 individuals per species and composition were selected as focal trees. Each selected composition was represented by two plot replicates per experiment (except BIOTREE-Kaltenborn). Focal trees were selected according to the following criteria: (1) dominant or co-dominant trees within the species cohort to reduce the effect of different light availability on growth; (2) healthy trees with straight single stems to avoid sampling reaction wood; (3) trees in or near the plot center to avoid edge effects; (4) most of direct tree neighbors alive to avoid confounding density effects, and (5) direct neighborhood representing the species composition of the plot to maximize interspecific tree-tree interactions. Sampled trees of a given plot and species had comparable sizes. We sampled a total of 1,424 focal trees from 21 species (Table S2).

**Figure 2.**
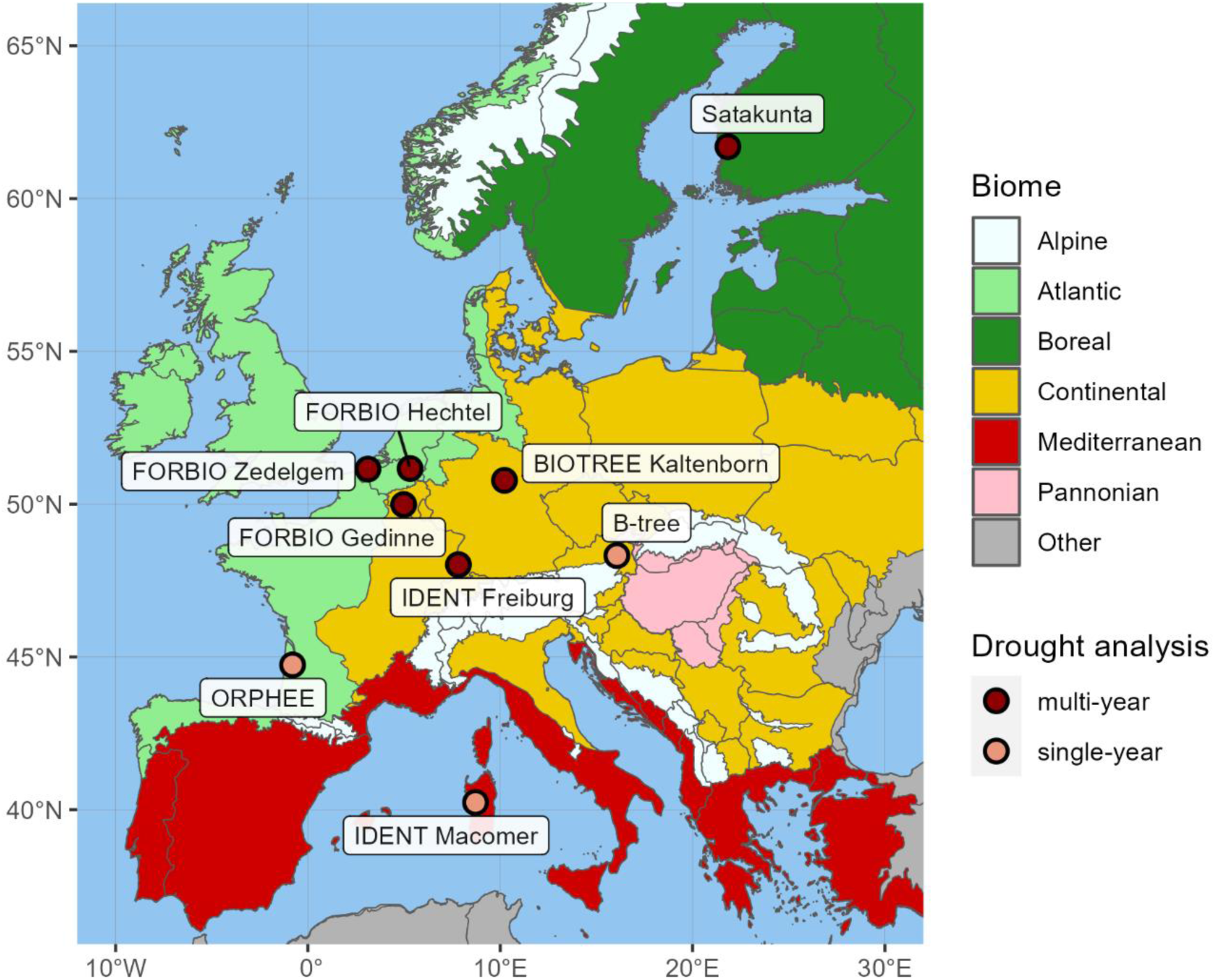
Location of experimental sites. All sites were analyzed for growth responses during a single-year drought (research question Q1) and six sites were analyzed during consecutive drought years (Q2, darker points). The biogeographical region layer is based on Cervellini et al. (2020). Background information on experiments (i.e. planting design, climate, soil characteristics) is detailed in Table S1. A list of mixture compositions considered in the analysis is detailed in Table S2.

We collected one increment core from each focal tree using a standard Pressler increment borer with an inner core diameter of 5 mm (Haglöf, Sweden). We took samples at a basal coring height of 30 cm to maximize the tree-ring series length. We cored trees primarily from the southern side to avoid potential eccentricity of tree rings from swaying in westerly winds, or perpendicular to the lean direction to avoid sampling reaction wood (Tumajer and Treml 2019; Visser et al. 2023). We collected all samples at the end of the 2021 growing season (November-December).

### Drought selection and drought-induced growth responses

Droughts were identified as periods of extreme deficit in water availability following a climate-based approach with the use of site-specific data (Schwarz et al. 2020; Slette et al. 2019). We obtained climate data from the ERA5Land of the Copernicus Climate Data Store (Muñoz-Sabater et al. 2021). We characterized drought conditions considering the Standardized Precipitation Evapotranspiration Index (SPEI) and relative extractable water (REW). First, we calculated SPEI from the local monthly climatic water balance (precipitation minus potential evapotranspiration) over a selected moving time window using the R-package SPEI (Vicente-Serrano et al. 2010; Beguería and Vicente-Serrano 2017). We considered ‘SPEI12 December’ for the annual conditions and ‘SPEI6 April-September’ for the growing season. We calculated SPEI over the reference 30-year period (1991-2021) to determine abnormally dry years for each experiment location, using the threshold −1.28 as the 10% quantile of all values following the drought classification of the SPEI global drought monitor (Agnew 2000; Beguería et al. 2022). Then, we calculated the minimum monthly REW during the growing season to characterize the drought intensity in terms of plant drought stress experienced by trees with the plant hydraulics model SurEau (Ruffault et al. 2013). The model was applied at each site using vegetation parameters specific to one species representative of the biome and the local species pool of each experiment as in Blondeel et al. (2024). Details on the calculation of drought indices are explained in Supplementary Method 1, input variables used in the SurEau simulations are detailed in Table S3. The different drought indices showed comparable temporal patterns and high correlation within each site (Figure S1, Figure S2).

Identification of recent drought years focused on the last five years before sample collection. To answer Q1, we focused our analysis on the growth response to the first drought year (hereafter, growth response year 1 or Resp_yr 1) in each experiment with severe drought conditions (i.e. SPEI12 < −1.28). Pre-drought reference year(s) were determined as the closest preceding year (or two years) to the corresponding first drought characterized by normal or wet climatic conditions (i.e. SPEI >/≈ 0). To answer Q2, growth responses were calculated for six experiments (FORBIO-Gedinne, FORBIO-Hechtel-Eksel, FORBIO-Zedelgem, BIOTREE-Kaltenborn, IDENT-Freiburg, and Satakunta) experiencing a multiyear drought event in 2018-2020. In these experiments, the initial drought year was severely dry (Resp_yr 1 with SPEI < - 1.28) and followed by two consecutive years with moderate drought conditions (Resp_yr 2-3 with SPEI ≈ −1). These sites experienced a post-drought year in 2021 characterized by normal or wet climatic conditions (Resp_yr 4 with SPEI >/≈ 0), which was included in the analysis to consider potential drought legacy effects. The final selection of study drought years for each site is shown in Table S4 and Figure S3.

### Radial growth measurements

We obtained annual series of tree-ring width (TRW) and mean wood density using X-ray micro-Computed Tomography (µCT) with the HECTOR µCT scanner (Masschaele et al. 2013) at the Ghent University Centre for X-ray tomography (UGCT; http://www.ugct.ugent.be). Before scanning, increment cores were dried at room temperature for one month to be in balance with the scanner room environment (20 °C, relative humidity 34 %) and mounted in master sample holders for batch scanning. We scanned cores at an approximate voxel pitch resolution of 50 μm following the workflow for µCT densitometry detailed in de de Mil et al. (2016). Reconstructions of the scanned sample batch were performed using the Octopus Reconstruction software (Vlassenbroeck et al. 2007). We extracted the three-dimensional images of each single core sample and converted them to density estimates using specific toolboxes (van den Bulcke et al. 2014; de Mil et al. 2016; de Mil and van den Bulcke 2023). Details of the methodology followed for the reconstruction, extraction and crossdating of tree ring series are explained in Supplementary Method 2. Rigorous quality filtering was performed to ensure that annual growth rings where correctly defined, discarding from the analysis those samples with ambiguous tree ring definitions after the crossdating processing and species with a substantial proportion of doubtful samples at a given site (> 40%). See Figure S5 for an overview of the wood anatomy of species. We analyzed a total of 948 trees from 16 species and 68 different species compositions (26 monocultures, 28 simple mixtures, 14 more diverse mixtures) (Table S2).

We calculated basal area increment (BAI, mm^2^ year^-1^) series from pith to bark based on the TRW series using the bai.in() function in the dplR package in R (Bunn 2008; Bunn et al. 2023). We estimated an indicator for radial biomass increment (hereafter BIOMinc, kg m^-1^, year^-1^) as the product of BAI and mean wood density (kg m^-3^) of the ring (Figure S8). This indicator at the individual tree level reflects the actual carbon allocation in radial growth better than TRW and BAI metrics (Camarero and Andrés 2024) and it can be considered a proxy of aboveground biomass growth when continuous inventory data in annual tree height growth is missing (Vannoppen et al. 2018; Bontemps et al. 2010). We systematically tested different transformation and detrending options for the growth variables, but all methods resulted on potential removal or overestimation of the drought-induced responses (Supplementary Method 2). Given the challenges to detrend age-effect in such short time series, we analyzed further drought-induced growth responses based on raw and undetrended series of the tree ring variables (Schwarz et al. 2020; Schnabel et al. 2024a). BIOMinc and BAI series were used as a more reliable indicator of temporal trends in radial growth for young trees than TRW (i.e. less influenced by biological age trends) (Biondi and Qeadan 2008). Finally, we quantified the drought-induced growth responses of each individual tree for a given response year as the relative growth index proposed by Lloret et al. (2011) as the ratio:

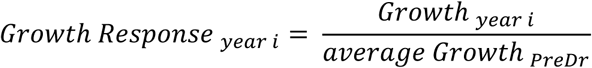

where *Growth_year i_* is the radial growth (either TRW, BAI, or BIOMinc) for the given year i (drought or post-drought year) and *average Growth_PreDr_* is the mean growth during the corresponding pre-drought reference year(s) for that site (Figure S6).

### Functional traits selection

To determine the functional identity of tree species and functional diversity of neighborhoods, we determined the species’ drought tolerance as the hydraulic safety margin (HSM_TLP_, MPa) based on both stomatal traits and hydraulic traits to encompass the spectrum of processes involved in drought tolerance but also resource use (Choat et al. 2012; Martin-StPaul et al. 2017). We defined HSM_TLP_ as the extent to which early stomatal closure protects the xylem from dysfunction during drought, calculated as the difference between the turgor loss point (TLP, Mpa, a proxy of the stomatal closure point) and the water potential at which 50% of xylem cavitates (P50, unit MPa) (HSM_TLP_ = TLP − P50). This HSM_TLP_ definition has been used to predict the risk of drought-induced tree mortality (Martin-StPaul et al. 2017; Powers et al. 2020) and is linked to stomatal control as TLP correlates with leaf water potential at stomatal closure (Bartlett et al. 2016; Brodribb and Holbrook 2003). HSM_TLP_, used as a trait characterizing species’ drought tolerance, differentiates from the traditional HSM metric (based on minimum water potential, Ψ_min_), which is considered a measure of drought stress (Choat et al. 2012). We considered species-specific mean trait values from multiple consolidated species-level datasets (see Supplementary Method 3 for the complete list of sources, and Table S6 for the final dataset of species-specific trait values). Additionally, we assessed the relation among these hydraulic traits and additional traits related to the whole-plant economics spectrum (i.e. leaf mass per area, leaf nitrogen concentration, wood density) using pairwise correlations and principal component analysis (PCA) (Supplementary Method 3, Table S6). HSM_TLP_ was associated with other drought-tolerance traits and aligned with the first PCA axis explaining most of the variation across traits (Figure S12, Figure S13). Finally, we compared the effect of HSM_TLP_ and additional functional traits on the growth responses during a single-year drought with a sensitivity analysis among different linear mixed-effect models as a function of each species-specific trait (Supplementary Method 4, Table S7, Figure S14). Based on this preliminary analysis, HSM_TLP_ was selected as the key physiological trait to quantify effects of the drought tolerance gradient on growth responses.

### Neighborhood competition and tree diversity

We defined neighborhoods of focal trees as all alive direct (first-order) neighbors and second-order neighbors that had crowns interacting with the focal tree’s crown within a certain neighborhood radius. This neighborhood radius was adapted to each site to account for the differences in planting density and design between experiments i.e. radius 2.9 m for most sites with larger planting distance, radius 1.5 m for sites with narrow planting distance (B-Tree, IDENT-Freiburg, IDENT-Macomer) (Figure S4, Table S5). We measured diameter of each focal and neighbor tree with a digital caliper at 1 mm resolution at the same coring height and direction. For each neighboring tree, we recorded its relative position to the focal tree and species identity. We quantified competition experienced by focal trees at the time of coring using these measured neighborhood data, except for IDENT-Freiburg and IDENT-Macomer for which we used inventory data from the same year. The spatial definition of the tree neighborhood around each focal tree was done using the sf R-package (Pebesma and Bivand 2023). The distance-dependent Hegyi’s index (Hegyi 1974)was calculated as a competition index for each focal tree using the following equation:

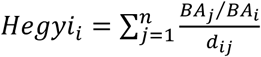

where *BA_i_* and *BA_j_* are the basal area of the focal tree *i* and each tree neighbor *j* respectively, and *d_ij_* is the distance between each tree neighbor and the focal tree, considering all *n* direct neighbor trees (from all species) within a given focal tree’s neighborhood. Alternative competition indices based on height were not considered as height data was not available for all neighborhood trees.

Neighborhood tree diversity was considered as the realized neighborhood species richness (nSR) and the functional diversity of HSM_TLP_ (FD_HSM_). FD_HSM_ for each focal tree’s neighborhood (community) was calculated as the abundance-weighted functional dispersion as described by Laliberté and Legendre (2010) using the FD R-package (Laliberté et al. 2014):

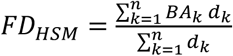

where *BA_k_* is the relative abundance of species *k* in a neighborhood with n species, calculated as the cumulative basal area, and *d_k_* is the distance in the traits space of species *k* to the weighted centroid of the neighborhood community. Functional dispersion measures the mean abundance-weighted distance of species along each trait gradient and represents the complementarity in functional strategies of co-occurring species within each neighborhood community (Laliberté and Legendre 2010).

### Statistical analysis

To answer research questions Q1 and Q3 in relation to the single-year drought response across all sites, we used linear mixed-effect models (LMMs) of the drought-induced growth responses during the first drought year to test the direct effect of neighborhood tree diversity (Q1) and its interaction with drought-tolerance (HSM_TLP_) of the focal-tree species (tree diversity x HSM_TLP_, Q3). We tested the effects of neighborhood tree diversity in separate models for the moderators nSR (m1a) and FD_HSM_ (m1b) using the same basic model structure. To account for the experimental design and differences between sites, we used a nested group-level (random) effect structure of plot nested within site. We also tested for extended versions of the model, by considering the three-way interaction effects of tree diversity, drought tolerance and the following predictors: neighborhood competition (tree diversity x HSM_TLP_ x Hegyi), tree basal area (tree diversity x HSM_TLP_ x BA), drought intensity in terms of SPEI12 (tree diversity x HSM_TLP_ x SPEI12), drought intensity in terms of SPEI6sept (tree diversity x HSM_TLP_ x SPEI6sept), and drought intensity in terms of minimum monthly REW during the growing season (tree diversity x HSM_TLP_ x REW).

To answer Q2 and Q3 regarding consecutive drought years, we used site-specific models to analyze growth responses considering the subset of six experiments that experienced a multi-year drought event. These models evaluated drought-induced growth responses across consecutive years as a function of all two-way interactions between the response year, HSM_TLP_ of the focal-tree species and the neighborhood tree diversity. Analyzing Q2 was based on the interaction tree diversity x Resp_yr, whereas analyzing Q3 was based on the interactions HSM_TLP_ x tree diversity and HSM_TLP_ x Resp_yr. We tested the effects of neighborhood tree diversity in separate models for the moderators nSR (m2a) and FD_HSM_ (m2b) using the same basic model structure. We accounted for non-linear drought responses across consecutive years including Resp_yr as a categorical fixed effect (Resp_yr 1, 2 and 3 for the consecutive drought years, Resp_yr 4 for the post-drought year). Models to address Q2 were fitted considering an intercept 0 as reference for pre-drought and included individual tree as a random effect intercept. Here, we did not consider drought intensity (i.e. SPEI) as fixed effect since it would confound the fixed effect of the consecutive Resp_yr. We used tree individual as random effect to account for the multiple Resp_yr observations per tree.

All models addressing Q1 and Q2 used a log transformation of the response variables to meet model assumptions (normality and heteroscedasticity). All models controlled for fixed effects of tree basal area (scaled to account for relative intraspecific differences in tree size within each site and species) and competition (standardized to account for relative interspecific competition within each site). Standardization of model predictors (to meet model assumptions and have comparable effect sizes across sites) and testing of alternative random structures are detailed in Supplementary Method 5. We compared the model performance and parsimony of fixed effects among the separate models based on the Akaike Information Criterion (AIC). The presentation of model results focusses on growth responses in terms of BIOMinc (Table S8). All analyses were computed using R version 4.3.1 (R Core Team 2023). All models were fitted using lme4 package (Bates et al. 2015) with restricted maximum likelihood estimation (REML). Model statistics and marginal effects were visualized using sjPlot package (Lüdecke et al. 2024).

## Results

### Growth responses during a single-year drought

We observed substantial variability in growth responses within each site, indicating that not all trees reduced growth during a single extreme drought year compared to pre-drought conditions (Figure 3, Figure S9). Models showed that drought-induced growth responses during the a single-year drought were significantly influenced by the focal-species HSM_TLP_ (p < 0.001) (Figure 3, Table S9). Overall, species with high HSM_TLP_ had less growth reduction compared to species with lower HSM_TLP_. The positive HSM_TLP_ effect was modulated by drought intensity (HSM_TLP_ x REW, p < 0.01), although this modulation was only significant when using REW, not SPEI indices (Table S9). Reduction in growth in low HSM_TLP_ species was more pronounced at sites experiencing the most extreme drought intensity (REW = 0), where differences between species with contrasting HSM_TLP_ were higher. Drought-tolerant species showed similar growth responses at sites with less extreme drought conditions (REW = 0.03), where differences with low-HSM_TLP_ species were less pronounced.

**Figure 3.**
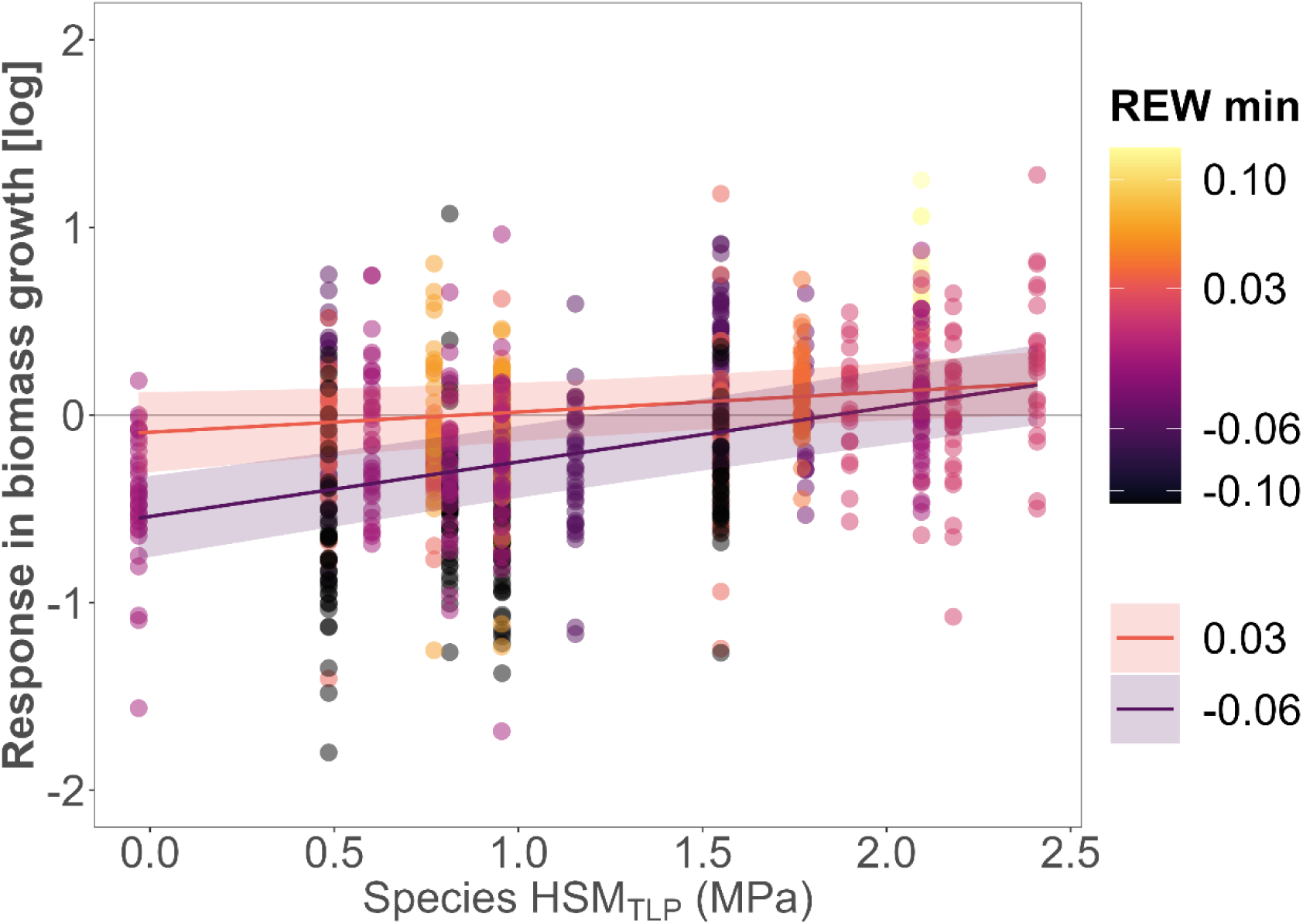
Growth response during a single-year drought and marginal effects of the hydraulic safety margin (HSM_TLP_) of the focal-tree species and the drought intensity (REW min, minimum monthly relative extractable water during growing season). Higher HSM_TLP_ (based on turgor loss point) indicate species with higher drought tolerance. Points represent tree growth responses (log transformed) in terms of radial biomass increment during the single-year drought relative to the pre-drought reference year, with values below the horizontal line y = 0 representing trees with relative growth reductions. Point colors represent the gradient of site-specific drought intensity, with darker points representing higher drought intensity (lower REW). Drought stress threshold is considered at REW = 0.4 and wilting point at REW = 0. Lines represent the marginal effects of the linear mixed-effect model fitted to REW corresponding to 1^st^ (−0.06) and 3^rd^ quartile values (0.03) of REW experienced among all sites; bands show a 95% confidence. See Table S9 for details of the fitted model. See Figure S10 for the details of the HSM_TLP_ ranges of focal species per site.

The model considering the 3-way interaction between FD_HSM_ x HSM_TLP_ x REW was the most parsimonious (lowest AIC) and with highest explanatory power (conditional R^2^ = 0.50, marginal R^2^ of fixed effects = 0.26). Nevertheless, there was no evidence that growth responses during a single-year drought were influenced by neighborhood tree diversity, either in terms of nSR or FD_HSM_ (Figure 4, Table S9). Additionally, no interaction was found between tree diversity and HSM_TLP_, as species with contrasting HSM_TLP_ responded similarly to increasing tree diversity. Likewise, there was no significant interaction effect between tree diversity and drought intensity, whether in terms of REW or SPEI indices. Single-year drought responses were also not influenced by relative neighborhood competition, either directly or in interaction with HSM_TLP_ and tree diversity (tree diversity x HSM_TLP_ x Hegyi) (Figure 4, Table S9, Table S10). In contrast, all models showed a positive direct effect of tree size (relative intraspecific differences) on the drought response (p < 0.01), although tree size did not modulate the interaction with HSM_TLP_ or tree diversity. Within each species, trees with larger BA consistently had more positive drought responses.

**Figure 4.**
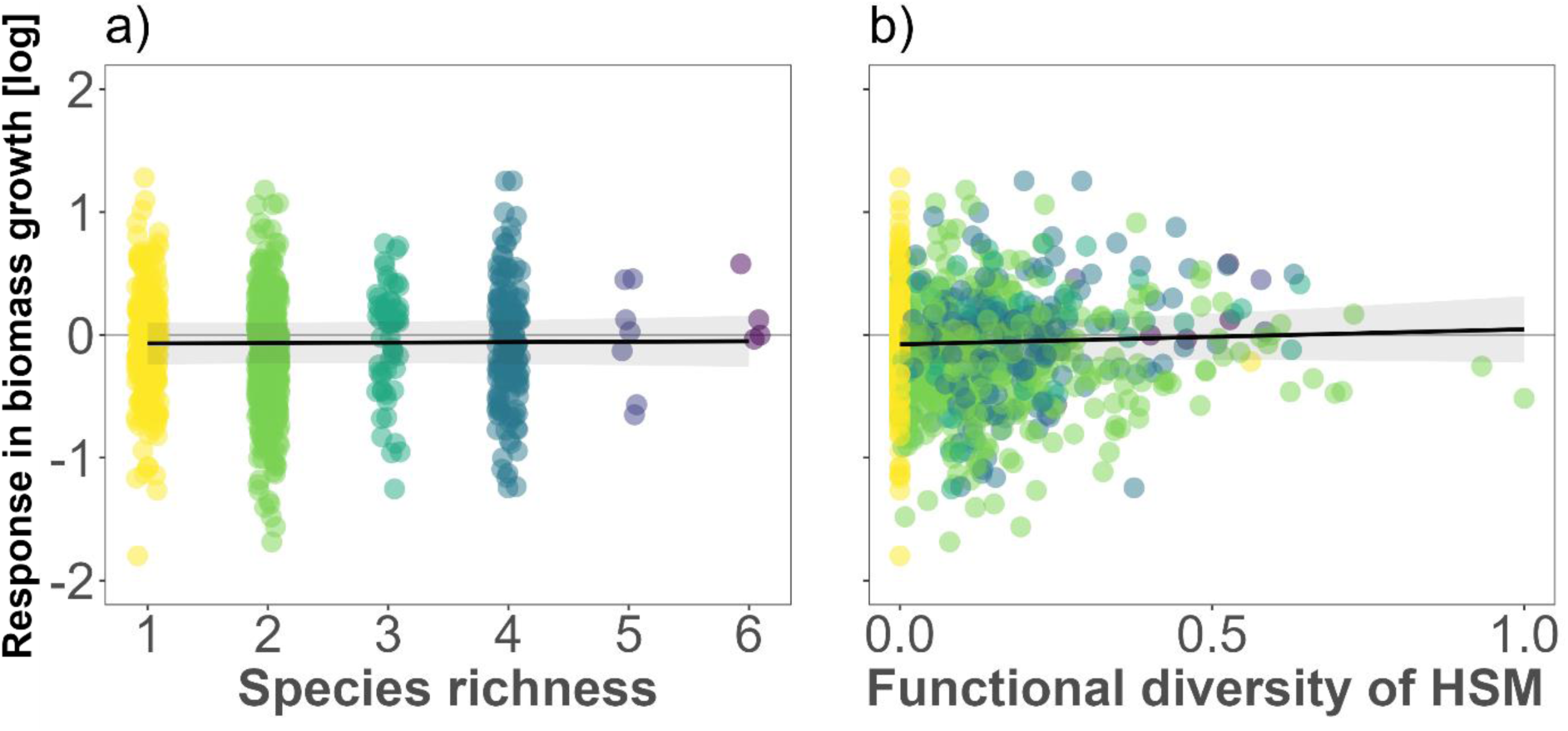
Growth response during the single-year drought and marginal effects of neighborhood tree species richness (a) and hydraulic functional diversity (b). Points represent the tree growth responses (log transformed) in terms of radial biomass increment during the single-year drought relative to the pre-drought reference year, with values below the horizontal line y = 0 representing trees with relative growth reductions. Points are colored according to the neighborhood species richness depicted in figure (a). Lines represent the marginal effects fits of the linear mixed-effect model and bands show a 95% confidence. Neighborhood functional diversity in terms of hydraulic safety margin (FD_HSM_) was standardized (via min-max normalization) across all sites to account for absolute differences between sites (Figure S16). See Table S9 for details of the fitted model.

### Growth responses during consecutive drought years

In site-specific models of multi-year drought events, we observed that the effects of neighborhood tree diversity on growth responses strengthened over consecutive drought years at most sites (interaction tree diversity x Resp_yr) (Figure 5, Table S11, Table S12). While no site showed significant diversity effects during the initial severe drought year (Resp_yr 1), diversity effects emerged during subsequent drought years and the post-drought year (Figure S3, Table S4). The direction and magnitude of diversity effects on growth responses over consecutive drought years varied by site. A positive diversity effect was evident only at IDENT-Freiburg (p < 0.001), where higher diversity increased relative growth during the following consecutive drought years (Resp_yr 2-3) and post-drought (Resp_yr 4). In contrast, a negative effect of functional diversity was evident during the third drought year (Resp_yr 3) at FORBIO-Zedelgem (p < 0.001) and Satakunta (p < 0.01), intensifying even under favorable conditions in the post-drought year at these sites (p < 0.001) and at BIOTREE-Kaltenborn (p < 0.01). A neutral effect of neighborhood tree diversity across consecutive years was found at FORBIO-Gedinne and FORBIO-Hechtel-Eksel. At all sites, tree diversity increased growth response variability. Models based on FD_HSM_ were more parsimonious (lower AIC) and with higher proportion of variance explained (marginal R2 ranging between 0.05 for Satakunta and 0.38 for IDENT-Freiburg) than models based on species richness (Table S11, Table S12).

**Figure 5.**
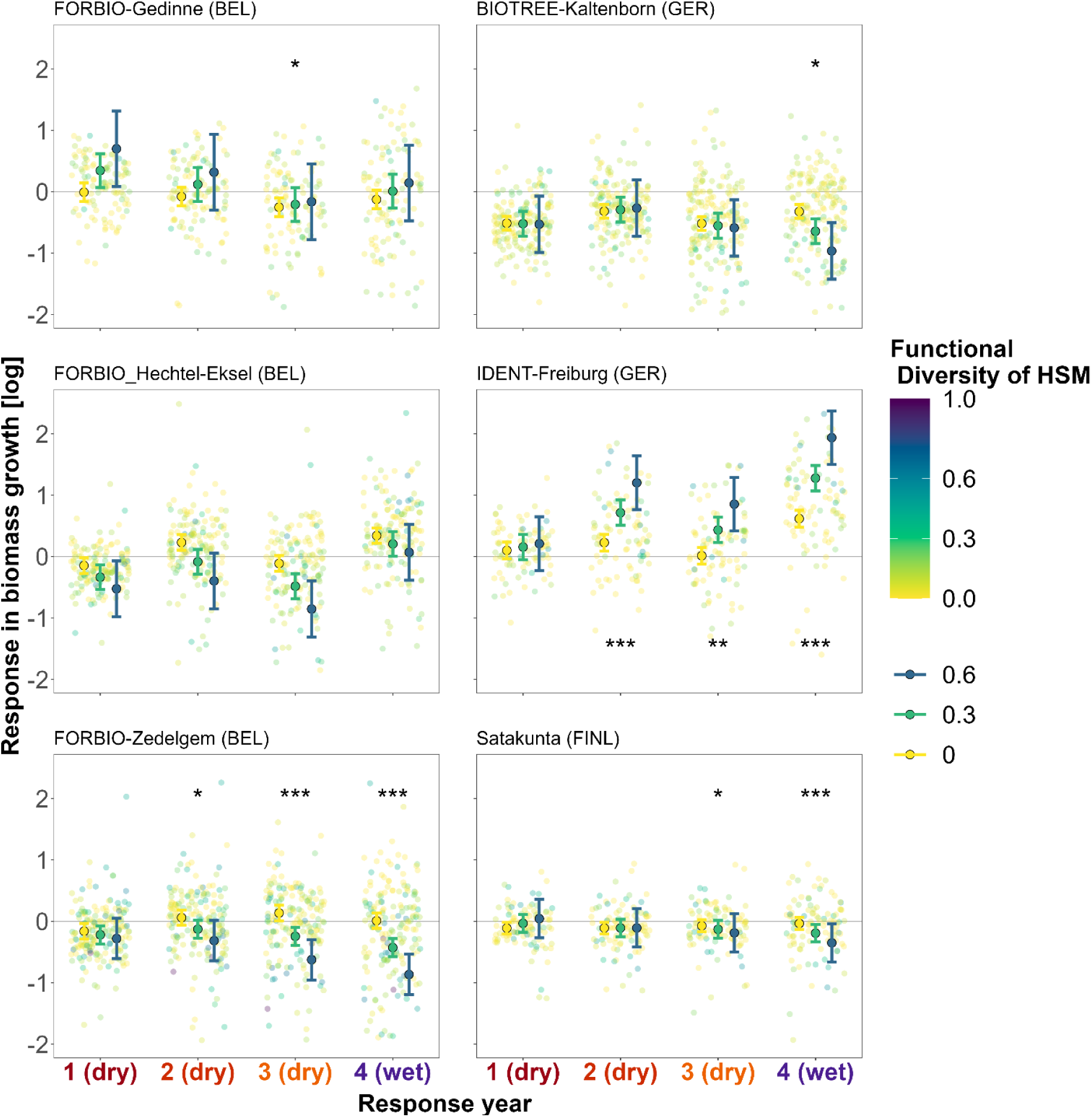
Marginal effect of neighborhood hydraulic diversity (FD_HSM_) on growth responses during three consecutive drought years (Resp_yr 1-3) and a post-drought year (Resp_yr 4) for each site-specific model. Background points represent tree growth responses (log-transformed) in terms of radial biomass increment, with values below the horizontal line y = 0 indicating trees that experienced growth reductions relative to pre-drought reference levels. Color legend represents neighborhood functional diversity in terms of hydraulic safety margin, standardized (via min-max normalization) across all sites to account for absolute differences. Marginal effects are visualized for monocultures (yellow, FD_HSM_ = 0) to intermediate- (green, FD_HSM_ = 0.3) and higher functional diversity (blue, FD_HSM_ = 0.6). Points with error bars represent the marginal effect fits with 95% confidence interval for the interaction FD_HSM_ x Resp_yr, while holding other predictors constant. Significant symbols represent significant fixed effects for the interaction FD_HSM_ x Resp_yr, with significance levels * p<0.05, ** p<0.01, *** p<0.001. See Table S12 for details on the fitted models. See Figure S16 for details on FD_HSM_ trait ranges per site.

Regarding the drought tolerance traits, no models showed significant interaction between tree diversity (nSR, FD_HSM_) and HSM_TLP_, indicating that species with contrasting HSM_TLP_ were not affected differently by increased diversity. However, the effect of HSM_TLP_ on growth responses varied depending on the site (Figure 6, Table S11, Table S12). A positive HSM_TLP_ effect across response years was only moderately evident at FORBIO-Zedelgem (p < 0.05), while most sites show no significant effect during the first drought year (unlike in the Q1 model). Nevertheless, drought-tolerant species with higher HSM_TLP_ generally showed improved growth in the post-drought year (Resp_yr 4) at FORBIO-Gedinne, FORBIO-Hechtel-Eksel (p < 0.001), IDENT-Freiburg and Satakunta (p < 0.01). Conversely, at BIOTREE-Kaltenborn, drought-tolerant species had lower relative growth during the consecutive drought and post-drought years (Resp_yr 2-4) compared to species with lower HSM_TLP_ (p < 0.001).

**Figure 6.**
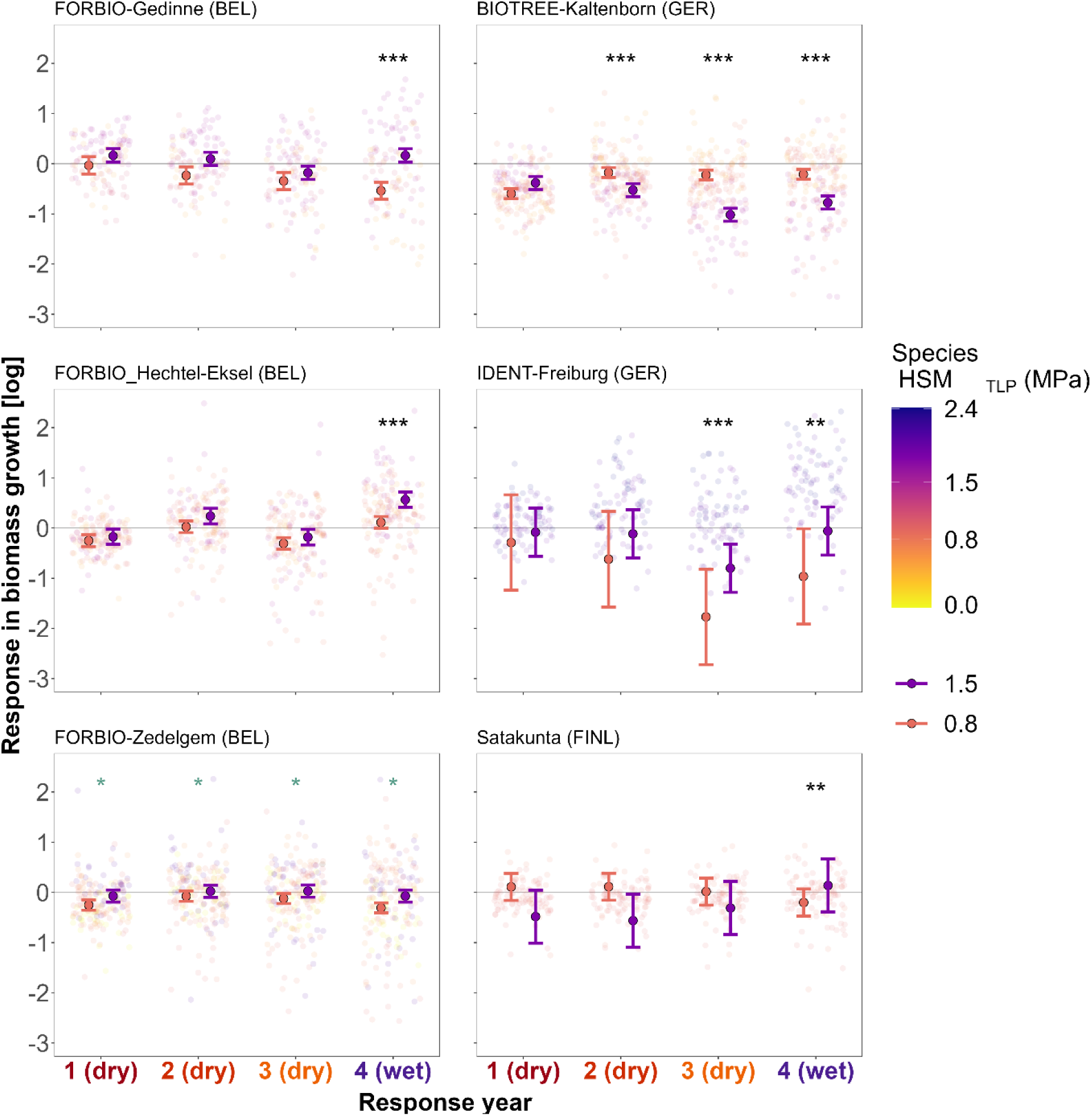
Marginal effect of hydraulic safety margin (HSM_TLP_) of focal-tree species on growth responses during three consecutive drought years (Resp_yr 1-3) and a post-drought year (Resp_yr 4) for each site-specific model. Background points represent tree growth responses (log-transformed) in terms of radial biomass increment, with values below the horizontal line y = 0 representing trees that experienced growth reductions relative to pre-drought reference levels. Color legend represents the species-specific HSM_TLP_ (based on turgor loss point). Marginal effects are visualized for species with relatively low drought tolerance (orange, HSM_TLP_ = 0.8) and high drought tolerance (violet, HSM_TLP_ = 1.5) corresponding to the 1st and 3rd quartile values of HSM_TLP_ across all analyzed species. Points with error bars represent the marginal effect fits with 95% confidence interval for the interaction HSM_TLP_ x Resp_yr while holding other predictors constant. Significant symbols represent significant fixed effects for the interaction HSM_TLP_ x Resp_yr, with significance levels * p<0.05, ** p<0.01, *** p<0.001. See Table S12 for details of the fitted models. See Figure S10 for details on HSM_TLP_ ranges of focal trees per site.

## Discussion

### Tree diversity effects on growth responses are strengthened over consecutive drought years

We found that increasing neighborhood tree diversity, whether quantified through species richness or hydraulic trait diversity, had a neutral effect on the growth responses during a single-year drought (Q1, Figure 4). This aligns with studies reporting mixed or neutral tree diversity effects during mild droughts, which can turn to negative effects during severe droughts (Haberstroh and Werner 2022; Grossiord et al. 2014b; Anderegg et al. 2013). However, we did not observe any significant interaction between tree diversity and drought intensity in radial growth responses during the first drought year. It is possible that those effects may not be evident as changes in radial growth during a single-year drought, even if physiological processes (i.e. stomatal regulation of water use) are strongly affected by drought (Jucker et al. 2017; Schnabel et al. 2022). The reason for this is that radial growth can remain stable if trees mobilize carbon reserves from previous years to sustain growth under initial drought conditions (Cailleret et al. 2017; Körner 2019; McDowell et al. 2022). However, our study data cannot ascertain whether these mechanisms played a role.

Although diversity effects were absent in the first drought year, tree diversity effects on site-specific growth responses strengthened during consecutive drought years and persisted into the post-drought year (Q2, Figure 5). Interestingly, the direction of this effect varied across sites: three sites showed increasingly negative diversity effects, two showed neutral effects, and one showed a positive effect. This corroborates the contrasting and site-specific diversity effects reported in earlier studies (Ratcliffe et al. 2017; Forrester et al. 2016; Grossiord 2020). Notably, our study is unique in showing these different response patterns across multiple experimental sites with manipulated gradients of tree species richness. The observed strengthening of tree diversity effects during consecutive years could indicate an intensification of tree-tree interactions under prolonged drought stress (Forrester et al. 2016; Soliveres et al. 2015; Maestre et al. 2009) or carry over diversity effects through drought legacy mechanisms (Anderegg et al. 2015; Vilonen et al. 2022; Bastos et al. 2020).

While the observed site-specific diversity effects during multiyear drought supported both proposed theoretical pathways (Figure 1), the underlying mechanisms driving these effects may be highly context-dependent. Positive effects in the IDENT-Freiburg experiment align with the pathway where functional diversity increasingly buffers drought stress during consecutive drought years. However, only species with comparably high-HSM_TLP_ were analyzed within this experiment. In a previous study at this experiment assessing only the initial drought year considered in our study, Hajek et al. (2022) found that species with higher HSM suffered less and tended to decrease the survival probability of their neighbors with low HSM, and vice versa. This suggests that thepositive effects observed here may result from selection effects leading to a competitive release of drought-tolerant species which profit from mortality suffered by drought-sensitive species (Grossiord 2020; Forrester and Bauhus 2016). Segregation of tree communities into ‘winner’ and ‘loser’ species in terms of diversity was also found in another tree diversity experiment under the same 2018-2020 drought (Sachsenmaier et al. 2024). As drought-sensitive species decline under intensifying water competition, positive diversity effects may increase for drought-tolerant species under prolonged drought stress (Maestre et al. 2009).

Conversely, negative diversity effects on growth observed in three sites support the pathway that prolonged drought stress can intensify with increasing functional diversity. Increased stress may result from greater water consumption in mixed-species stands compared to monocultures due to complementary resource-use strategies (Haberstroh and Werner 2022), early overyielding of tree biomass under favorable conditions (Jump et al. 2017; Jacobs et al. 2021), or selection of species with high water demands (Grossiord 2020; Forrester and Bauhus 2016). In the FORBIO-Zedelgem experiment, complementarity and selection effects were reported to drive early growth overyielding, enhancing canopy packing and stand transpiration (van de Peer et al. 2018; Wang et al. 2024). This positive diversity effects under favorable conditions could have negative impacts under increasing drought stress, when microclimate offsets from tree mixing might be insufficient to counteract drought stress (Zhang et al. 2022). In the Satakunta experiment, tree diversity did not enhance aboveground productivity or transpiration under non-limiting water availability (Grossiord et al. 2013). However, boreal forests with similar tree diversity showed higher water use efficiency and stomatal regulation than monocultures under drier conditions, indicating that species interactions can reduce soil moisture (Grossiord et al. 2014a). While complementarity in stomatal regulation, below-ground complementarity and water redistribution can mitigate drought impact to some extent (Bello et al. 2019; Moreno et al. 2024), these mechanisms may be overridden under severe or prolonged droughts (Grossiord et al. 2019; Mas et al. 2024b).

### Hydraulic trait identity determines growth responses to drought

HSM_TLP_ of focal species did not influence how tree diversity affected the growth responses during either a single-year drought or consecutive droughts (Q3). Species with contrasting HSM_TLP_ showed similar responses to increased tree diversity (Figure 3), differing from studies suggesting that drought-vulnerable species benefit more from higher species richness (Fichtner et al. 2020; Schnabel et al. 2024a; Sachsenmaier et al. 2024). However, species with high HSM_TLP_ maintained stable growth during a single-year drought, while lower-HSM_TLP_ species showed significant reductions independently of tree diversity. These differences were amplified under extreme drought, which suggests that xylem integrity under severe drought allow enhanced functioning and biomass growth (Choat et al. 2012; Martin-StPaul et al. 2017; Sanchez-Martinez et al. 2023; Anderegg et al. 2016). Notably, the significant interaction between drought intensity and HSM_TLP_ was evident only when using REW as a measure of water availability, emphasizing the importance of quantifying local soil water conditions for assessing tree drought stress over standardized climatic indices like SPEI (Zang et al. 2020; Schwarz et al. 2020).

Across consecutive drought years (Figure 6), most sites showed improved relative growth in the post-drought year for species with high HSM_TLP_, suggesting lower legacy effects for these drought-tolerant species (Anderegg et al. 2015). In contrast, at BIOTREE-Kaltenborn, drought-intolerant species had lower legacy effects compared to the species with higher HSM_TLP_, which showed higher growth reductions during consecutive drought and post-drought years. However, these contrasting HSM_TLP_ effects should be interpreted cautiously, given the different HSM_TLP_ ranges between sites and limited number of species analyzed per site. Additionally, species-specific mean trait values were collected from published databases, lacking site-specific data on trait variability. Intraspecific trait variability at the individual tree level can exceed interspecific differences (Anderegg 2015; Pritzkow et al. 2020). Trait plasticity can be driven by interspecific interactions even in early tree development stages, in turn influencing stand productivity and drought responses (Serrano-León et al. 2022; Benavides et al. 2019; Gazol et al. 2023). To date, the only study on tree diversity effects on intra-species HSM_TLP_ variability found that HSM_TLP_ in a subset of the experiments studied here is primarily driven by species identity and not by tree diversity, though some significant diversity effects were found for a limited number of species composition (Decarsin et al. 2024). The discrepancy in HSM_TLP_ effect across our sites aligns with findings that hydraulic traits do not always explain the magnitude of growth declines under intense, prolonged droughts (Song et al. 2022b; Smith-Martin et al. 2023).

### Applicability of results

A common limitation in dendrochronological studies like ours is the sampling bias toward canopy-dominant trees, aimed at reducing canopy shading effects on growth responses (Kannenberg et al. 2020; Duchesne et al. 2019). Although recent research shows that growth responses to short-term climate variability and drought legacy effects may not differ largely depending on sampling approach (Nehrbass-Ahles et al. 2014; Kannenberg et al. 2019b), drought responses can differ between dominant and suppressed trees. Larger trees can be less sensitive to competition for water and exhibit adaptations like greater water uptake, storage capacity, and more efficient water use and transport (Fernández-de-Uña et al. 2023; Colangelo et al. 2017). Our findings show that relative tree size within species has a positive effect on growth responses, while neighborhood competition had no significant impact. This aligns with reports that drought responses are more influenced by tree size, species identity and drought characteristics than by competition (Gillerot et al. 2021; Castagneri et al. 2022; Del Campo et al. 2022). We did not observe that tree size nor neighborhood competition affected the interaction between tree diversity and hydraulic traits, but it should be noted that our sampling strategy focused on dominant and co-dominant trees may not have captured a sufficiently wide range in tree sizes to elucidate such an effect. No comprehensive study has yet analyzed these factors across a tree diversity gradient including small and mid-sized trees of all species across varying levels of competition. Hence, the influence of these factors on tree diversity effects under drought remains unclear.

By using planted experiments with manipulated tree species richness, we minimized confounding effects of environmental heterogeneity within each site, which often obscure diversity effects in observational studies (Scherer-Lorenzen et al. 2007; Bauhus et al. 2017; Pardos et al. 2021). However, observed effects in young tree diversity experiments may differ from those in mature semi-natural forests, where drought responses can be influenced by tree age distributions and the development of interspecific interactions with progressing succession (Kambach et al. 2019; Leuschner et al. 2009). Hence, our findings are particularly relevant for plantation forestry and young, intensively managed forests (Messier et al. 2022; Depauw et al. 2024; Camarero et al. 2021). As species interactions are expected to strengthen over time (Guerrero-Ramírez et al. 2017; Jucker et al. 2017), continued monitoring of these plantations will help bridge the gap between observational studies and experimental findings.

Our findings underscore the importance of examining tree responses over extended periods to understand the effects of diversity on tree responses to drought. However, since we only assessed one post-drought year, we cannot draw robust conclusions about the recovery and resilience after these drought years, as drought legacy effects can persist for several years. Nonetheless, our results suggest that conclusions based on single-year drought responses may not hold under accelerated global warming, where prolonged drought events are increasingly followed by only brief recovery periods. As climate change leads to more frequent droughts, acclimation to chronic drought stress may become more crucial than recovery from isolated events (Boeck et al. 2017). However, the underlying processes remain unclear. Slow recovery might reflect controlled acclimation to optimize long-term survival (Gessler et al. 2020) or a gradual growth decline inducing tree mortality (DeSoto et al. 2020; Cailleret et al. 2017). Understanding the long-term impacts of consecutive droughts on forest ecosystems and the role of tree diversity to mitigate chronic stress is crucial as these events become more frequent.

## Conclusion

Our study is the first to examine the role of tree diversity and functional trait identity on tree growth responses across multiple experimental sites during unprecedented, multi-year droughts. We demonstrated that tree diversity effects on growth responses can intensify over consecutive drought years, with contrasting diversity effects varying from neutral to positive or even negative effects depending on the site. The context dependency of these diversity effects underlines the importance of considering multiple sites and neighborhood-scale processes in understanding how interspecific interactions shape tree growth responses under prolonged drought. Hydraulic traits played a significant role in determining drought-induced growth responses, emphasizing the need for trait-based approaches to assess drought impacts on forest ecosystems. Ultimately, integrating process-based models and hydraulic traits could provide forest managers with evidence-based guidelines to design more resilient, mixed-species plantations. By selecting species mixtures that are better adapted to local conditions and incorporating knowledge of hydraulic traits, drought-resilient mixed plantations can be used to enhance ecosystem resilience in the face of unprecedented droughts.

## Acknowledgments

We thank Kris Ceunen, Iris Moeneclaey, Bingbin Wen and Eline Lorer for assisting the fieldwork campaign; Toon Gheyle, Stijn Willen Ivan Josipovic for supporting the sample preparation and for scanning the samples; Leon Schmidt and Philip Nunner for assisting sample measurement and field data digitalization; Zoé Doucet and Wolf Wildpret for preparing the IDENT-Freiburg growth inventory dataset; Céline Meredieu for coordinating the management of the ORPHEE experiment; and Bart Muys for coordinating the management of the FORBIO Hechtel-Eksel experiment.

## Data availability

Data will be archived in the FunDivEUROPE data portal (https://fundiv.befdata.biow.uni-leipzig.de) upon publication and available upon request under Creative Commons licence (CC-BY).

