## Supplemental for "Multi-year drought strengthens positive and negative functional diversity effects on tree growth response"

#### Study experiments

Table S1. Details of experiment sites information and planting design. Biogeographical region is based on Cervellini et al. (2020). Climate corresponds with Köppen-Geiger classification (Kottek et al. 2006): Cfb = warm temperate fully humid warm summer, Csa = warm temperate summer dry hot summer, Dfc = snow fully humid cool summer. Species richness levels considered in the analysis are marked in bold. A list of the mixtures compositions considered in the analysis is detailed in Table S2.

| **Experiment**<br><br>(experiment subsites) | **B-Tree** | **BIOTREE-Kaltenborn** | **FORBIO-Gedinne**<br><br>(Gribelle & Gouverneurs) | **FORBIO-Hechtel-Eksel** | **FORBIO-Zedelgem** | **IDENT-Freiburg** | **IDENT-Macomer** | **ORPHEE** | **Satakunta**<br><br>(Area 1 & Area 3) |
|---|---|---|---|---|---|---|---|---|---|
| **Site code** | **B** | **K** | **F_G** | **F_H** | **F_Z** | **I_F** | **I_M** | **O** | **S** |
| **Country** | Austria | Germany | Belgium | Belgium | Belgium | Germany | Italy | France | Finland |
| **Latitude** (°) | 48.317639 N | 50.778835 N | 49.995833 N & 49.97819 N | 51.165 N | 51.148141 N | 48.019643 N | 44.739167 N | 61.715611 N & 61.67911 N | 61.715611 & 61.679111 |
| **Longitude** (°) | 16.066833 E | 10.221128 E | 4.976667 E & 4.983583 E | 5.311667 E | 3.119223 E | 7.82639 E | -0.798333 W | 21.978917 E & 21.699972 E | 21.978917 & 21.699972 |
| **Altitude** (m.a.s.l) | 200 | 325 | 370-420 | 55-56 | 11-16 | 240 | 615 | 60 | 20-60 |
| **Biogeographical region** | continental | continental | continental | Atlantic | Atlantic | continental | Mediterranean | Atlantic | boreal |
| **Climate** | Cfb | Cfb | Cfb | Cfb | Cfb | Cfb | Csa | Cfb | Dfc |
| **MAT** (°C) | 9.7 | 7.8 | 8.3 | 10.2 | 10.9 | 10.4 | 13.75 | 13.8 | 5.4 |
| **MAP** (mm) | 624 | 650 | 1336 | 887 | 850 | 887 | 915 | 944 | 550 |
| **Soil** | moist chernozem | orthoeutric arenosol | dry stony loam soils | dry sandy soil with gravel substrate | dry sandy soil to wet loamy sand soil | cambisol, loam | silt loam | sandy podzol on sandstone bedrock | podzol clay to podzol organic |
| **Plot size** ($m^2$) | 170-300 | 1920 | 1575-1764 | 1296 | 1764 | 13 | 10 | 400 | 400 |
| **Species richness levels** | **1, 2, 4** | **1, 2,** (3), **4** | **1, 2,** (3), **4** | **1, 2,** (3), **4** | **1, 2,** (3), **4** | **1, 2**, (4), **6** | **1, 2, 4**, (6) | **1, 2, 3, 4**, (5) | **1, 2**, (3), **5** |
| **Planting year** | 2013 | 2003 | 2010 | 2012 | 2009, 2010 | 2013 | 2014 | 2008 | 1999 |

| | | | | | | | | | |
|---|---|---|---|---|---|---|---|---|---|
| (seedling age at planting) | (2) | (3-4) | (1-3) | (2-3) | (2-4) | (2-3) | (1-2) | (1) | (1-2) |
| **Distance between rows** (m) | 1 | * | 1.5 | 1.5 | 1.5 | 0.45 | 0.4 | 2 | 1.5 |
| **Planting scheme** (tree-to-tree) | hexagonal | * | square | square | square | square | square | square | square |
| **Tree mixing** | stem-wise randomisation | patch-wise (monospecific patches 8m x 8m) | patch-wise (monospecific patches 4.5m x 4.5m) | patch-wise (monospecific patches 4.5m x 4.5m) | patch-wise (monospecific patches 4.5m x 4.5m) | stem-wise randomisation | stem-wise randomisation | stem-wise regular alternate pattern | stem-wise randomisation |

** Note on BIOTREE-Kaltenborn: Planting distance within planting rows: 2 m; distance between rows: 2m (gymnosperms) or 1m (angiosperms). Planting scheme (tree-to-tree): square (gymnosperms), rectangular (angiosperms).*

Table S2. Study species and mixtures compositions for each species considered at each site. Species in bold correspond with focal trees analyzed for drought responses.

| Site code | Species | Sp codes | Mixture intermediate-SR | Mixture high-SR | N cored trees | N analyzed samples |
|---|---|---|---|---|---|---|
| **B** | *Acer platanoides L.* | ACPL | ACPL-TICO | ACPL-CABE-QURO-TICO | 30 | |
| | *Carpinus betulus L.* | CABE | CABE-QURO | ACPL-CABE-QURO-TICO | 30 | |
| | ***Quercus robur** L.* | **QURO** | CABE-QURO | ACPL-CABE-QURO-TICO | 30 | **27** |
| | *Tilia cordata Mill.* | TICO | ACPL-TICO | ACPL-CABE-QURO-TICO | 30 | |
| **F_G** | *Acer pseudoplatanus L.* | ACPS | ACPS-PSME | ACPS-FASY-LAEU-QUPE, ACPS-FASY-PSME-QUPE | 35 | |
| | *Fagus sylvatica L.* | FASY | FASY-QUPE | ACPS-FASY-LAEU-QUPE, ACPS-FASY-PSME-QUPE | 39 | |
| | ***Larix × eurolepis** Henry* | **LAEU** | LAEU-PSME, LAEU-QUPE | ACPS-FASY-LAEU-QUPE | 40 | **37** |
| | ***Pseudotsuga menziesii** (Mirb.) Franco* | **PSME** | ACPS-PSME, LAEU-PSME | ACPS-FASY-PSME-QUPE | 40 | **29** |
| | ***Quercus petraea** (Matt.) Liebl.* | **QUPE** | FASY-QUPE, LAEU-QUPE | ACPS-FASY-LAEU-QUPE, ACPS-FASY-PSME-QUPE | 50 | **46** |
| **F_H** | *Betula pendula Roth* | BEPE | BEPE-LAKA | BEPE-LAKA-PISY-QUPE, BEPE-PISY-PSME-QUPE | 40 | |
| | ***Larix kaempferi** (Lamb.) Carrière* | **LAKA** | BEPE-LAKA, LAKA-PSME | BEPE-LAKA-PISY-QUPE | 40 | **35** |
| | ***Pinus sylvestris** L.* | **PISY** | PISY-PSME, PISY-QUPE | BEPE-LAKA-PISY-QUPE, BEPE-PISY-PSME-QUPE | 50 | **47** |
| | ***Pseudotsuga menziesii** (Mirb.) Franco* | **PSME** | LAKA-PSME, PISY-PSME | BEPE-PISY-PSME-QUPE | 40 | **35** |
| | ***Quercus petraea** (Matt.) Liebl.* | **QUPE** | PISY-QUPE | BEPE-LAKA-PISY-QUPE, BEPE-PISY-PSME-QUPE | 40 | **38** |
| **F_Z** | ***Betula pendula** Roth* | **BEPE** | BEPE-PISY, BEPE-QURO | BEPE-FASY-PISY-TICO | 40 | **35** |
| | ***Fagus sylvatica** L.* | **FASY** | FASY-QURO | BEPE-FASY-PISY-TICO, FASY-PISY-QURO-TICO | 40 | **31** |
| | ***Pinus sylvestris** L.* | **PISY** | BEPE-PISY, PISY-TICO | BEPE-FASY-PISY-TICO, FASY-PISY-QURO-TICO | 50 | **45** |
| | ***Quercus robur** L.* | **QURO** | BEPE-QURO, FASY-QURO | FASY-PISY-QURO-TICO | 40 | **38** |
| | ***Tilia cordata** Mill.* | **TICO** | PISY-TICO | BEPE-FASY-PISY-TICO, FASY-PISY-QURO-TICO | 40 | **38** |
| **K** | ***Fagus sylvatica** L.* | **FASY** | FASY-PIAB, FASY-PSME, FASY-QUPE | FASY-PIAB-PSME-QUPE | 50 | **47** |
| | ***Picea abies** (L.) H. Karst.* | **PIAB** | FASY-PIAB, PIAB-PSME, PIAB-QUPE | FASY-PIAB-PSME-QUPE | 50 | **43** |
| | ***Pseudotsuga menziesii** (Mirb.) Franco* | **PSME** | FASY-PSME, PIAB-PSME, PSME-QUPE | FASY-PIAB-PSME-QUPE | 50 | **44** |
| | ***Quercus petraea** (Matt.) Liebl.* | **QUPE** | FASY-QUPE, PIAB-QUPE, PSME-QUPE | FASY-PIAB-PSME-QUPE | 50 | **47** |

Table S2 (cont.) Study species and species-compositions of mixtures considered in each site. Species in bold correspond with focal trees analyzed for drought responses.

| Site code | Species | Sp codes | Mixture intermediate-SR | Mixture high-SR | N cored trees | N analyzed samples |
|---|---|---|---|---|---|---|
| **I_F** | ***Acer platanoides*** *L.* | **ACPL** | ACPL-QURU | ACPL-ACSA-BEPA-BEPE-QURO-QURU | 33 | **25** |
| | ***Acer saccharum*** *Marshall* | **ACSA** | ACSA-QURO | ACPL-ACSA-BEPA-BEPE-QURO-QURU | 32 | **29** |
| | *Betula papyrifera Marshall* | BEPA | | ACPL-ACSA-BEPA-BEPE-QURO-QURU | 13 | |
| | *Betula pendula Roth* | BEPE | | ACPL-ACSA-BEPA-BEPE-QURO-QURU | 17 | |
| | ***Quercus robur*** *L.* | **QURO** | ACSA-QURO | ACPL-ACSA-BEPA-BEPE-QURO-QURU | 27 | **21** |
| | ***Quercus rubra*** *L.* | **QURU** | ACPL-QURU | ACPL-ACSA-BEPA-BEPE-QURO-QURU | 33 | **27** |
| **I_M** | *Arbutus unedo L.* | ARUN | ARUN-ACMO | | 18 | |
| | ***Pinus pinea*** *L.* | **PIPI** | PIPI-FROR | ACMO-FROR-PIPI-QUPU | 27 | **27** |
| | *Acer monspessulanum L.* | ACMO | | | | |
| | *Fraxinus ornus L.* | FROR | | | | |
| | *Quercus pubescens Willd.* | QUPU | | | | |
| **O** | *Betula pendula Roth* | BEPE | BEPE-PIPT, BEPE-QUIL | BEPE-PIPT-QUIL, BEPE-PIPT-QUIL-QUPY, BEPE-PIPT-QUIL-QURO | 60 | |
| | ***Pinus pinaster*** *Aiton* | **PIPT** | BEPE-PIPT, PIPT-QUIL | BEPE-PIPT-QUIL, BEPE-PIPT-QUIL-QUPY, BEPE-PIPT-QUIL-QURO | 60 | **59** |
| | *Quercus ilex L.* | QUIL | | | | |
| | *Quercus pyrenaica Willd.* | QUPY | | | | |
| **S** | *Alnus glutinosa (L.) Gaertn.* | ALGL | ALGL-BEPE | ALGL-BEPE-LASI-PIAB-PISY | 30 | |
| | *Betula pendula Roth* | BEPE | ALGL-BEPE, BEPE-PISY | ALGL-BEPE-LASI-PIAB-PISY | 40 | |
| | ***Larix sibirica*** *Ledeb.* | **LASI** | LASI-PIAB | ALGL-BEPE-LASI-PIAB-PISY | 29 | **24** |
| | ***Picea abies*** *(L.) H. Karst.* | **PIAB** | LASI-PIAB, PIAB-PISY | ALGL-BEPE-LASI-PIAB-PISY | 40 | **40** |
| | ***Pinus sylvestris*** *L.* | **PISY** | BEPE-PISY, PIAB-PISY | ALGL-BEPE-LASI-PIAB-PISY | 40 | **37** |

### Drought selection

#### Supplementary Method 1. Calculation of drought indices

Droughts were identified as periods of extreme deficit in water availability following a climate-based approach with the use of site-specific data, as opposed to growth-based indices (Schwarz et al. 2020; Slette et al. 2019). We obtained climate data for the period 1991-2021 from the ERA5Land of the Copernicus Climate Data Store (https://cds.climate.copernicus.eu; (Muñoz-Sabater et al. 2021), which includes monthly averaged data at 0.1° x 0.1° grid-cell resolution (9 km horizontal resolution) on total precipitation (P), total evaporation (ET), potential evaporation (PET), and air temperature (Temp) measured at a 2-m height (Figure S1). We identified recent drought years and pre-drought reference years by comparing different drought indices. First, we calculated the Standardized Precipitation Evapotranspiration Index (SPEI) (Slette et al. 2019) using the R-package SPEI (Vicente-Serrano et al. 2010; Beguería and Vicente-Serrano 2017). SPEI is calculated from the local monthly climatic water balance (precipitation minus potential evapotranspiration) over a selected moving time window, with its deviation from the long-term mean expressed as a standardized Gaussian variable with a mean of zero and a standard deviation of one (Beguería et al. 2010; Beguería et al. 2014; Vicente-Serrano et al. 2010). Based on the monthly meteorological data, we calculated the monthly SPEI for different window periods corresponding to annual SPEI12 December, and growing season as SPEI6 April-September. We used the SPEI calculated over the reference 30-year period (1991-2021) to determine abnormally dry growing seasons for each experiment location using the threshold -1.28 as the 10% quantile of all values following the drought classification of the SPEI global drought monitor (Agnew 2000; Beguería et al. 2022).

In addition to the climatic indices, we accounted for the importance of local soil characteristics on the capacity to hold plant available water, which can either amplify or dampen tree growth reactions to climatic droughts (Schwarz et al. 2020). We characterized the drought intensity in terms of plant drought stress experienced by the trees calculating the relative extractable water (REW) in the soil of each site using the generic process-based model SurEau (Ruffault et al. 2022) and following the methodology used in Blondeel et al. (2024). SurEau assesses forest stand water balance and extreme drought stress impacts on vegetation based on plant hydraulic theory (Ruffault et al. 2013). The overall REW is computed from soil water content as follows:

$$REW = \frac{\theta - \theta_{\wp}}{\theta_{fc} - \theta_{\wp}}$$

where $\theta$ is the actual soil water content, $\theta_{fc}$ content at field capacity, and $\theta_{\wp}$ is the soil water content at the wilting point (i.e. at -1.5MPa). The model is driven by daily aggregated hourly climatic data, along with soil properties (such as moisture retention curves and depth), plant characteristics related to stomatal control, drought-induced cavitation resistance, and leaf area index (LAI). It calculates the water balance at the stand level and predicts tree water potential at an hourly resolution. The model was applied at each site using vegetation parameters specific to one species representative of the biome and the local species pool of each experiment as in Blondeel et al. (2024). Details of input variables for the SurEau simulations are available in Table S3. Trait data was sourced from global databases (see 'Functional traits selection'). Soil parameters were extracted from the SoilGrids database (Poggio et al. 2021). LAI input data for the SurEau model were obtained from the high-resolution (333m) Copernicus database (Baret et al. 2013; Lacaze et al. 2015). Climate data for each study site was obtained from ERA5-Land hourly data (Muñoz-Sabater et al. 2021). Minimum monthly REW during the growing season was calculated as a measure of drought intensity. REW value 1 corresponds with soil water at field capacity, 0 when soil water content reaches the wilting point. The drought stress threshold is considered at a REW value of 0.4 (Granier et al. 1999). Negative values at soil water content below the permanent wilting point can occur in SurEau due to delayed stomatal closure or residual transpiration.

Table S3. Input variables for the calculation of the SurEau water balance model. Simulations of plant drought stress were carried out for the site-specific soil conditions and based on one representative species for each experimental site. LAI_max refers to the maximum Leaf Area Index (LAI) used for the analysis. SoilClass corresponds with the soil classification from SoilGrids (C = Clay, L = Loam, S = Sand, Si = Silt). TAW_target is the total soil available water used in the model, computed for each site based on the texture information available from the corresponding SoilClass variable. The species-specific trait values used in the model simulations are detailed in Table S6.

| **Site** | **Longitude** | **Latitude** | **LAI_max** | **SoilClass** | **Species** | **TAW_target** |
|---|---|---|---|---|---|---|
| B-Tree | 16.06 | 48.33 | 2.1 | L | *Quercus robur* | 24.2 |
| BIOTREE_Kaltenborn | 10.22 | 50.78 | 6.1 | CL | *Fagus sylvatica* | 166.4 |
| FORBIO_Gedinne | 4.98 | 49.98 | 5.7 | SiL | *Fagus sylvatica* | 145 |
| FORBIO_Hechtel | 5.31 | 51.17 | 4.5 | SL | *Betula pendula* | 121.8 |
| FORBIO_Zedelgem | 3.12 | 51.15 | 6.1 | L | *Quercus robur* | 232.8 |
| IDENT_Freiburg | 7.83 | 48.02 | 5.2 | L | *Betula pendula* | 147.3 |
| IDENT_Macomer | 8.72 | 40.24 | 2.5 | CL | *Arbutus unedo* | 76.7 |
| ORPHEE | -0.77 | 44.67 | 4.5 | SCL | *Betula pendula* | 214.6 |
| Satakunta | 21.98 | 61.72 | 5 | L | *Betula pendula* | 71.3 |

Table S4. Drought periods, reference pre-drought years and post-drought years at the study sites considered for the analysis. For research question Q1, all sites were analyzed for single-year drought responses corresponding with the initial drought year (Resp_yr 1). For Q2, sites in bold experiencing the same multiyear drought period were analyzed for the growth responses during the same consecutive drought years (Resp_yr 1-3) and post-drought (Resp_yr 4).

| | | | | **Response years for analysis** | | | |
|---|---|---|---|---|---|---|---|
| | | | | **Resp_yr 1** | **Resp_yr 2** | **Resp_yr 3** | **Resp_yr 4** |
| **Site** | **Planting year** | **Pre-drought wet yr1** | **Pre-drought wet yr2** | **Drought yr1** | **Drought yr2** | **Drought yr3** | **Post-drought wet yr1** |
| B-Tree | 2013 | 2016 | | 2017 | 2018 | 2019 | 2020 |
| **BIOTREE_Kaltenborn** | 2004 | 2017 | | **2018** | **2019** | **2020** | **2021** |
| **FORBIO_Gedinne** | 2010 | 2016 | 2017 | **2018** | **2019** | **2020** | **2021** |
| **FORBIO_Hechtel-Eksel** | 2012 | 2016 | 2017 | **2018** | **2019** | **2020** | **2021** |
| **FORBIO_Zedelgem** | 2009 | 2016 | | **2018** | **2019** | **2020** | **2021** |
| **IDENT_Freiburg** | 2013 | 2016 | | **2018** | **2019** | **2020** | **2021** |
| IDENT_Macomer | 2014 | 2016 | | 2017 | | | 2018 |
| ORPHEE | 2008 | 2017 | 2018 | 2019 | | | 2020 |
| **Satakunta** | 2000 | 2016 | 2017 | **2018** | **2019** | **2020** | **2021** |

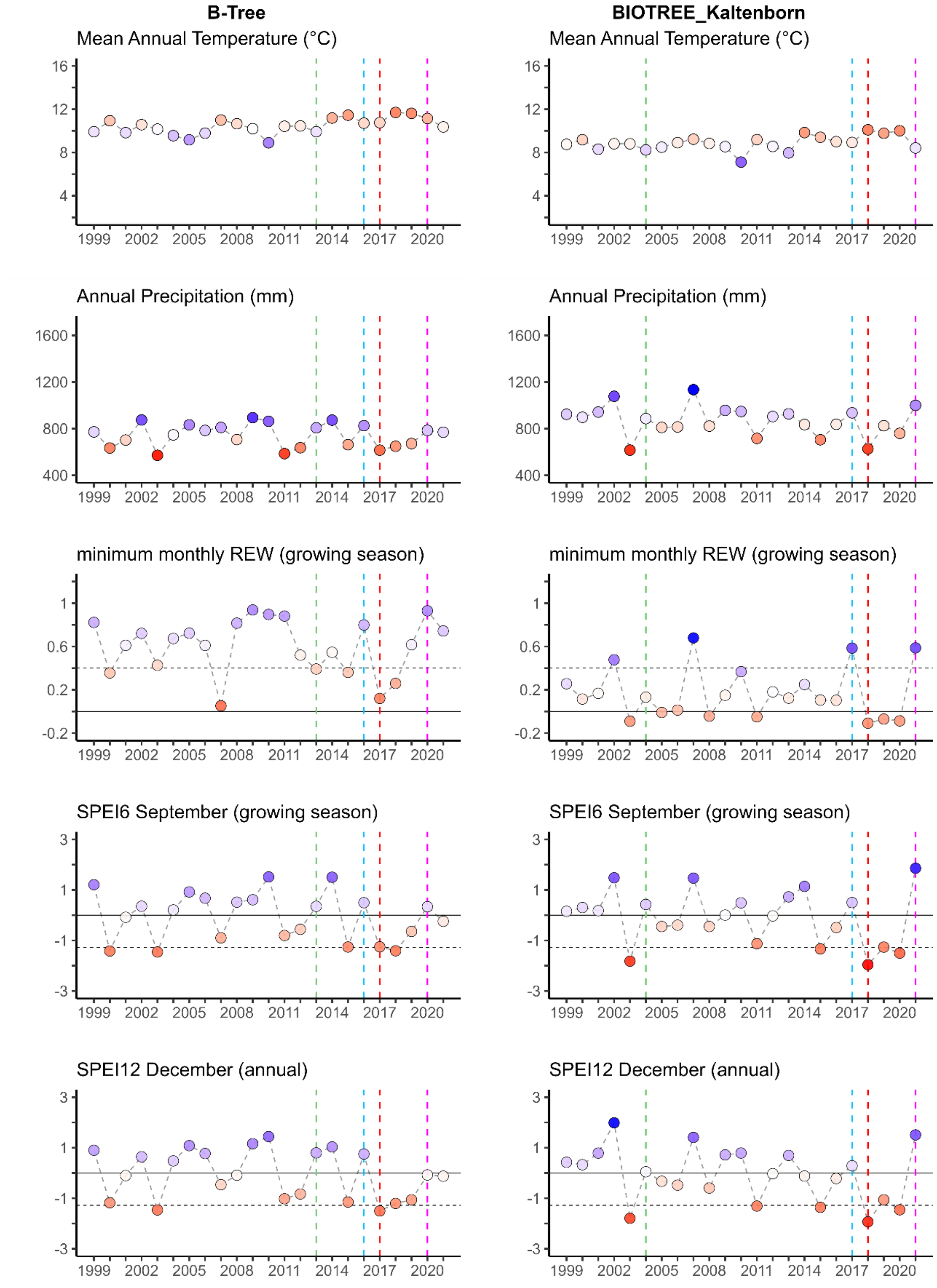


Figure S1. Long-term climate conditions at the study sites used for Q1 analysis of a single-year drought. Red points indicate drier/hotter conditions and blue points indicate wetter/colder conditions compared to the long-term mean for the period 1991-2021. SPEI = -1.28 indicates the threshold for selection of abnormally dry years. REW = 0.4 indicates the drought stress in terms of minimum monthly relative extractable water, and REW = 0 indicates soil water content reaching the wilting point. Vertical dashed lines represent the experiment planting year (green), the pre-drought reference year(s) (blue), the initial drought year (red), and the post-drought year with normal or wet conditions (violet). Figure continue in following pages for other sites.

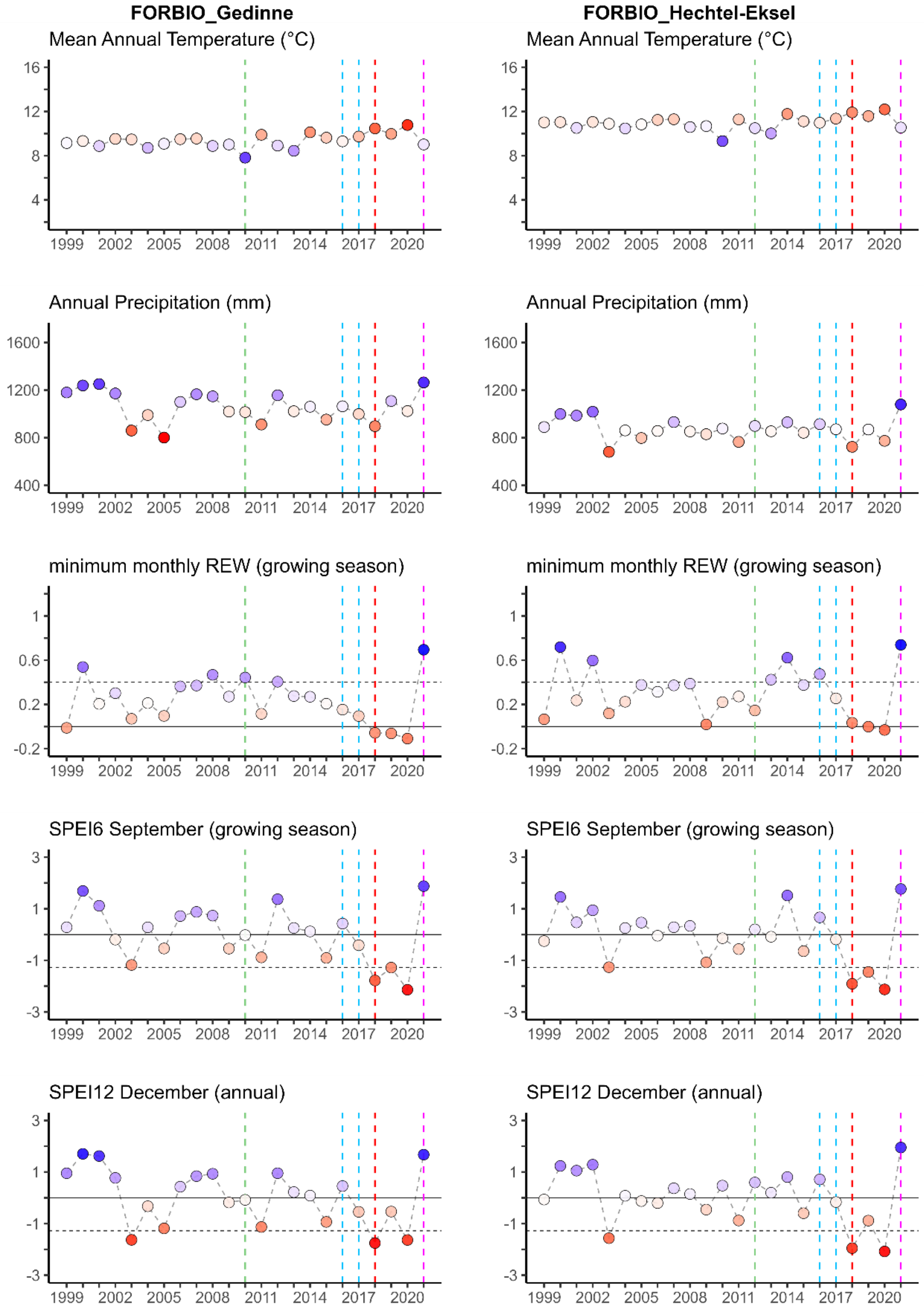


Figure S1 (cont.). Long-term climate conditions at the study sites.

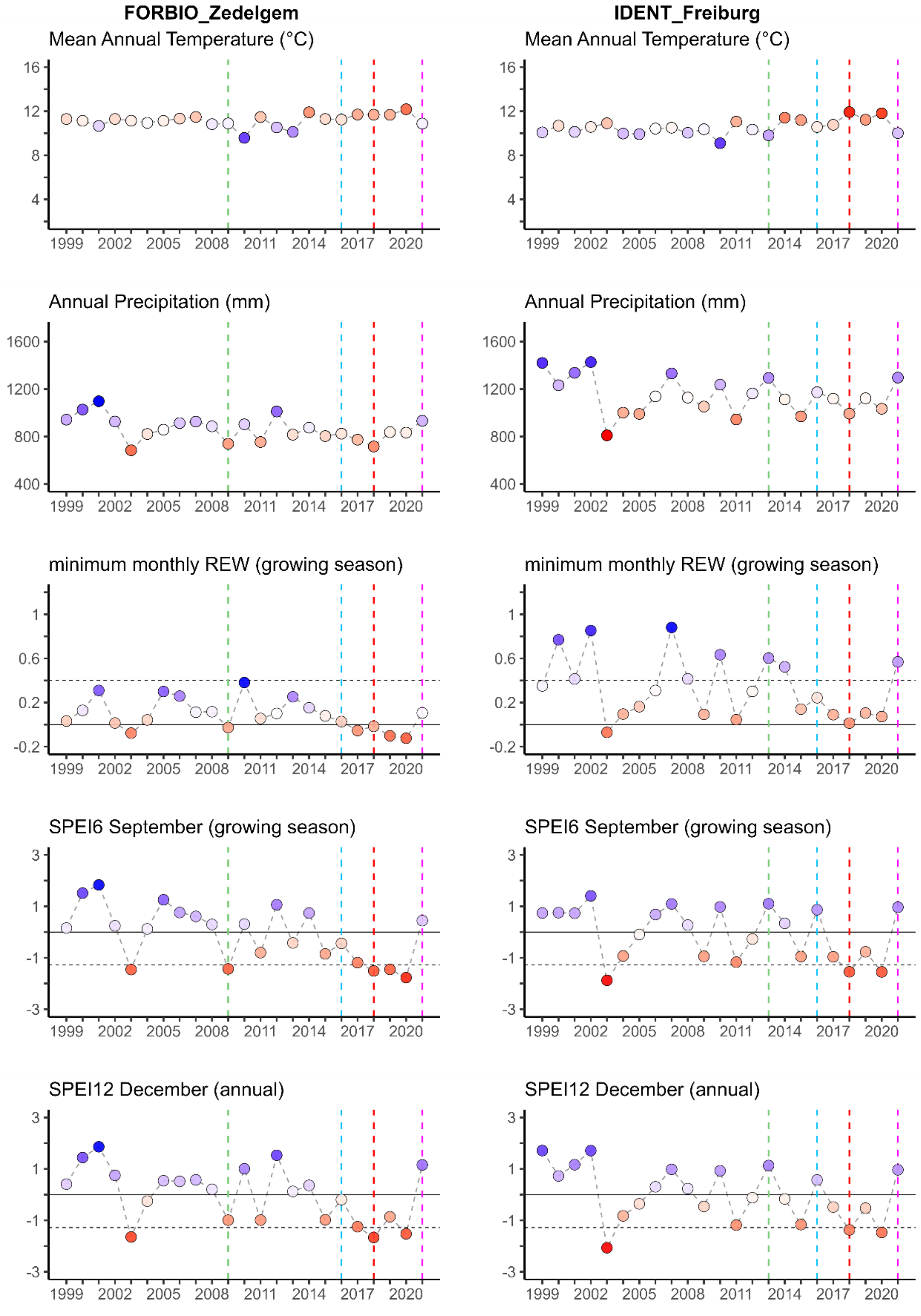


Figure S1 (cont.). Long-term climate conditions at the study sites.

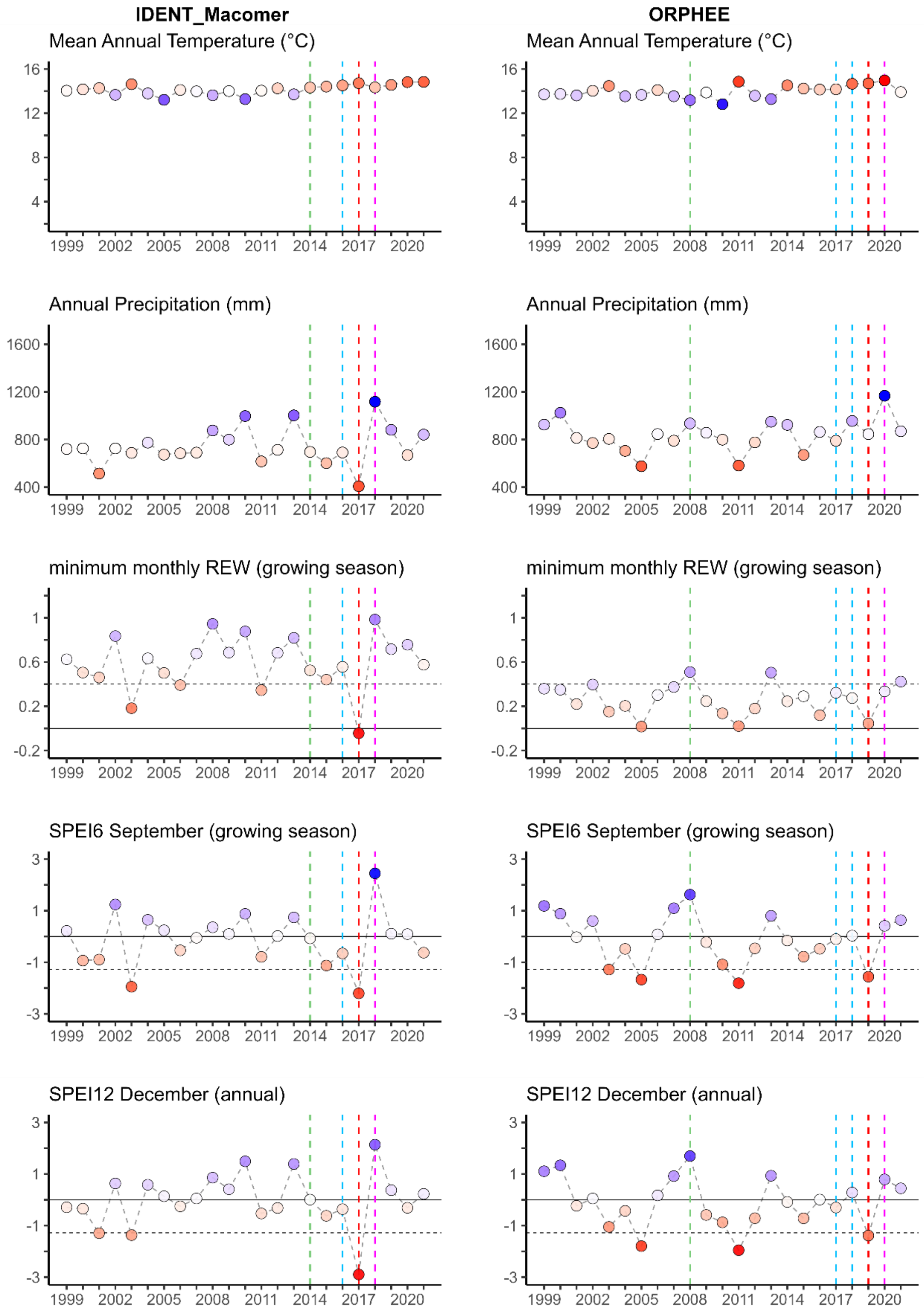


Figure S1 (cont.). Long-term climate conditions at the study sites.

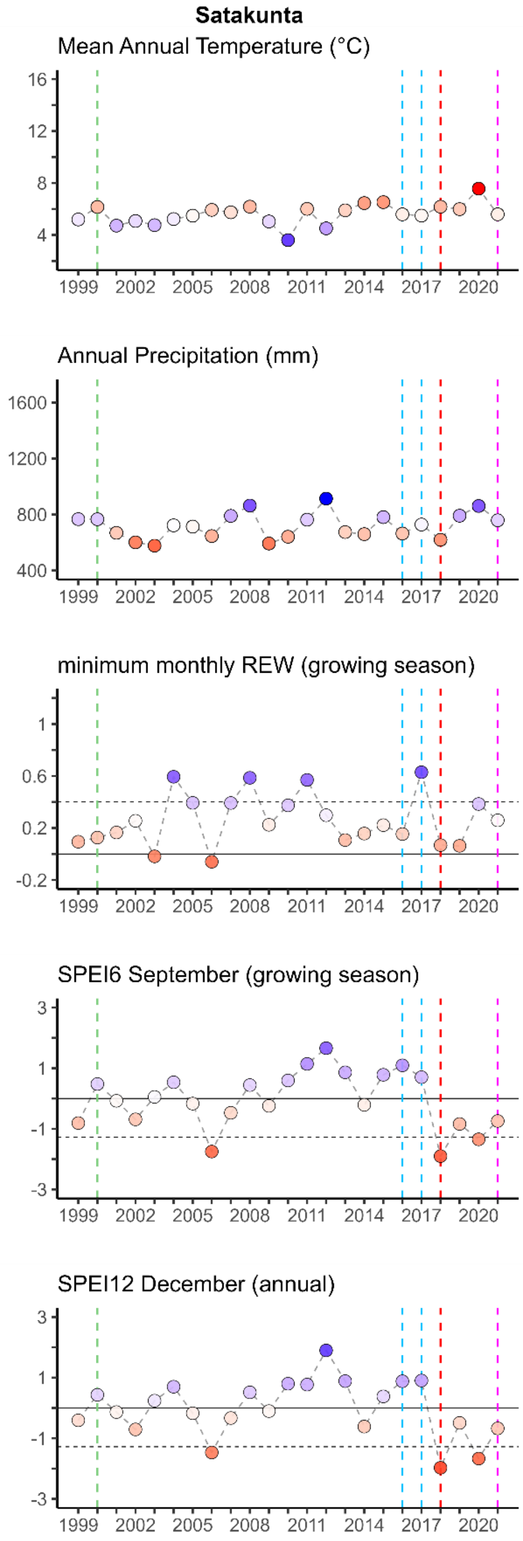


Figure S1 (cont.). Long-term climate conditions at the study sites.

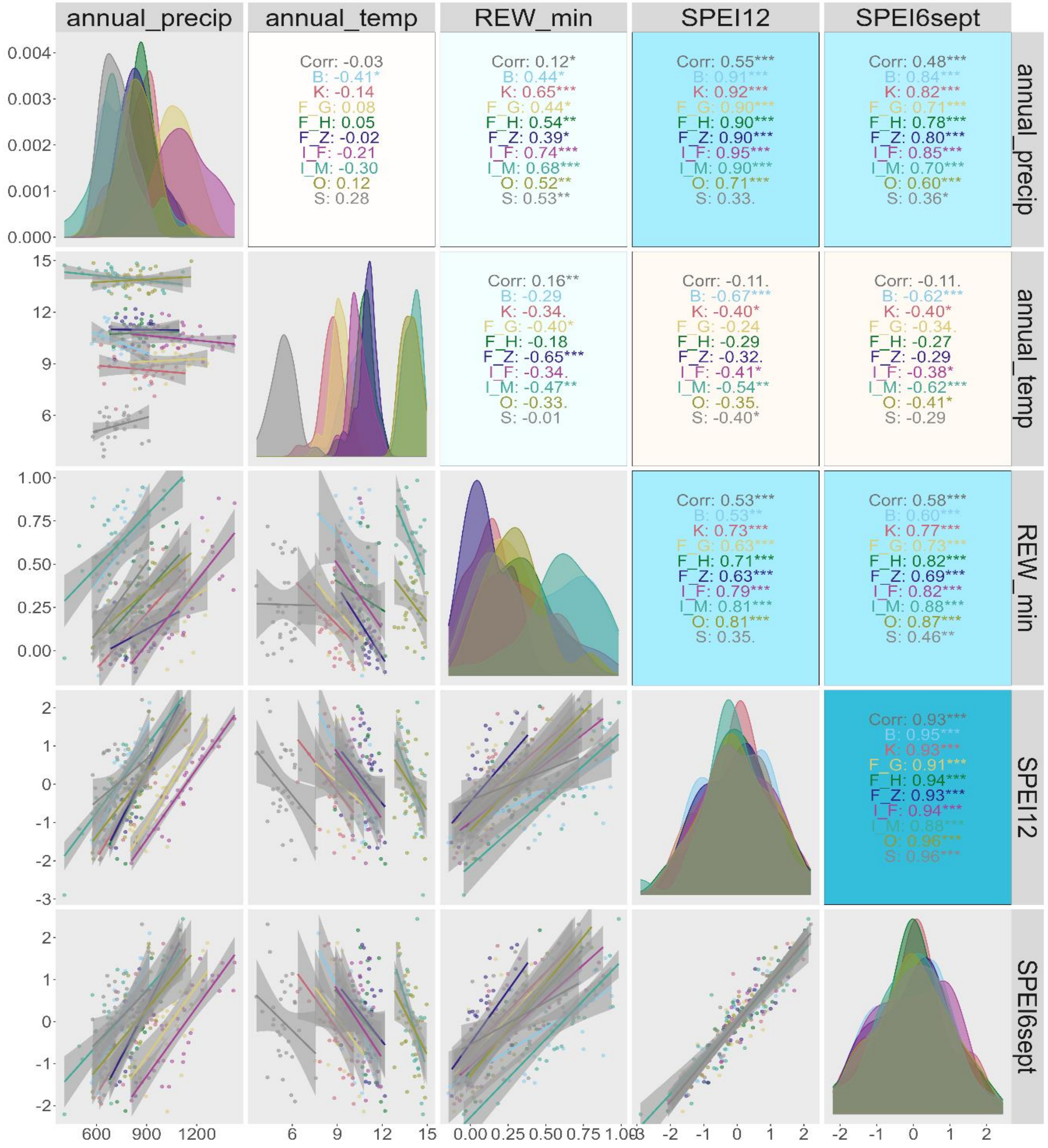


Figure S2. Pairwise correlations between different climatic variables for the period 1991-2021: annual precipitation (mm), annual temperature (°C), minimum monthly relative extractable water (REW min) during the growing season, Standardized Precipitation Evapotranspiration Index (SPEI) for the annual conditions (SPEI12 December) and the growing season (SPEI6 April-September). The different drought indices used for the selection of drought years (REW min, SPEI12 December, SPEI6 April-September) have comparable temporal patterns and high correlation within each site. Upper-right panels show the pairwise correlations, overall and within site, with significance levels referred as * p<0.05, ** p<0.01 and *** p<0.001 and background heatmap colors representing the correlation strength and direction (positive correlations in blue, negative in red). Diagonal panels show the density plots of the variables for each site. Lower panels show the bivariate linear regressions with 95% confidence level interval for each site. Pairwise correlations calculated using the function ggpairs() in the R packages GGally (Schloerke et al. 2024).

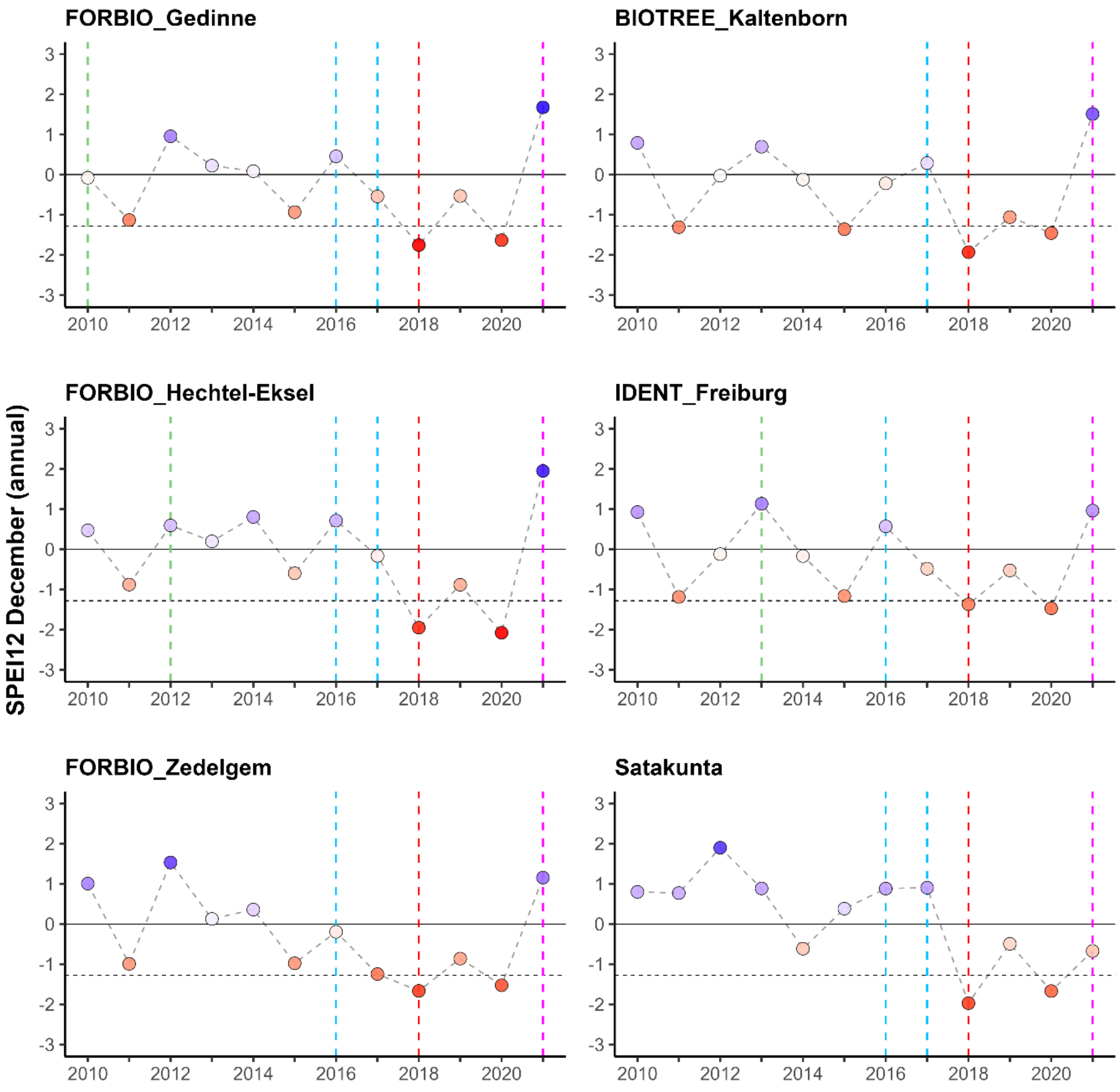


Figure S3. Drought years selected for the analysis of research question Q2 in sites experiencing a multiyear drought event. Drought selection based on the Standardized Precipitation Evapotranspiration Index (SPEI) for the annual conditions (SPEI12 December). Red points indicate drier/hotter conditions and blue points indicate wetter/colder conditions compared to the long-term mean. SPEI below -1.28 (horizontal dashed line) indicates the threshold for selection of abnormally dry years. Blue vertical dashed lines represent the pre-drought reference year(s) for growth responses. Red vertical dashed lines represent the initial drought year being severely dry (Resp_yr 1 with SPEI < -1.28) followed by two consecutive years with moderate drought conditions (Resp_yr 2-3 with SPEI ≈ -1). Violet lines represent the post-drought year with normal or wet conditions (Resp_yr 4 in analysis). Green vertical dashed lines represent the experiment planting year.

#### Neighborhood selection

Table S5. Neighborhood radius definition to focal trees at each study site. Neighborhood radius adapted to the site-specific planting pattern and planting distance in order to include all direct 1st-order neighbors and 2nd-order neighbors with crowns interacting with the focal tree's crown.

| | **B** | **F_G, F_H, F_Z, S** | **O** | **K** | **I_F** | **I_M** |
|---|---|---|---|---|---|---|
| **Planting pattern** | hexagonal | square | square | square/ rectangular | square | square |
| **$1^{st}$-order neighborhood:** | | | | | | |
| **min distance (m)** | 1 | 1.5 | 2 | 1 | 0.4 | 0.45 |
| **max distance (m)** | | 2.1 | 2.8 | 2.8 | 0.6 | 0.6 |
| **$2^{nd}$-order neighborhood:** | n.a. | n.a. | n.a. | n.a. | | |
| **min distance (m)** | 1.7 | 3 | 4 | 3 | 0.8 | 0.9 |
| **max distance (m)** | | | | | 1.1 | 1.3 |
| **$3^{rd}$-order neighborhood:** | n.a. | n.a. | n.a. | n.a. | | |
| **min distance (m)** | | | | | 1.2 | 1.4 |
| **max distance (m)** | | | | | 1.7 | 1.9 |
| **$4^{th}$-order neighborhood:** | n.a. | n.a. | n.a. | n.a. | n.a. | n.a. |
| **min distance (m)** | | | | | 1.6 | 1.8 |
| **max distance (m)** | | | | | 2.2 | 2.6 |
| **Neighborhood radius definition (m):** | | | | | | |
| **only $1^{st}$-order available data** | **< 1.5** | **< 2.9** | **< 2.9** | **< 2.9** | | |
| **including all $1^{st}$- and $2^{nd}$-order + closest $3^{rd}$-order** | | | | | **< 1.5** | **< 1.5** |

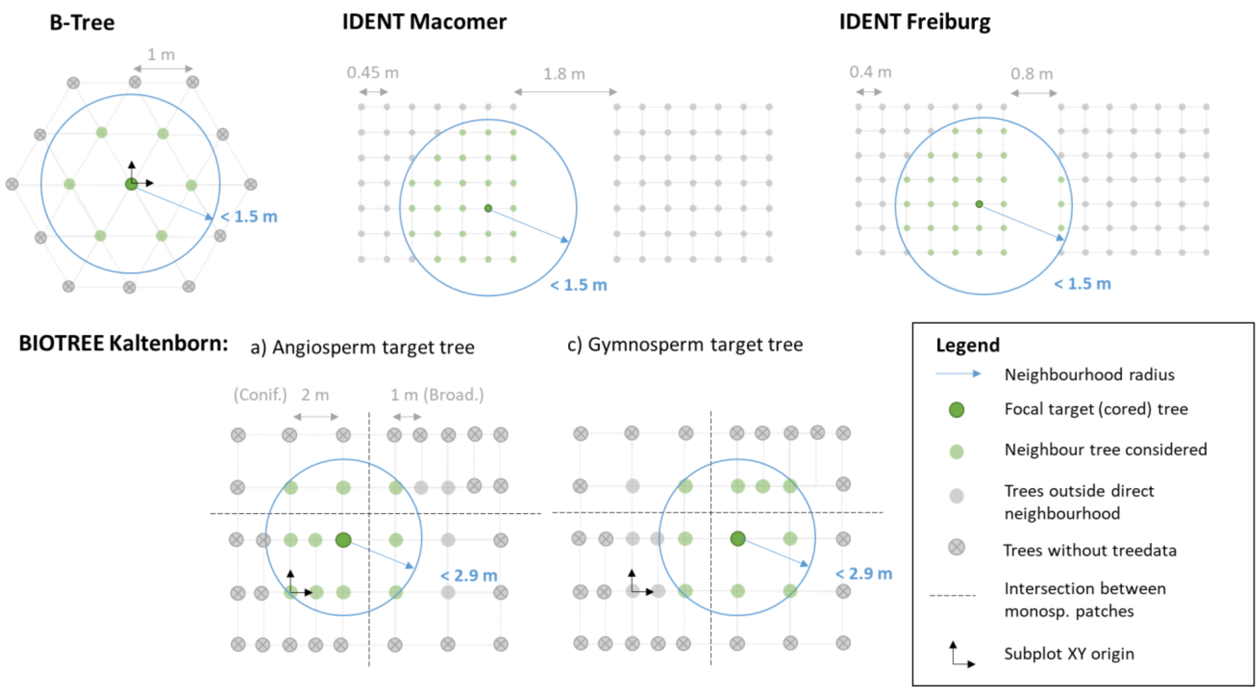


Figure S4. Neighborhood definition at sites that required adapted radius definition to the specific planting design and planting distance.

#### Tree ring measurements

Supplementary Method 2. Tree ring measurements. µCT scanning and tree ring series crossdating

We obtained annual series of tree-ring width (TRW) and mean wood density using X-ray micro-Computed Tomography (µCT) with the HECTOR µCT scanner (Masschaele et al. 2013) at the Ghent University Centre for X-ray tomography (UGCT; http://www.ugct.ugent.be). Before scanning, increment cores were dried at room temperature for one month to be in balance with the scanner room environment (20.05 ± 0.13 ºC, relative humidity 34.04 ± 1.62 %). We prepared and scanned increment cores following the workflow for µCT densitometry detailed in de de Mil et al. (2016). Cores were dried at room temperature for one month to be in balance with the scanner room environment (20.05 ± 0.13ºC, relative humidity 34.04 ± 1.62 %). Cores were mounted in customized cardboard sample holders and positioned on the rotation stage of the scanner in a dedicated master sample holder, which allowed batch scanning (de Mil et al. 2016). The samples were scanned at an energy of 100 kV and with a 1 mm aluminium filter, acquiring 2200 projections at an exposure time of 1 s per projection. Scanned three-dimensional (3D) images had an approximate voxel pitch (hereafter referred as resolution) of 50 µm. Reconstructions of the scanned sample batch were performed using the Octopus Reconstruction software (Vlassenbroeck et al. 2007). We extracted the 3D images of each individual core from the reconstructed volumes, adjusting tilt and tangential alignment of each core perpendicular to the grain direction, and converted to density estimates using using specific MATLAB-based densitometry toolboxes for editing dendrochronological µCT images (van den Bulcke et al. 2014; de Mil et al. 2016; de Mil and van den Bulcke 2023). The Standard Operating Procedure (SOP) for the Densitometry, DHXCT and CoreComparison toolbox is available in https://dendrochronomics.ugent.be/#software. Using the DensitometryToolbox, wood density was estimated by rescaling the full greyscale volume based on the known density of 3 reference slots of air and a reference material with similar elemental composition to wood (de Mil et al. 2016).

TRW and wood density profiles were measured using the DHXCT software toolbox for editing dendrochronological µCT images. Tree ring boundaries were manually indicated as well as fibre and tree-ring angles were also considered, and TRW was calculated by the toolbox considering these parameters. Tree ring outputs of the DHXCT toolbox were imported in the CoreComparison toolbox to visualize Gleichläufigkeit coefficient (glk) and correlation parameters of the tree ring series (de Mil et al. 2016), and to export the series of TRW and density values (mean/max/min/min per quartile/ max per quartile) of every tree ring per core sample.

Crossdating of tree ring series was performed following several steps: (1) visual crossdating of marker rings (narrow rings, rings with intra-annual density fluctuations IADF) based on the DHXCT images; (2) glk coefficient, correlation and pattern matching of tree ring series per species/site using the CoreComparison toolbox; and (3) crossdating with the DplR package in R (Bunn 2010; Bunn et al. 2023) using correlation between each tree-ring series and a master leave-one-out chronology (function corr.rwl.seg) and cross-correlation of a single series with a running correlation window and different lags using (function corr.series.seg). Samples included in the analysis underwent a rigorous quality filtering procedure to ensure reliability of tree ring definition, excluding those samples with ambiguous tree ring definitions after the crossdating processing. Additionally, to maintain statistical robustness and ensure representative data across species richness levels, species with a substantial proportion of doubtful samples (i.e. > 40%) at a given site were omitted from the analysis. A total of 948 trees from 16 species and 68 different species compositions (26 monocultures, 28 intermediate-richness mixtures, 14 high-richness mixtures) were included in the following analysis (Table S2, see also Figure S5 for an overview of the wood anatomy of the species).

We calculated basal area increment (BAI, $mm^2$ $year^{-1}$) series from pith to bark based on the TRW series using the bai.in() function in the dplR package in R (Bunn 2008; Bunn et al. 2023). We calculated BAI from pith to bark as opposed to the alternative routine bai.out() from bark to pith, which is based on the bark thickness and outer diameter measurements (Visser et al. 2023; Bunn 2008). We followed this approach since the bark thickness is highly variable depending on the coring point and largely shrink during the drying process, hence not being representative for the diameter measurements in the field. We estimated an indicator for radial biomass increment (hereafter BIOMinc, kg $m^{-1}$, $year^{-1}$) as the product of BAI and mean wood density (kg $m^{-3}$) of the ring. This indicator better reflects the actual carbon investment in radial growth than TRW and BAI metrics (Camarero and Andrés 2024) and it can be considered a proxy of aboveground biomass growth when continuous inventory data in annual tree height growth is missing (Vannoppen et al. 2018; Bontemps et al. 2010). Note that ring series of mean wood density (kg $m^{-3}$) were obtained from the DHXCT density profiles scanned after drying the cores at room air humidity. Although extrapolating findings to forest biomass for carbon sequestration estimations would require further conversion (i.e. to basic wood density after oven drying) (Vieilledent et al. 2018; Bondarenko et al. 2018), this approach allows us to analyse differences in biomass increment at individual tree-level across the experimental gradients.

In order to reduce the presence of age trends in growth series in the dynamic growth phase of the juvenile or recently released experiment trees (Schwarz et al. 2020), we tested individual series standardization with different detrending methods (modified Hugershoff curve, Cook and Peters smoothing spline, Friedman's super smoother, age-dependent spline) using the detrend() function in DplR (Bunn 2008; Bunn et al. 2023). However, given the short time length of the growth series, none of the methods allowed for the fitting of all individual series. Testing of different smoothness windows (nyrs parameter) of the fitted curves resulted in either fitting fails or spline overfitting in multiple samples. Measurements of the drought-induced responses showed a strong sensitivity to the detrending method, which would result on potential removal or overestimation of the drought effect depending on the method (results not shown). Alternatively, detrending using regional curve standardization (RCS, reference curve fixed for all tree series per site and species) assumes that trees representing a similar habitat with negligible variations in competition and age structure (Helama et al. 2017), which was not adequate for our study given the large variations in juvenile competition between plots and different individual growth patterns. Given the challenges to detrend age-effect in such sort time series and the strong influence of the detrending method, we analyzed drought-induced growth responses based on raw undetrended series of both TRW, BAI and BIOMinc and compared the three radial growth proxies (Schwarz et al. 2020; Schnabel et al. 2024). BIOMinc series are less influenced by biological age trends than TRW (Figure S8), being a more reliable indicator of temporal trends in radial growth for young, open-grown trees (Biondi and Qeadan 2008).

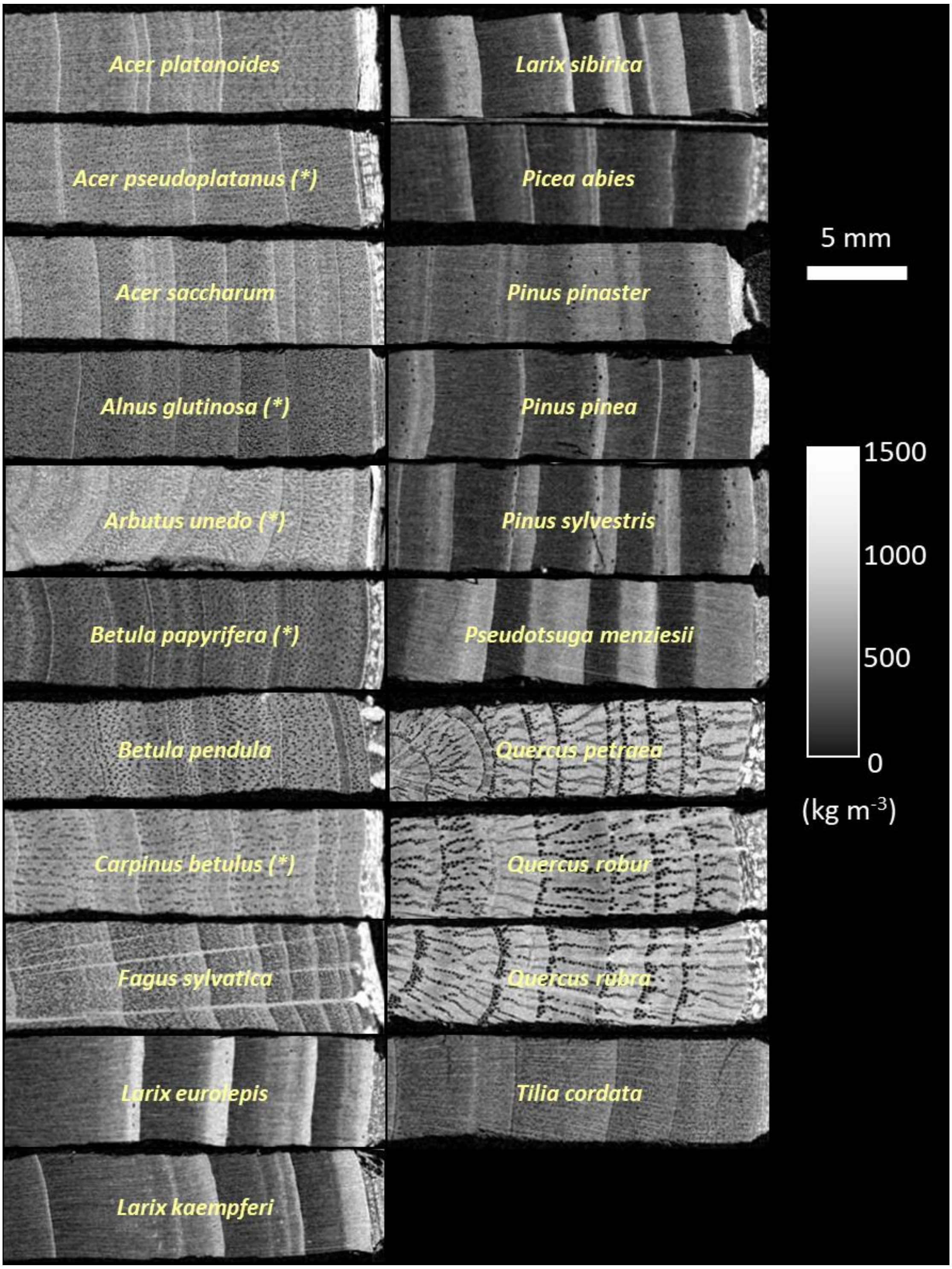


Figure S5. Wood anatomy of the tree species sampled in this study. Images are transversal sections of exemplary cores per species. Images were taken from scanned 3D volumes of the wood cores with X-ray micro-Computed Tomography (µCT) and densitometry toolboxes for editing dendrochronological helical XCT (DHXCT) images (van den Bulcke et al. 2014; de Mil et al. 2016; de Mil and van den Bulcke 2023). The greyscale represents the wood density (kg $m^{-3}$). Species marked with (*) were excluded from the analysis given the substantial proportion of samples with ambiguous tree ring definitions after the crossdating processing. See Table S2 for the list of species per site considered in the analysis.

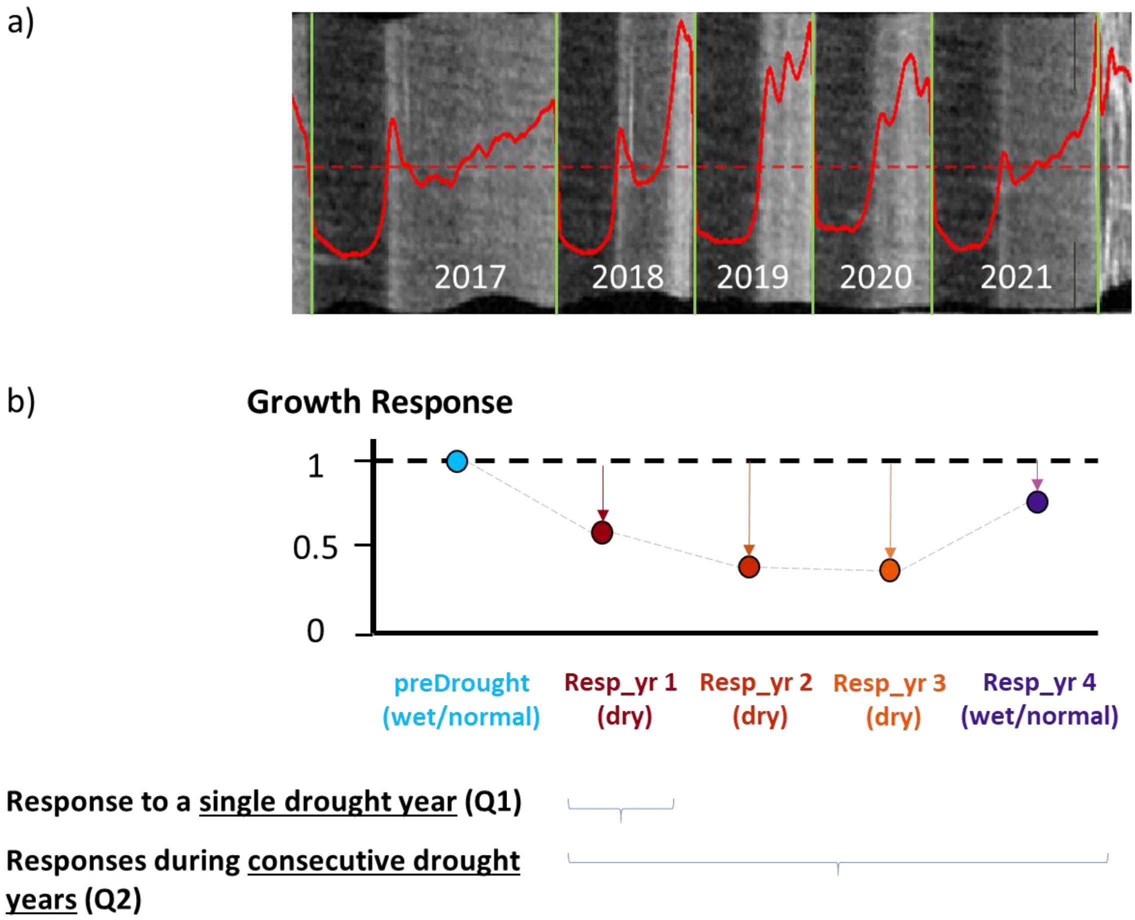


Figure S6. Example of calculation of radial growth responses to consecutive drought years. a) Radial growth responses are estimated on wood cores scanned with X-ray micro-Computed Tomography. We determine annual aboveground biomass increment of growth rings as a proxy of radial growth responses, which was determined based on the basal area increment (derived from the distance between ring boundaries, green lines) and mean wood density (red line showing wood density profile, greyscale indicating wood density estimates). b) Drought-induced growth responses are calculated as the ratio between the growth during the corresponding response year and the pre-drought reference growth. Values below the horizontal line (growth response < 1) representing trees responding with growth reductions during drought relative to the reference pre-drought year. Here, the initial drought year (Resp_yr 1) was followed by two consecutive drought years (Resp_yr 2-3), and a post-drought year with normal or wet conditions (Resp_yr 4). Study analysis focused on the growth responses during the single initial drought year (Q1) and during consecutive years over the prolonged drought period (Q2).

#### Radial growth

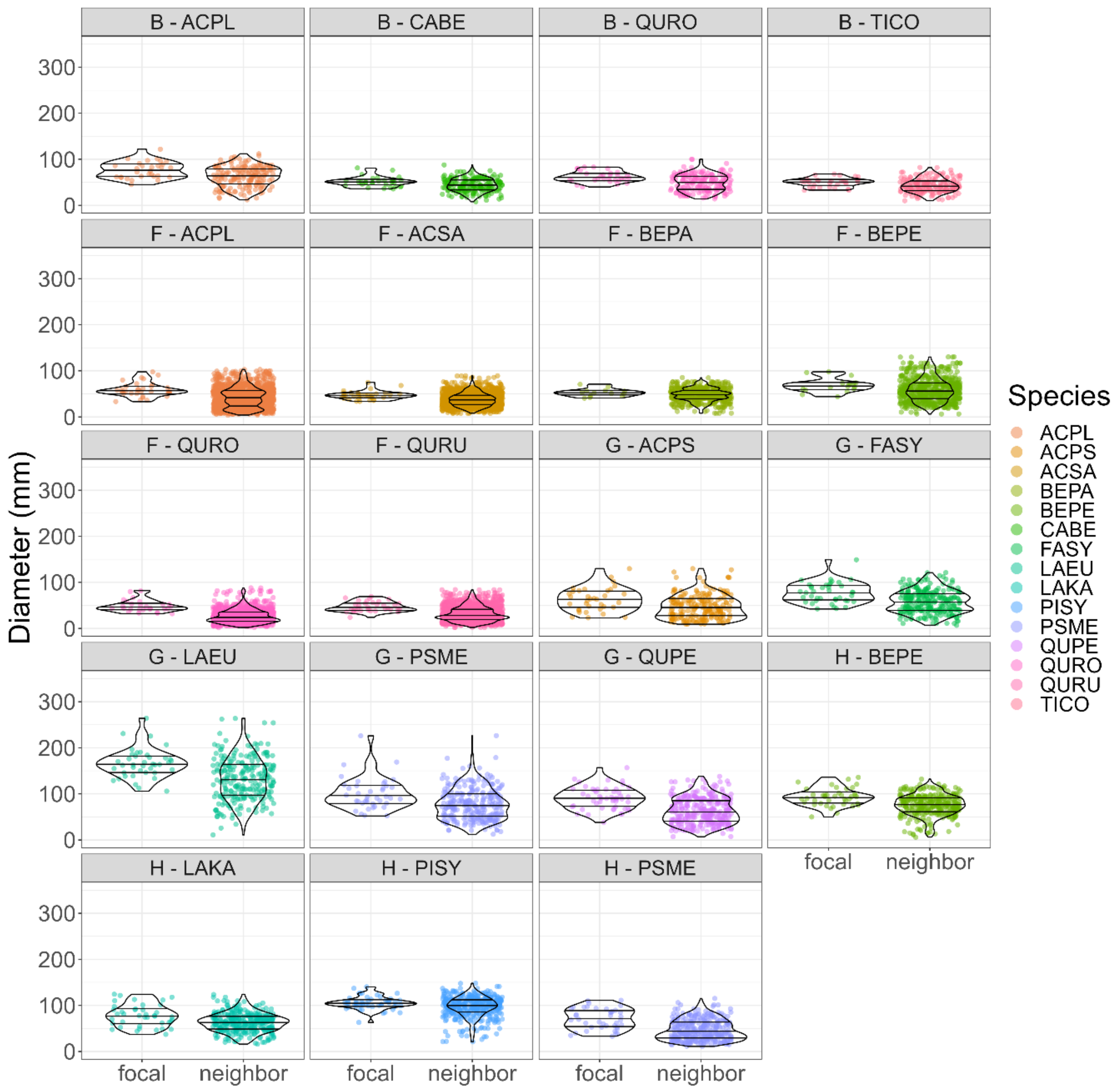


Figure S7. Diameter distributions per species and site. Diameter was measured above bark at the coring height of 30 cm. Distributions include all sampled focal trees and the corresponding neighbors. See Table S2 for the list of species per site considered in the analysis. Horizontal lines within each violin-plot distribution represent the 1$^{st}$, 2$^{nd}$ (median) and 3$^{rd}$ distribution quantiles.

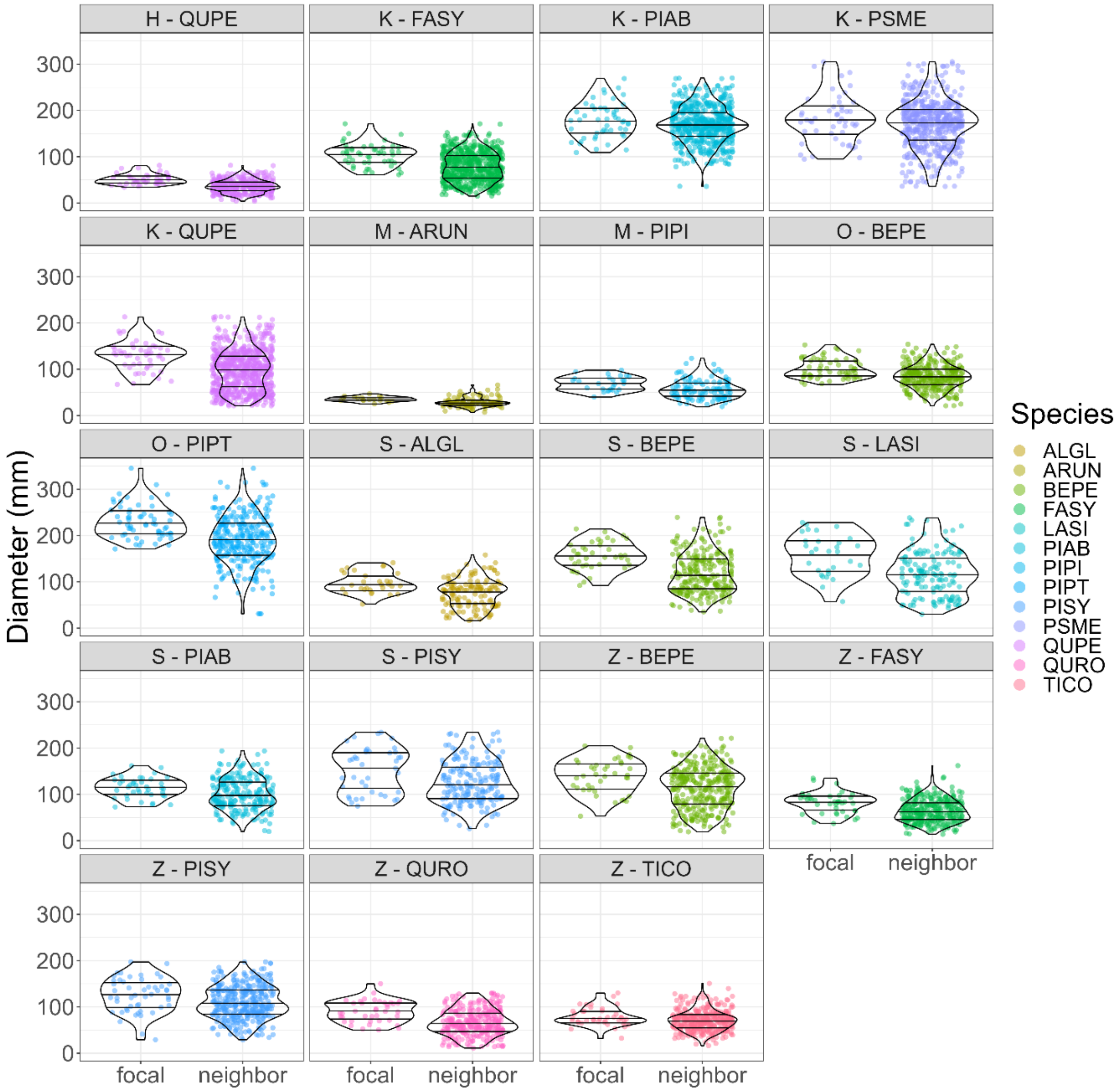


Figure S7 (cont). Diameter distributions per species and site.

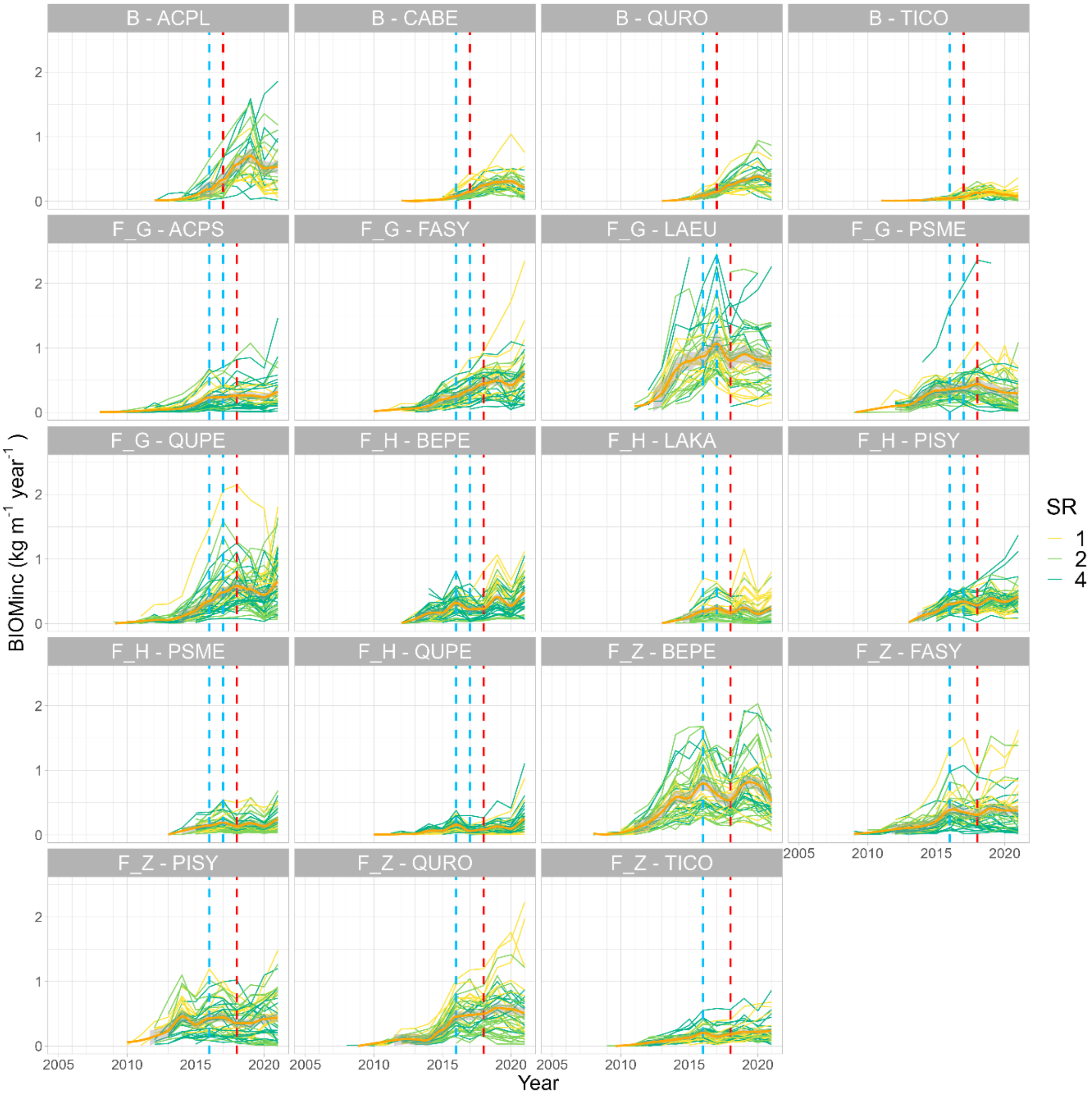


Figure S8. Radial biomass increment (BIOMinc, kg / m$^{-1}$ year$^{-1}$) series of focal trees at individual-tree level per species and site. See Table S2 for the list of species per site considered in the analysis. Legend indicates the species richness (SR) at plot-level for each focal tree. The solid orange line in each plot represents the average BIOMinc trend per species and site estimated via scatterplot smoothing (LOESS) curve, with the grey shaded area surrounding indicating the 95% confidence interval for the fitted values. Vertical dashed lines represent the pre-drought reference year(s) (blue) and the initial drought year (red) considered in the analysis. Figure continue in the following page for the other species and sites.

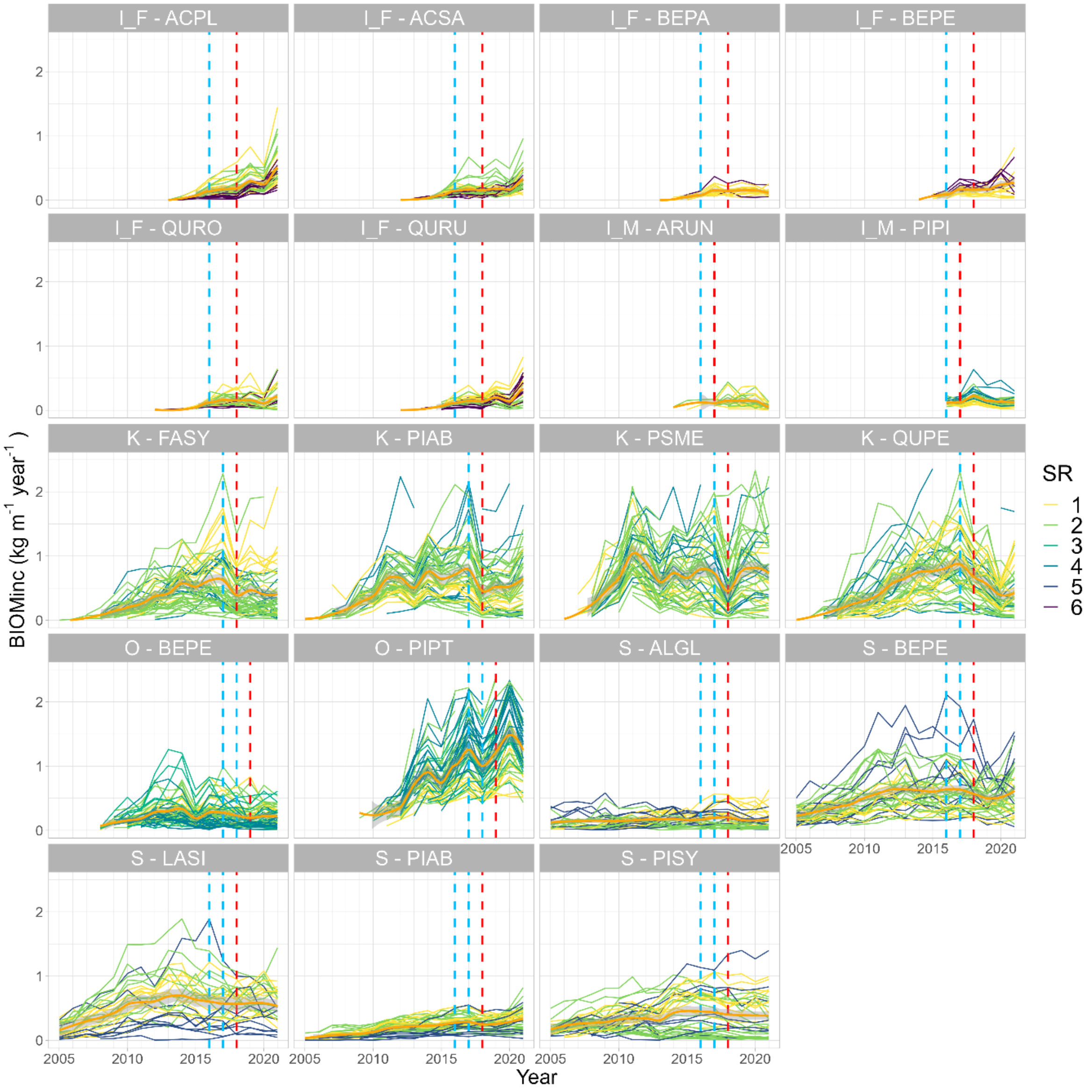


Figure S8. (cont.). Comparison of radial biomass increment (BIOMinc, kg / $m^{-1}$ $year^{-1}$) series per site and species.

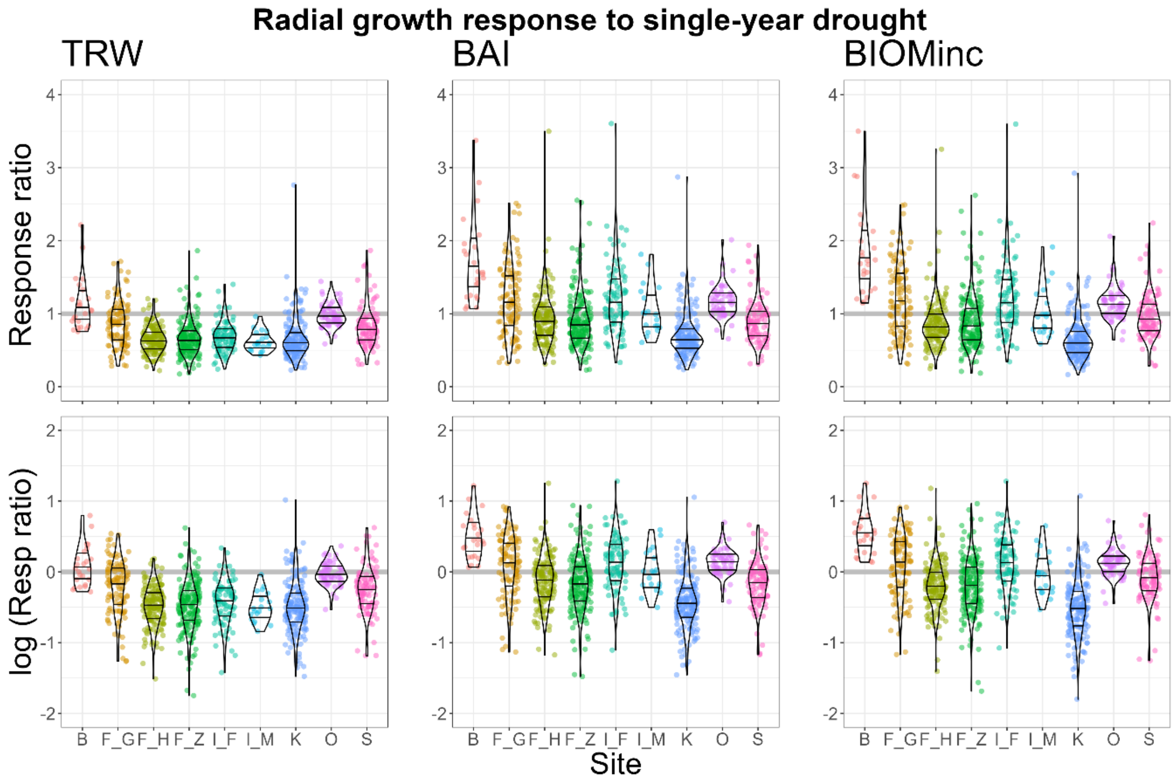


Figure S9. Distributions per site of relative radial growth responses during the single-year drought in terms of tree ring width (TRW, left), basal area increment (BAI, center), and radial biomass increment (BIOMinc, right). Responses are calculated as growth ratio to the reference pre-drought growth (above) and the log-transformation of that ratio (below). Points represent the measured values (or logarithm) of growth responses, with values below the horizontal line (response ratio < 1, logarithm < 0) representing trees with growth reductions during drought compared to the reference pre-drought. Distributions of growth responses per site include all species per site included in Q1 analysis. Note only one species per site was analyzed for Q1 in B, I_F and I_M. See Table S2 for details on species per site. Horizontal lines within each violin-plot distribution represent the 1$^{st}$, 2$^{nd}$ (median) and 3$^{rd}$ distribution quantiles.

#### Functional traits selection

##### Supplementary Method 3. Functional traits selection

We compiled species-level hydraulic traits, leaf traits, and whole-plant economic traits from published databases to test how the functional identity of the tree species and the functional diversity of the neighborhood relate to the tree responses to drought. Hydraulic traits included water potential at which 50% of xylem cavitates (P50, MPa), turgor loss point (TLP, MPa), and hydraulic safety margin ($HSM_{TLP}$, MPa) calculated as the difference between TLP-P50 (Martin-StPaul et al. 2017; Powers et al. 2020). Leaf traits included leaf mass per area (LMA, mg $mm^{-2}$), leaf nitrogen concentration (leafN, mg $g^{-1}$), and leaf size ($cm^2$). Whole plant's economic traits included wood density (WD, g $cm^{-3}$) and potential maximum height (m). We also included seed mass (g) as a trait related to reproduction contributing to the plant's ecological strategy through the stature-recruitment trade-off (Díaz et al. 2016; Westoby 1998; Guillemot et al. 2022).

We considered species-specific mean trait values from consolidated species-level datasets, without considering potential intraspecific trait variability and plasticity within the sites as records of the species-specific trait variability are scarce and incomplete for the studied species within the study mixtures compositions. Leaf and whole plant economic traits were obtained mainly from Díaz et al. (2022), TRY (Kattge et al. 2011) and BIOMASS databases (Réjou-Méchain et al. 2017). For *Pinus pinaster*, we replaced Díaz et al. (2022) data with own data because the value provided for SLA were out the possible range for a pine species. Hydraulic traits were collected from multiple dataset sources. For vulnerability to cavitation we used an updated version of Martin-StPaul et al. (2017) using data from multiple sources (Skelton et al. 2021; Song et al. 2022; Ziegler et al. 2019; Chen et al. 2019; Lobo et al. 2018; Liu et al. 2019; Sjöman et al. 2015), as well as additional recently published data from our group (Decarsin et al. 2024). For leaf hydraulic traits (TLP) we used species-level data from multiple sources (Bartlett et al. 2012; Choat et al. 2018; Guillemot et al. 2022; Kunert and Tomaskova 2020; Maitner et al. 2018), as well as additional own data (Moreno et al. 2024).

Trait datasets included values for all the studied species for leafN, leaf size, maximum height and wood density. Missing P50 values for *Q. pyrenaica* and *L. kaempferi* were imputed using the values of the closest available taxon in our database (which had the same wood density within a +/- 0.1 range), namely *Q. pubescens and L. sibirica,* respectively. Missing TLP values for *L. kaempferi*, *L. sibirica*, *P. pinea*, *Q. pyrenaica* were imputed by using the relationship between P12 (potential at which 12% of xylem cavitates) and TLP found in the whole dataset (as leaf turgor maintains gas exchange while avoiding embolism) (Martin-StPaul et al. 2017).Finally, three remaining missing trait values (LMA for *L. sibirica;* seed mass for *P. pinaster* and *Q. pyrenaica*) were imputed with multiple imputation with chained equations (MICE) using the predicted mean value matching with 500 runs using the R package mice (van Buuren and Groothuis-Oudshoorn 2011). The final dataset of species-specific trait values is detailed in Table S6.

Table S6**.** Functional traits: species-specific mean trait values and scaled values (sc) considered in the analysis. Species in bold correspond with focal trees analyzed for drought responses.

| **Species** | **Sp code** | **Focal tree** | **Tree type** | **P50 (Mpa)** | **P50 sc** | **TLP (Mpa)** | **TLP sc** | **$HSM_{TLP}$ (Mpa)** | **$HSM_{TLP}$ sc** | **LMA (mg / $mm^2$)** | **LMA sc** |
|---|---|---|---|---|---|---|---|---|---|---|---|
| *Acer monspessulanum* | ACMO | neighbor | angiosp | -6.74 | -1.69 | -3.73 | -2.69 | 3.01 | 0.80 | 84.90 | -0.29 |
| ***Acer platanoides*** | **ACPL** | **focal** | **angiosp** | **-4.21** | **-0.07** | **-1.80** | **0.96** | **2.41** | **0.40** | **51.66** | **-0.80** |
| *Acer pseudoplatanus* | ACPS | neighbor | angiosp | -3.28 | 0.53 | -1.83 | 0.89 | 1.45 | -0.23 | 72.12 | -0.48 |
| ***Acer saccharum*** | **ACSA** | **focal** | **angiosp** | **-4.01** | **0.06** | **-1.83** | **0.89** | **2.18** | **0.25** | **46.15** | **-0.89** |
| *Alnus glutinosa* | ALGL | neighbor | angiosp | -1.63 | 1.59 | -1.77 | 1.01 | -0.13 | -1.27 | 68.11 | -0.55 |
| *Arbutus unedo* | ARUN | neighbor | angiosp | -7.84 | -2.40 | -1.26 | 1.96 | 6.58 | 3.14 | 121.24 | 0.27 |
| *Betula papyrifera* | BEPA | neighbor | angiosp | -1.80 | 1.48 | -1.33 | 1.83 | 0.47 | -0.88 | 63.58 | -0.62 |
| ***Betula pendula*** | **BEPE** | **focal** | **angiosp** | **-1.86** | **1.44** | **-1.89** | **0.79** | **-0.03** | **-1.21** | **69.71** | **-0.52** |
| *Carpinus betulus* | CABE | neighbor | angiosp | -3.96 | 0.09 | -2.58 | -0.52 | 1.38 | -0.27 | 51.02 | -0.81 |
| ***Fagus sylvatica*** | **FASY** | **focal** | **angiosp** | **-3.15** | **0.62** | **-2.33** | **-0.05** | **0.81** | **-0.65** | **65.37** | **-0.59** |
| *Fraxinus ornus* | FROR | neighbor | angiosp | -6.36 | -1.45 | -2.22 | 0.16 | 4.14 | 1.54 | 75.23 | -0.44 |
| ***Larix eurolepis*** | **LAEU** | **focal** | **gymnosp** | **-3.16** | **0.61** | **-2.00** | **0.57** | **1.16** | **-0.42** | **118.91** | **0.24** |
| ***Larix kaempferi*** | **LAKA** | **focal** | **gymnosp** | **-3.43** | **0.43** | **-2.66** | **-0.67** | **0.77** | **-0.68** | **63.03** | **-0.62** |
| ***Larix sibirica*** | **LASI** | **focal** | **gymnosp** | **-3.43** | **0.43** | **-2.66** | **-0.67** | **0.77** | **-0.68** | **75.23** | **-0.44** |
| ***Picea abies*** | **PIAB** | **focal** | **gymnosp** | **-3.64** | **0.30** | **-2.68** | **-0.72** | **0.96** | **-0.56** | **225.18** | **1.88** |
| ***Pinus pinaster*** | **PIPT** | **focal** | **gymnosp** | **-3.77** | **0.21** | **-2.00** | **0.57** | **1.77** | **-0.02** | **262.00** | **2.45** |
| ***Pinus pinea*** | **PIPI** | **focal** | **gymnosp** | **-4.34** | **-0.15** | **-2.56** | **-0.49** | **1.78** | **-0.01** | **237.62** | **2.07** |
| ***Pinus sylvestris*** | **PISY** | **focal** | **gymnosp** | **-3.17** | **0.60** | **-2.22** | **0.16** | **0.95** | **-0.56** | **209.68** | **1.64** |
| ***Pseudotsuga menziesii*** | **PSME** | **focal** | **gymnosp** | **-3.46** | **0.42** | **-2.97** | **-1.26** | **0.49** | **-0.87** | **161.13** | **0.89** |
| *Quercus ilex* | QUIL | neighbor | angiosp | -7.01 | -1.86 | -2.18 | 0.23 | 4.83 | 1.99 | 165.29 | 0.96 |
| ***Quercus petraea*** | **QUPE** | **focal** | **angiosp** | **-3.97** | **0.09** | **-2.42** | **-0.22** | **1.55** | **-0.16** | **72.34** | **-0.48** |
| *Quercus pubescens* | QUPU | neighbor | angiosp | -5.22 | -0.72 | -2.84 | -1.01 | 2.38 | 0.38 | 88.74 | -0.23 |
| *Quercus pyrenaica* | QUPY | neighbor | angiosp | -5.22 | -0.72 | -2.68 | -0.71 | 2.54 | 0.49 | 59.84 | -0.67 |
| ***Quercus robur*** | **QURO** | **focal** | **angiosp** | **-4.74** | **-0.41** | **-2.65** | **-0.65** | **2.09** | **0.19** | **72.50** | **-0.48** |
| ***Quercus rubra*** | **QURU** | **focal** | **angiosp** | **-4.43** | **-0.21** | **-2.53** | **-0.43** | **1.90** | **0.07** | **67.44** | **-0.56** |
| ***Tilia cordata*** | **TICO** | **focal** | **angiosp** | **-2.88** | **0.79** | **-2.27** | **0.05** | **0.60** | **-0.79** | **42.17** | **-0.95** |

Table S6 (cont.)

| **Species** | **Sp code** | **leafN (mg / g)** | **leafN sc** | **leaf size (cm²)** | **leaf size sc** | **seed mass (g)** | **seed mass sc** | **height max (m)** | **height max sc** | **WD (g / cm³)** | **WD sc** |
|---|---|---|---|---|---|---|---|---|---|---|---|
| *Acer monspessulanum* | ACMO | 19.56 | 0.18 | 1233 | -0.49 | 0.04 | -0.49 | 7.7 | -1.25 | 0.90 | 2.14 |
| ***Acer platanoides*** | **ACPL** | **19.42** | **0.15** | **7085** | **1.95** | **0.12** | **-0.40** | **21.9** | **-0.15** | **0.58** | **-0.12** |
| *Acer pseudoplatanus* | ACPS | 23.63 | 0.98 | 8457 | 2.52 | 0.08 | -0.45 | 24.5 | 0.05 | 0.56 | -0.27 |
| ***Acer saccharum*** | **ACSA** | **18.05** | **-0.12** | **4736** | **0.97** | **0.06** | **-0.47** | **32.9** | **0.70** | **0.66** | **0.42** |
| *Alnus glutinosa* | ALGL | 30.94 | 2.41 | 3070 | 0.28 | 0.00 | -0.53 | 17.8 | -0.47 | 0.46 | -1.02 |
| *Arbutus unedo* | ARUN | 13.47 | -1.02 | 1507 | -0.38 | 0.12 | -0.40 | 4.5 | -1.49 | 0.78 | 1.29 |
| *Betula papyrifera* | BEPA | 23.83 | 1.02 | 2536 | 0.05 | 0.00 | -0.53 | 14.7 | -0.70 | 0.56 | -0.29 |
| ***Betula pendula*** | **BEPE** | **25.14** | **1.27** | **1136** | **-0.53** | **0.00** | **-0.53** | **12.0** | **-0.91** | **0.61** | **0.09** |
| *Carpinus betulus* | CABE | 21.13 | 0.49 | 2191 | -0.09 | 0.04 | -0.48 | 16.7 | -0.55 | 0.63 | 0.19 |
| ***Fagus sylvatica*** | **FASY** | **22.27** | **0.71** | **2062** | **-0.14** | **0.18** | **-0.33** | **31.5** | **0.58** | **0.68** | **0.57** |
| *Fraxinus ornus* | FROR | 20.20 | 0.30 | 6203 | 1.58 | 0.03 | -0.49 | 7.9 | -1.22 | 0.67 | 0.46 |
| ***Larix eurolepis*** | **LAEU** | **22.65** | **0.78** | **70** | **-0.97** | **0.01** | **-0.53** | **40.0** | **1.24** | **0.40** | **-1.44** |
| ***Larix kaempferi*** | **LAKA** | **16.30** | **-0.47** | **53** | **-0.98** | **0.00** | **-0.53** | **31.1** | **0.56** | **0.43** | **-1.22** |
| ***Larix sibirica*** | **LASI** | **14.66** | **-0.79** | **70** | **-0.97** | **0.01** | **-0.52** | **35.7** | **0.91** | **0.47** | **-0.97** |
| ***Picea abies*** | **PIAB** | **12.36** | **-1.24** | **32** | **-0.99** | **0.01** | **-0.52** | **40.7** | **1.29** | **0.42** | **-1.27** |
| ***Pinus pinaster*** | **PIPT** | **10.13** | **-1.68** | **254** | **-0.90** | **0.04** | **-0.48** | **21.8** | **-0.16** | **0.53** | **-0.53** |
| ***Pinus pinea*** | **PIPI** | **11.50** | **-1.41** | **138** | **-0.94** | **0.67** | **0.21** | **27.4** | **0.27** | **0.58** | **-0.14** |
| ***Pinus sylvestris*** | **PISY** | **12.87** | **-1.14** | **72** | **-0.97** | **0.01** | **-0.52** | **25.9** | **0.15** | **0.54** | **-0.43** |
| ***Pseudotsuga menziesii*** | **PSME** | **11.84** | **-1.34** | **31** | **-0.99** | **0.02** | **-0.51** | **65.3** | **3.18** | **0.46** | **-1.03** |
| *Quercus ilex* | QUIL | 13.98 | -0.92 | 763 | -0.68 | 1.83 | 1.50 | 9.3 | -1.12 | 0.90 | 2.14 |
| ***Quercus petraea*** | **QUPE** | **19.87** | **0.24** | **2986** | **0.24** | **1.50** | **1.14** | **31.4** | **0.58** | **0.58** | **-0.16** |
| *Quercus pubescens* | QUPU | 17.23 | -0.28 | 2892 | 0.20 | 1.79 | 1.45 | 15.8 | -0.62 | 0.71 | 0.78 |
| *Quercus pyrenaica* | QUPY | 17.56 | -0.22 | 3026 | 0.26 | 0.04 | -0.49 | 20.0 | -0.30 | 0.83 | 1.60 |
| ***Quercus robur*** | **QURO** | **21.81** | **0.62** | **2992** | **0.24** | **2.86** | **2.64** | **27.1** | **0.25** | **0.58** | **-0.14** |
| ***Quercus rubra*** | **QURU** | **21.01** | **0.46** | **6217** | **1.58** | **2.95** | **2.75** | **18.3** | **-0.43** | **0.66** | **0.40** |
| ***Tilia cordata*** | **TICO** | **23.93** | **1.04** | **2818** | **0.17** | **0.04** | **-0.49** | **18.8** | **-0.39** | **0.45** | **-1.07** |

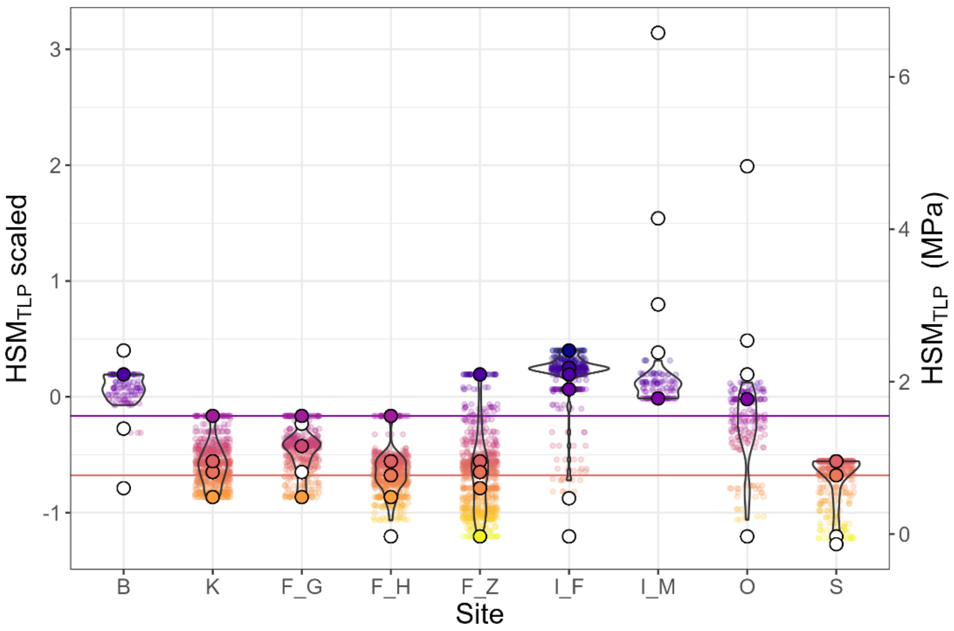


Figure S10**.** $HSM_{TLP}$ ranges of focal-trees (trait identity) and neighborhood weighted means (NWM) per study site considered in the analysis. Colored circles represent the species-specific $HSM_{TLP}$ values of the focal trees and white circles correspond to the species-specific values of the neighbor trees (see Table S6 for details on the trait values for each species). Colored points with jitter represent the neighborhood-weighted mean ($NWM_{HSM}$, weighted by basal area) for the focal trees' neighborhoods, with violin points representing its distribution of values. Right vertical axis represents the trait dataset values, while left vertical axis corresponds with the scaled values used in the mixed-model analysis. Standardization was done by scaling-centering across all species (focal and neighbors) and sites. Horizontal lines represent the $HSM_{TLP}$ values for 1$^{st}$ and 3$^{rd}$ quartile across all analyzed samples that were used for visualization of marginal effects across all models and sites in Figure 6.

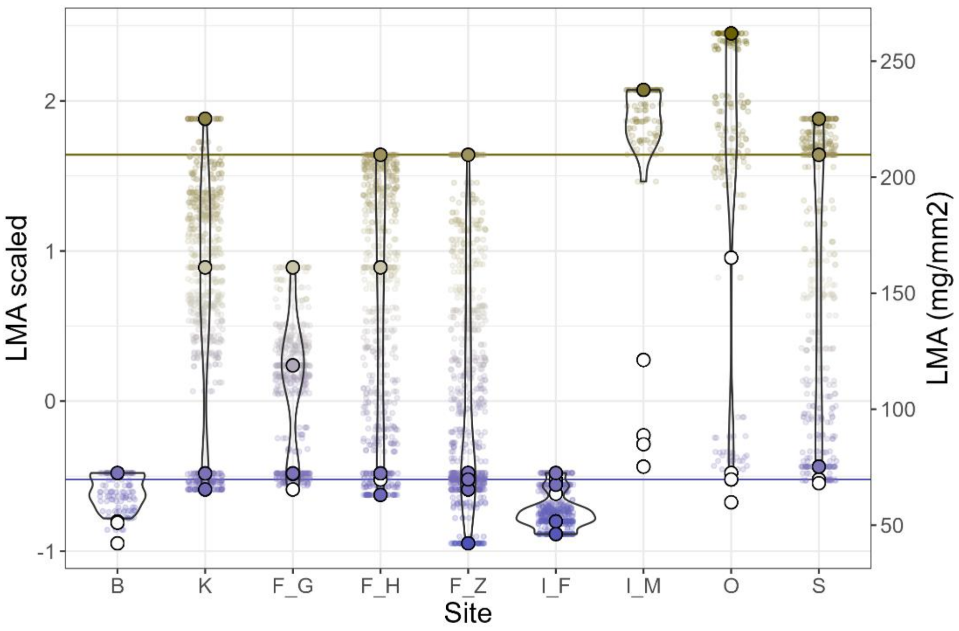


Figure S11**.** LMA ranges of focal-trees (trait identity) and neighborhood weighted means (NWM) per study site considered in the analysis. Colored circles represent the species-specific LMA values of the focal trees and white circles correspond to the species-specific values of the neighbor trees (seeTable S6 for details on the trait values for each species). Colored points with jitter represent the neighborhood-weighted mean ($NWM_{LMA}$, weighted by basal area) for the focal trees' neighborhoods, with violin points representing its distribution of values. Right vertical axis represents the trait dataset values, while left vertical axis corresponds with the scaled values used in the mixed-model analysis. Standardization was done by scaling-centering across all species (focal and neighbors) and sites. Color legend represents the range of LMA values of the focal trees. Horizontal lines represent the LMA scaled values for 1$^{st}$ and 3$^{rd}$ quartile across all analyzed samples.

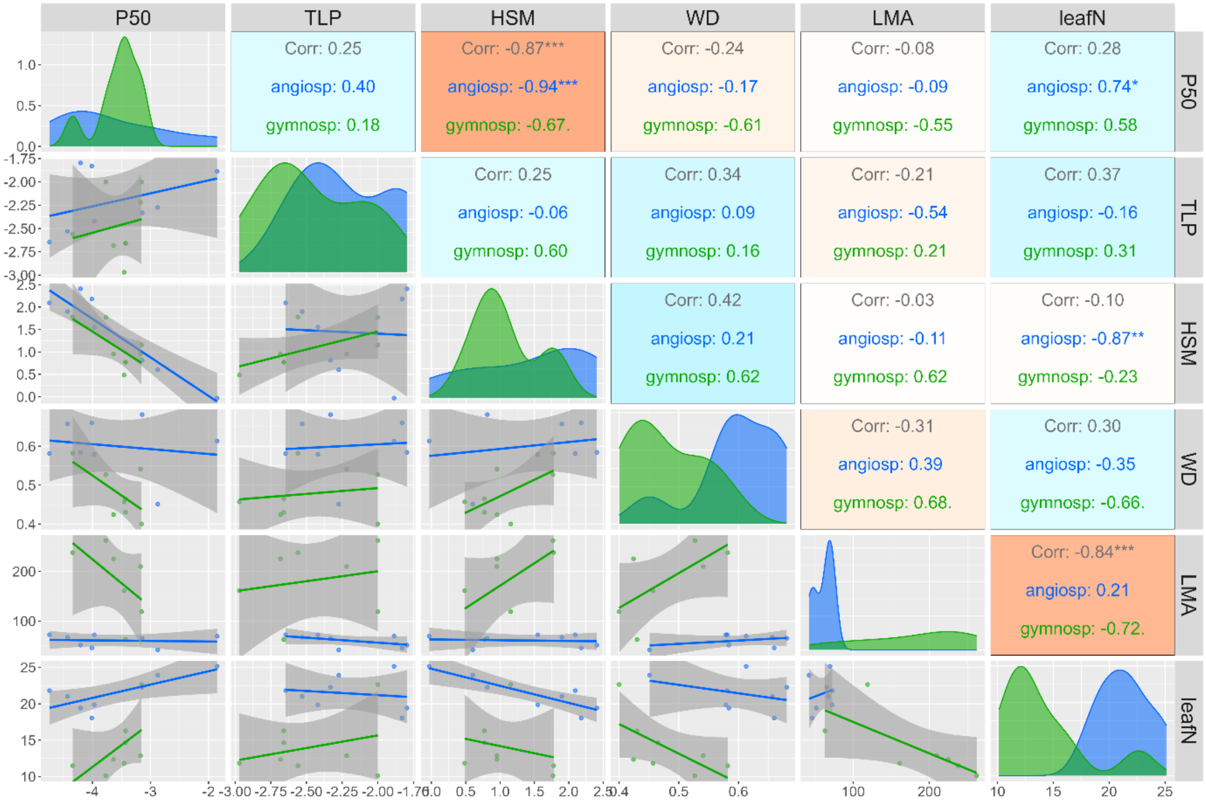


Figure S12. Pairwise correlations between the functional traits with most dataset coverage across the focal-tree species. Panel plots across the diagonal represent the density distributions of each variable, with colors representing tree type (angiosperms trees in blue, gymnosperms in green). Lower panel plots correspond to the bivariate scatter plots with smoothed lines per tree type. Upper plots correspond to the bivariate Pearson's r correlation coefficients, with the blue (red) color palette representing higher positive (negative) correlations, with significance levels * p<0.05, ** p<0.01, *** p<0.001. See Table S6 for details on the trait values for each species.

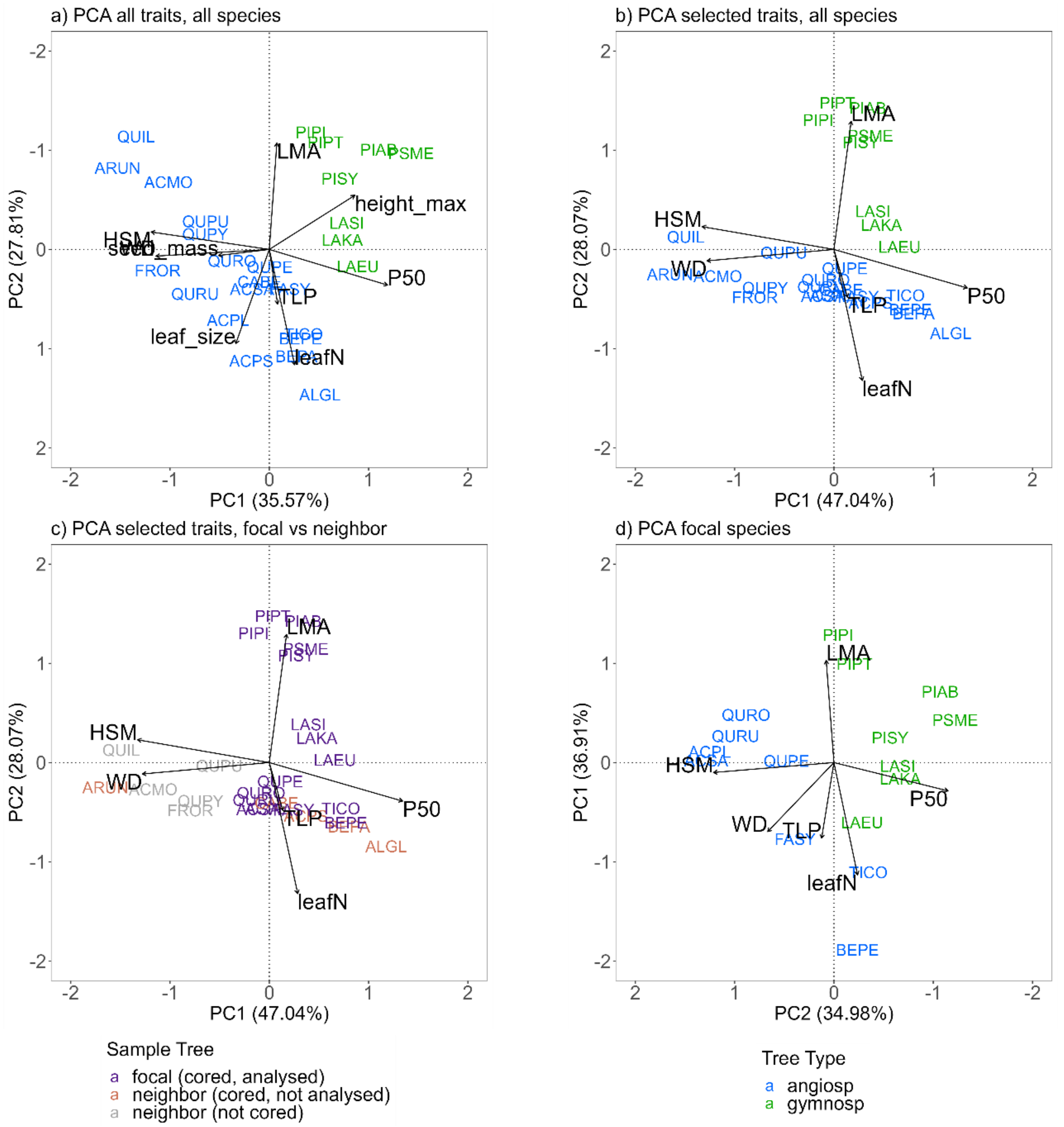


Figure S13. Principal component analysis (PCA) biplots for the selection of functional trait gradients, comparing different subsets of functional traits and species: (a) PCA for all functional traits and species in the dataset, (b) subset of the main traits with the best coverage and PCA alignment (i.e. excluding leaf size, seed mass, and potential maximum height), (c) subset of main traits with differentiation between focal and neighbor tree species, and (d) subset of main traits only for focal tree species. See Table S6 for details on the species lists and trait values for each species. PCA was performed with the rda function in the vegan package (Oksanen et al. 2012). Varimax rotated principal components (R Core Team 2023) were used to improve the alignment of the two orthogonal trait gradients with the first and second PCA axis. Vertical PC2 axis of subplot (a) was reversed, and PCA axis of subplot (d) were flipped, to coincide with the vertical and horizontal axis of other plots. PCA partitioned the variation among functional traits into two orthogonal trait gradients associated with different ecological strategies. PC1 axis aligned with $HSM_{TLP}$, WD and (opposite with) P50 explained most of the variation across traits. These traits have been associated with drought tolerance and provide a mechanistic explanatory framework for vulnerability to drought (Choat et al. 2012; McDowell et al. 2008; Kröber et al. 2014; Anderegg et al. 2018). PC2 axis was aligned with LMA and (opposite with) leafN, associated to the plant economics spectrum (Reich et al. 1999; Wright et al. 2004). Alignment of trait gradients with PCA axis was maintained when PCA were performed for all functional traits in the dataset (a)

compared to a subset of the main traits with the best coverage and PCA alignment (b), as well as for all the neighborhood species (c) compared to only the analyzed focal tree species (d).

Supplementary Method 4. Sensitivity analysis for the effect of functional traits on the growth responses during the single-year drought

Prior to the final model used for the analysis, we compared the effect of $HSM_{TLP}$ on the growth responses during the single-year drought to the effect of other main functional traits (i.e. traits aligned with the PCA axis) with a sensitivity analyses. We fitted separate linear mixed-effect models (LMMs) to model the growth responses during the single-year drought (i.e. in terms of TRW, BAI or BIOMinc) as a function of each species-specific trait. We used a modelling approach where single traits represent ecological strategies, as opposed to using a multivariate approach (i.e. integrated PCA scores). We accounted for the experimental design and differences between sites through a nested random effect structure of plot nested within site. We used a log transformation of the growth responses to normalise residuals, and centred and scaled all functional trait predictors (via subtracting μ and dividing by σ) before analysis to compare fixed effect estimates. We compared the model performance among the different single-trait models based on the Akaike Information Criterion (AIC). HSM-based models were selected as the most parsimonious and with higher effect (absolute coefficient estimate) of all single-trait models, being consistent across the different growth response models (TRW, BAI, BIOMinc) (Table S7, Figure S14). Models based on P50 and WD have similar AIC than the $HSM_{TLP}$ models, but their trait values were highly correlated with $HSM_{TLP}$ (Figure S12). Based on this, we used $HSM_{TLP}$ in the following analysis to determine the functional identity of the tree species and functional diversity of the neighborhood, as the key physiological trait to quantify the effect of the drought tolerance gradient on the growth responses.

Table S7. Comparison of the single-trait linear-mixed effects models ranked according to their second-order Akaike's information criterion (AICc).

| **m1.0 - TRW model selection** | | | | | |
|---|---|---|---|---|---|
| **Model** | **K** | **AICc** | **ΔAICc** | **AICc weight** | **log-Likelihood** |
| **HSM** | **5** | **526.52** | **0** | **0.9** | **-258.23** |
| P50 | 5 | 530.91 | 4.39 | 0.1 | -260.42 |
| LMA | 5 | 539.52 | 13 | 0 | -264.73 |
| Height_max | 5 | 550.61 | 24.09 | 0 | -270.27 |
| leafN | 5 | 564.03 | 37.51 | 0 | -276.98 |
| WD | 5 | 566.28 | 39.76 | 0 | -278.11 |
| TLP | 5 | 571.18 | 44.66 | 0 | -280.56 |

| **m1.0 - BAIin model selection** | | | | | |
|---|---|---|---|---|---|
| **Model** | **K** | **AICc** | **ΔAICc** | **AICc weight** | **log-Likelihood** |
| HSM | 5 | 772.65 | 0 | 0.94 | -381.29 |
| P50 | 5 | 777.99 | 5.34 | 0.06 | -383.96 |
| LMA | 5 | 807.32 | 34.66 | 0 | -398.63 |
| Height_max | 5 | 807.89 | 35.24 | 0 | -398.92 |
| leafN | 5 | 822.75 | 50.1 | 0 | -406.34 |
| WD | 5 | 828.22 | 55.57 | 0 | -409.08 |
| TLP | 5 | 831.7 | 59.05 | 0 | -410.82 |

| **m1.0 - BIOMinc model selection** | | | | | |
|---|---|---|---|---|---|
| **Model** | **K** | **AICc** | **ΔAICc** | **AICc weight** | **log-Likelihood** |
| **HSM** | **5** | **744.98** | **0** | **1** | **-367.46** |
| P50 | 5 | 758.22 | 13.24 | 0 | -374.08 |
| WD | 5 | 782.98 | 38 | 0 | -386.46 |
| Height_max | 5 | 783.65 | 38.67 | 0 | -386.79 |
| LMA | 5 | 788.42 | 43.44 | 0 | -389.18 |
| leafN | 5 | 798.12 | 53.14 | 0 | -394.03 |
| TLP | 5 | 803.18 | 58.2 | 0 | -396.56 |

*Table note: Individual models were fitted separately for each growth response and functional trait. Response variables were log-transformed growth response in terms of TRW, BAI and BIOMinc, respectively. All functional traits were standardized to compare the effects across single-trait models, with standardization via scaling-centering across the functional trait dataset of species-specific values including all focal tree and neighbor tree species. Linear-mixed effects models were fitted using maximum likelihood estimation (ML, for AICc comparison among models) with package lme4 (Bates et al. 2015). AICc values were calculated with package AICcmodavg (Mazerolle 2023).*

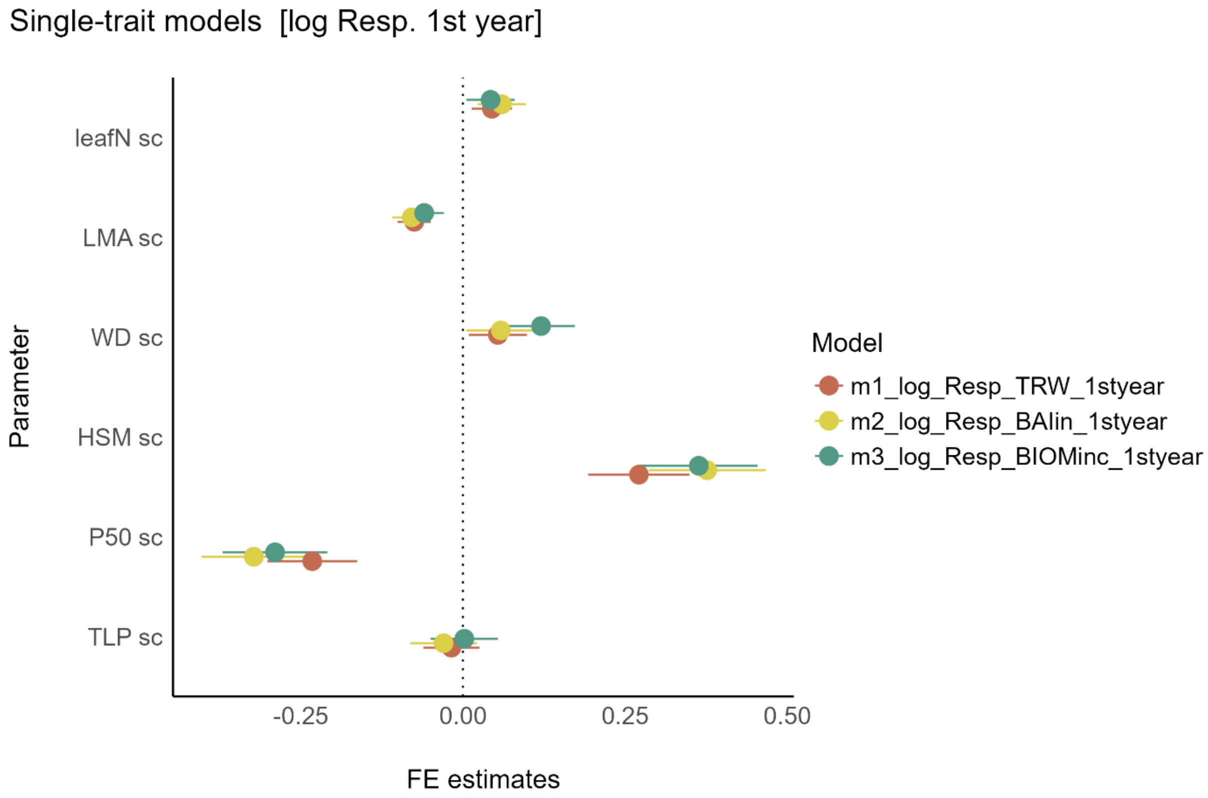


Figure S14. Comparison of the linear-mixed effects estimate parameters of the single-trait models. Individual models were fitted separately for each radial-growth response and main functional trait. Response variables were log-transformed resistance in terms of TRW, BAI and BIOMinc, respectively. All functional traits were standardized to compare the effects across single-trait models (standardization via scaling-centering across the functional trait dataset of species-specific values including all focal tree and neighbor tree species).

#### Neighborhood diversity

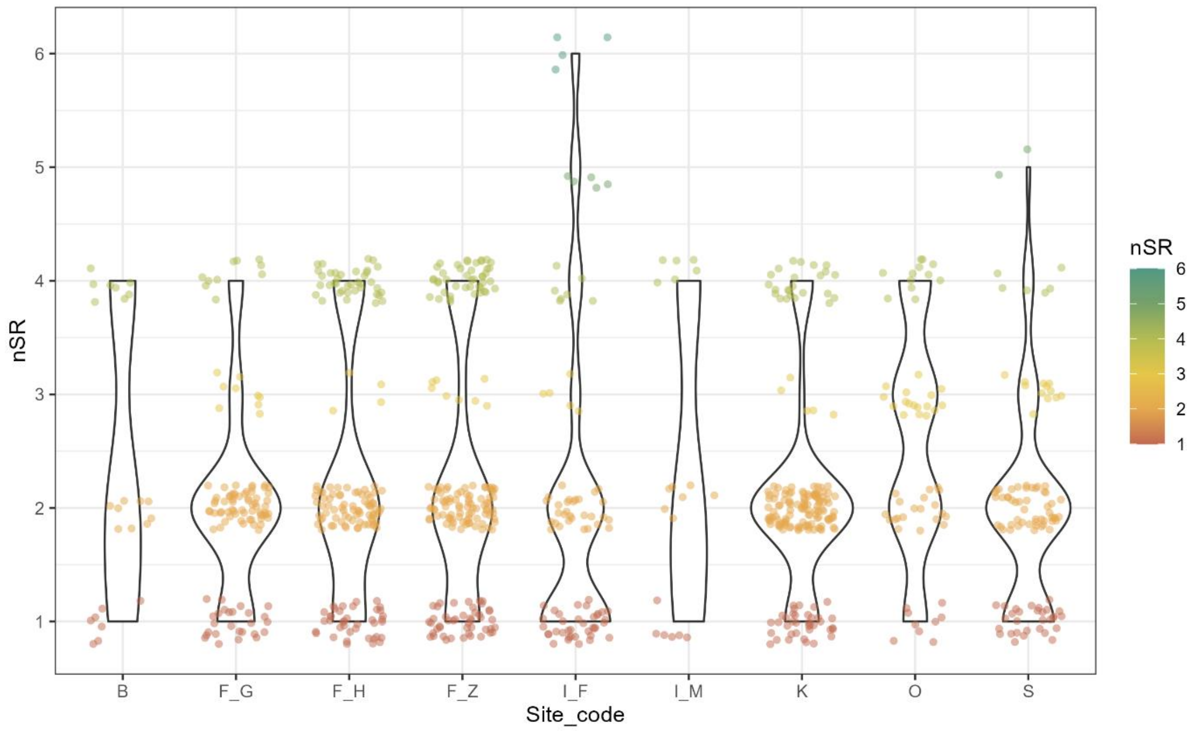


Figure S15. Neighborhood species richness (nSR) value range per study site. Colored points and legend represent the nSR values for the focal trees' neighborhoods.

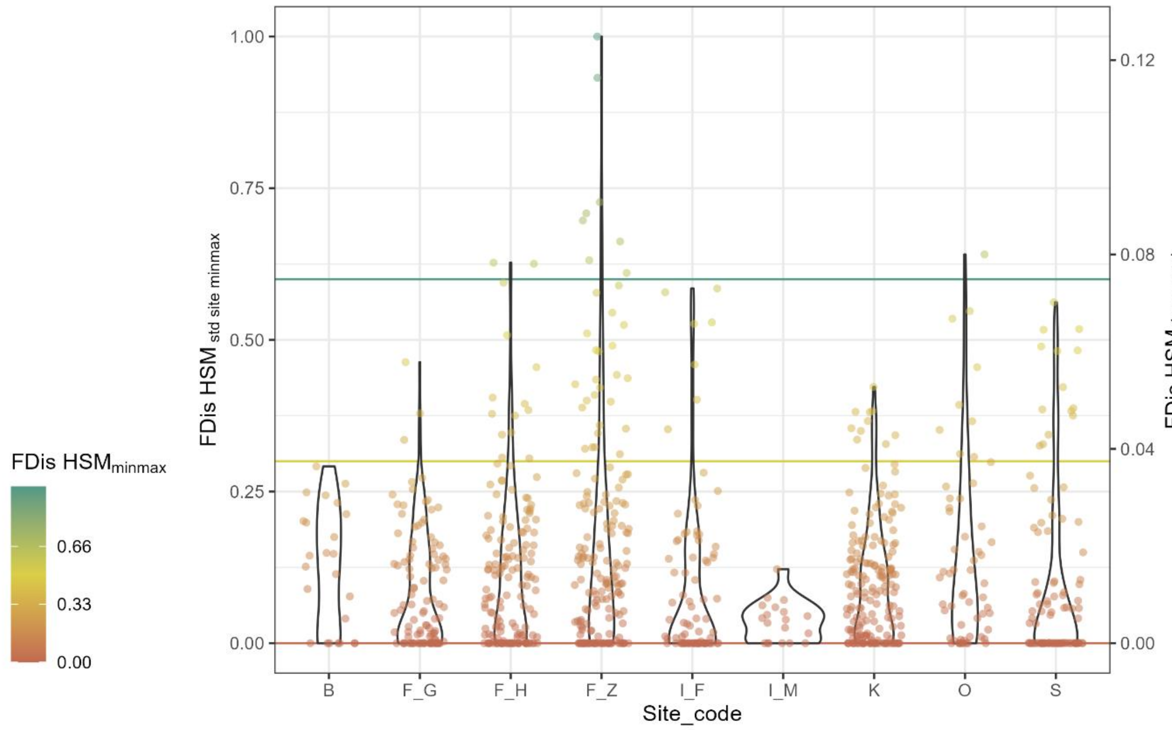


Figure S16. Neighborhood functional diversity of hydraulic safety margin ($FD_{HSM}$) per study site. Right vertical axis represents the trait dataset values, while left vertical axis corresponds with the scaled values used in the mixed-model analysis ($FD_{HSM\ std\_site\_minmax}$). Standardization was done via min-max normalization across the focal tree neighborhood dataset of all sites. Color points and legend represent the range of $FD_{HSM\ std\_site\_minmax}$ values of the focal trees' neighborhoods. Legend breaks and corresponding horizontal lines represent the $FD_{HSM\ std\_site\_minmax}$ values used for visualization of marginal effects across all models and sites in Figure 4 and Figure 5.

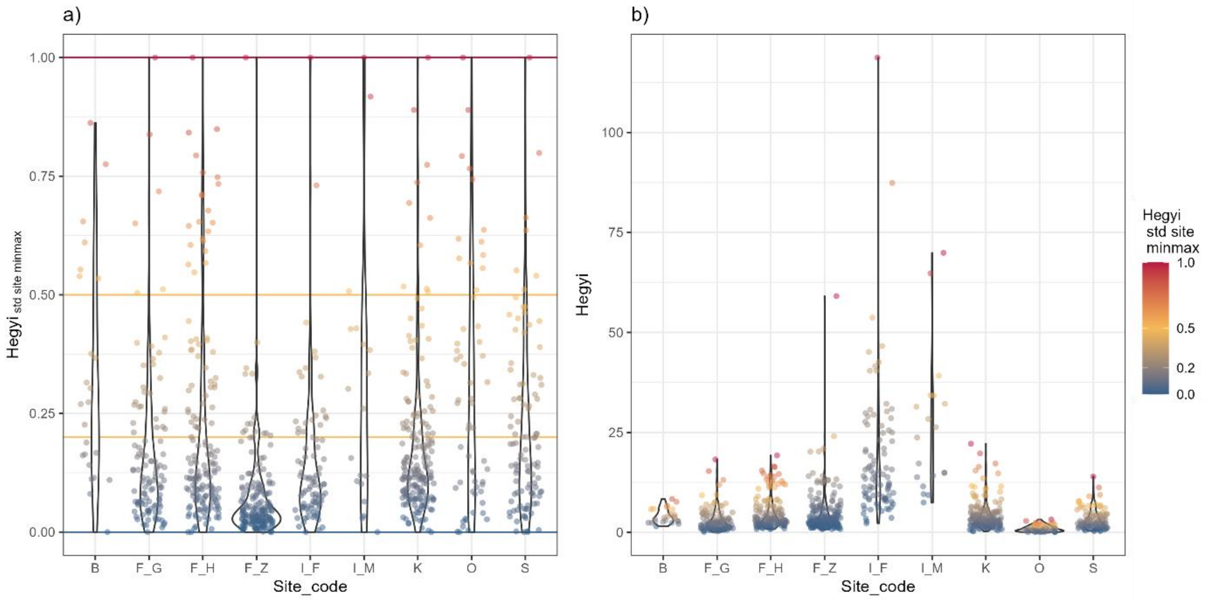


Figure S17. Neighborhood competition (Hegyi) per study site. Right plot (a) represents the trait dataset values, while left plot corresponds (b) with the standardized values used in the mixed-model analysis ($Hegyi_{std\_site\_minmax}$). Standardization was done via min-max normalization per site ($Hegyi_{std\_site\_minmax}$) to account for the relative interspecific competition within each site, instead of absolute differences between sites. Color points and legend represent the range of $Hegyi_{std\_site\_minmax}$ values of the focal trees' neighborhoods. Legend breaks and corresponding horizontal lines represent the $Hegyi_{std\_site\_minmax}$ values used for visualization of marginal effects across all models and sites.

#### Growth responses during a single-year drought

**Supplementary Method 5. Model variables standardization and alternative random structures**

All models used a log transformation of the response variables to improve normalization of residuals in comparison to the alternative untransformed variables and square-root transformation. Model assumptions (normality and heteroscedasticity) were visually checked via quantile-quantile plots and examining model residuals.

For all Q1 and Q2 models, $HSM_{TLP}$ and $FD_{HSM}$ were standardized across the full dataset for all sites to have comparable effect sizes across site-specific models. $HSM_{TLP}$ was standardized by scaling-centering across the functional trait dataset of species-specific values including all focal tree and neighbor tree species (Figure S10, Figure S11). $FD_{HSM}$ was standardized via min-max normalization ($FD_{trait\_minmax}$) across the full dataset of focal tree neighborhoods for all sites (Figure S16).

All models controlled for fixed effects of the relative intraspecific differences in tree size within each site and species ($BA_{std\_sp\_site}$) and the relative interspecific competition within each site ($Hegyi_{std\_site}$). Tree size was standardized by scaling-centering basal area per species and site ($BA_{std\_sp\_site}$), to account for relative intraspecific differences in tree size and not absolute differences between species and sites (Figure S7). Hegyi index was standardized via min-max normalization per site ($Hegyi_{std\_site\_minmax}$) to account for the relative competition within each site, instead of absolute differences between sites, due to different initial planting densities among experiments (Figure S17). Alternative Hegyi standardization by scaling-centering per site was tested but models results did not differ. Both $Hegyi_{std\_site\_minmax}$ and $BA_{std\_sp\_site}$ were calculated from field measurements during the coring year (corresponding to the post-drought Resp_yr 4), assuming not significant changes in competition between response years, as annual tree inventory data was not available for all analysis years. Both $Hegyi_{std\_site\_minmax}$ and $BA_{std\_sp\_site}$ were included as predictors in all models after testing for non-multicollinearity in regression analysis (i.e. variance inflation factor, low VIF < 5 or moderate VIF < 10, with only relative correlation between variables $r = -0.54$, $p < 0.001$; Figure S18).

We compared the model performance and parsimony of fixed effects among the separate models based on the Akaike Information Criterion (AIC). Alternative random effect structures including random slope on the effect of each predictor were tested, but did not show significant effects. Alternative crossed random intercept of species was tested but not considered as it reduced the fixed effect of the species-specific functional trait, neither a random intercept for plot to avoid singular fit due to the limited number of trees per plot. Alternative variance structures (i.e. power, exponential) and optimization methods to model heteroscedasticity were also tested but did not improve the model assumptions nor parsimony.

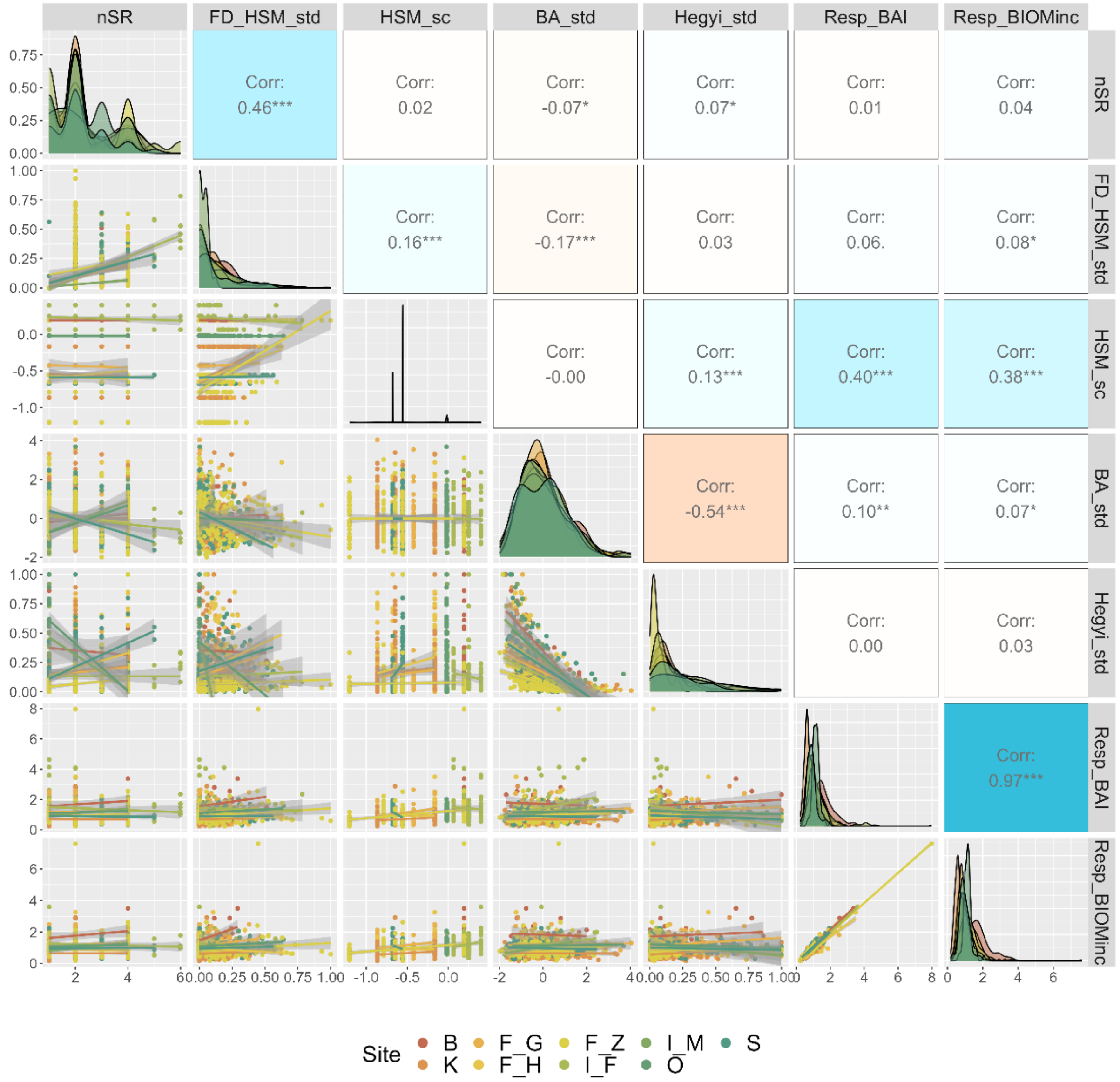


Figure S18. Pairwise correlations between the radial-growth responses during the single-year drought used for the Q1 model (Resp_BIOMinc, Resp_BAI,) and the model predictors neighborhood species richness (nSR), neighborhood functional diversity in hydraulic safety margin ($FD_{HSM\_std\_min-max}$), hydraulic safety margin of the focal-tree species ($HSM_{sc}$), focal-tree basal area ($BA_{std\_sp\_site}$), and neighborhood competition ($Hegyi_{std\_site\_minmax}$). Predictors were standardized to meet model assumptions and have comparable effect sizes across sites (standardization details in Supplementary Method 5). Panel plots across the diagonal represent the density distributions of each variable with different colors per site. Lower panel plots correspond to the bivariate scatter plots with linear regressions per site (color legend). Upper plots correspond to the bivariate Pearson's r correlation coefficients, with the blue (red) color palette representing higher positive (negative) correlations, with significance levels * p<0.05, ** p<0.01, *** p<0.001.

Table S8. Comparison of mixed-effect model statistics for the growth responses during a single-year drought in terms of biomass increment (BIOMinc) and basal area increment (BAI), considering the effects of the drought tolerance (as hydraulic safety margin of the focal-tree species, $HSM_{TLP}$) and the neighborhood tree diversity as species richness (nSR, m1a) and functional trait diversity ($FD_{HSM}$, m1b).

| | **m1.a Resp BIOMinc [log] (1st year)** | | **m1.a Resp BAI [log] (1st year)** | | **m1.b Resp BIOMinc [log] (1st year)** | | **m1.b Resp BAI [log] (1st year)** | |
|---|---|---|---|---|---|---|---|---|
| *Predictors* | *Estimates* | *CI* | *Estimates* | *CI* | *Estimates* | *CI* | *Estimates* | *CI* |
| (Intercept) | 0 | -0.15 – 0.16 | 0.03 | -0.11 – 0.16 | 0.05 | -0.08 – 0.19 | 0.08 | -0.04 – 0.19 |
| BA | **0.04 **** | **0.02 – 0.07** | **0.04 **** | **0.01 – 0.06** | **0.05 **** | **0.02 – 0.07** | **0.04 **** | **0.01 – 0.07** |
| Hegyi | 0.09 | -0.08 – 0.26 | 0.05 | -0.11 – 0.22 | 0.08 | -0.09 – 0.24 | 0.04 | -0.12 – 0.20 |
| HSM | **0.22 *** | **0.05 – 0.39** | **0.24 **** | **0.07 – 0.41** | **0.37 ***** | **0.24 – 0.49** | **0.39 ***** | **0.27 – 0.51** |
| nSR | 0.02 | -0.02 – 0.06 | 0.02 | -0.02 – 0.05 | | | | |
| HSM × nSR | 0.05 | -0.01 – 0.12 | 0.05 | -0.02 – 0.11 | | | | |
| FDis HSM | | | | | 0.01 | -0.22 – 0.24 | -0.02 | -0.24 – 0.21 |
| HSM × $FDis_{HSM}$ | | | | | -0.24 | -0.72 – 0.25 | -0.28 | -0.75 – 0.20 |
| **Random Effects** | | | | | | | | |
| $\sigma^2$ | 0.11 | | 0.1 | | 0.11 | | 0.1 | |
| $\tau_{00}$ | 0.02 Plot:Site | | 0.03 Plot:Site | | 0.02 Plot:Site | | 0.03 Plot:Site | |
| | 0.03 Site | | 0.02 Site | | 0.03 Site | | 0.02 Site | |
| ICC | 0.33 | | 0.32 | | 0.34 | | 0.32 | |
| N | 69 Plot | | 69 Plot | | 69 Plot | | 69 Plot | |
| | 9 Site | | 9 Site | | 9 Site | | 9 Site | |
| Observations | 912 | | 912 | | 912 | | 912 | |
| Marginal $R^2$ / Conditional $R^2$ | 0.109 / 0.407 | | 0.124 / 0.402 | | 0.112 / 0.413 | | 0.132 / 0.413 | |
| AIC | 490.1 | | 469.43 | | 483.624 | | 462.461 | |

** p<0.05 ** p<0.01 *** p<0.001*

*Table note: Response variables are log-transformed growth response in terms of BAI and BIOMinc. Individual models were fitted separately to test different approaches of the effect of the neighborhood tree diversity in terms of species richness (nSR, m1a) and hydraulic functional diversity ($FD_{HSM}$, m1b). Tree size is represented by $BA_{std\_sp\_site}$ (standardized per species and site) and neighborhood competition is represented by $Hegyi_{std\_site\_minmax}$ (standardized via min-max normalization per site). $HSM_{TLP}$ was standardized by scaling-centering across the functional trait dataset of species-specific values including all focal tree and neighbor tree species. $FD_{HSM}$ is represented by the neighborhood functional diversity in terms of $HSM_{TLP}$ and was standardized via min-max normalization across all sites to account for absolute differences between sites. Values correspond with the estimates of regression coefficients and 95% confidence intervals (CI) for each predictor variable. Significant fixed effects with $p<0.05$ are highlighted in bold. Random effect variance is represented by the variance of the model residuals ($\sigma^2$), random intercept variance ($\tau_{00}$), intraclass correlation coefficient (ICC) and the number of groups/observations (N). Linear-mixed effects models were fitted using restricted maximum likelihood estimation (REML) with lme4 R-package(Bates et al. 2015). Table statistics were created using the sjPlot R-package (Lüdecke et al. 2024).*

Table S9. Comparison of mixed-effect model statistics for the growth responses (BIOMinc) during a single-year drought considering the modulating effects of drought intensity (in terms of SPEI12, SPEI6sept, or minimum monthly REW during the growing season) on the interaction between the drought tolerance (hydraulic safety margin $HSM_{TLP}$) of the focal-tree species and the neighborhood tree diversity (species richness nSR, or hydraulic trait diversity $FD_{HSM}$).

| **Response BIOMinc** [log] | **m1a (nSR * HSM)** | | **m1a (nSR * HSM * SPEI12)** | | **m1a (nSR * HSM * SPEI6sept)** | | **m1a (nSR * HSM * REWmin)** | |
|---|---|---|---|---|---|---|---|---|
| *Predictors* | *Estimates* | *CI* | *Estimates* | *CI* | *Estimates* | *CI* | *Estimates* | *CI* |
| (Intercept) | 0 | -0.15 – 0.16 | 0.6 | -0.05 – 1.25 | **0.98 *** | **0.01 – 1.96** | -0.01 | -0.16 – 0.14 |
| BA sp | **0.04 ** ** | **0.02 – 0.07** | **0.04 ** ** | **0.01 – 0.07** | **0.04 ** ** | **0.01 – 0.07** | **0.05 ** ** | **0.02 – 0.07** |
| Hegyi | 0.09 | -0.08 – 0.26 | 0.1 | -0.07 – 0.27 | 0.09 | -0.08 – 0.26 | 0.09 | -0.08 – 0.25 |
| HSM | **0.22 *** | **0.05 – 0.39** | 1.27 | -0.60 – 3.13 | 0.52 | -0.96 – 2.01 | 0.13 | -0.05 – 0.31 |
| nSR | 0.02 | -0.02 – 0.06 | -0.12 | -0.28 – 0.05 | -0.1 | -0.38 – 0.19 | 0.02 | -0.01 – 0.06 |
| HSM × nSR | 0.05 | -0.01 – 0.12 | -0.49 | -1.15 – 0.17 | -0.28 | -0.84 – 0.29 | 0.06 | -0.01 – 0.12 |
| SPEI12 | | | **0.37 *** | **0.00 – 0.73** | | | | |
| HSM × SPEI12 | | | 0.64 | -0.44 – 1.71 | | | | |
| nSR × SPEI12 | | | -0.09 | -0.19 – 0.01 | | | | |
| (HSM × nSR) × SPEI12 | | | -0.33 | -0.71 – 0.05 | | | | |
| SPEI6sept | | | | | 0.57 | -0.00 – 1.15 | | |
| HSM × SPEI6sept | | | | | 0.2 | -0.70 – 1.10 | | |
| nSR × SPEI6sept | | | | | -0.07 | -0.25 – 0.10 | | |
| (HSM × nSR) × SPEI6sept | | | | | -0.2 | -0.54 – 0.14 | | |
| REWmin | | | | | | | 2.42 | -0.17 – 5.01 |
| HSM × REWmin | | | | | | | -1.82 | -5.60 – 1.95 |
| nSR × REWmin | | | | | | | -0.44 | -1.24 – 0.36 |
| (HSM × nSR) × REWmin | | | | | | | -0.46 | -1.88 – 0.96 |
| **Random Effects** | | | | | | | | |
| $\sigma^2$ | 0.11 | | 0.11 | | 0.11 | | 0.11 | |

| | | | | |
|---|---|---|---|---|
| $\tau_{00}$ | 0.02 Plot:Site | 0.02 Plot:Site | 0.02 Plot:Site | 0.02 Plot:Site |
| | 0.03 Site | 0.03 Site | 0.02 Site | 0.03 Site |
| ICC | 0.33 | 0.31 | 0.29 | 0.32 |
| N | 69 Plot | 69 Plot | 69 Plot | 69 Plot |
| | 9 Site | 9 Site | 9 Site | 9 Site |
| Observations | 912 | 912 | 912 | 912 |
| Marginal $R^2$ / Conditional $R^2$ | 0.109 / 0.407 | 0.141 / 0.408 | 0.196 / 0.429 | 0.251 / 0.491 |
| AIC | 490.1 | 500.083 | 499.404 | 479.372 |

** p<0.05   ** p<0.01   *** p<0.001*

Table S9 (cont.)

| **Response BIOMinc** [log] | **m1b (FD$_{HSM}$ * HSM)** | | **m1b (FD$_{HSM}$ * HSM * SPEI12)** | | **m1b (FD$_{HSM}$ * HSM * SPEI6sept)** | | **m1b (FD$_{HSM}$ * HSM * REWmin)** | |
|---|---|---|---|---|---|---|---|---|
| *Predictors* | *Estimates* | *CI* | *Estimates* | *CI* | *Estimates* | *CI* | *Estimates* | *CI* |
| (Intercept) | 0.05 | -0.08 – 0.19 | 0.24 | -0.28 – 0.75 | **0.63** | **-0.13 – 1.40** | 0.04 | -0.08 – 0.17 |
| BA | **0.05 **** | **0.02 – 0.07** | **0.05 **** | **0.02 – 0.08** | **0.05 **** | **0.02 – 0.07** | **0.05 **** | **0.02 – 0.07** |
| Hegyi | 0.08 | -0.09 – 0.24 | 0.09 | -0.08 – 0.25 | 0.07 | -0.10 – 0.24 | 0.07 | -0.09 – 0.24 |
| HSM | **0.37 ***** | **0.24 – 0.49** | -0.07 | -1.39 – 1.26 | -0.44 | -1.59 – 0.71 | **0.28 ***** | **0.15 – 0.42** |
| FD$_{HSM}$ | 0.01 | -0.22 – 0.24 | 1.48 | -0.42 – 3.39 | 0.99 | -1.54 – 3.53 | 0.04 | -0.19 – 0.27 |
| HSM × FD$_{HSM}$ | -0.24 | -0.72 – 0.25 | 1.18 | -4.69 – 7.06 | 1.83 | -3.47 – 7.13 | -0.12 | -0.61 – 0.37 |
| SPEI12 | | | 0.09 | -0.19 – 0.37 | | | | |
| HSM × SPEI12 | | | -0.28 | -1.03 – 0.48 | | | | |
| FD$_{HSM}$ × SPEI12 | | | 0.9 | -0.26 – 2.07 | | | | |
| (FD$_{HSM}$ × HSM ) × SPEI12 | | | 0.95 | -2.46 – 4.36 | | | | |
| SPEI6sept | | | | | 0.32 | -0.13 – 0.76 | | |
| HSM × SPEI6sept | | | | | -0.48 | -1.17 – 0.20 | | |
| FD$_{HSM}$ × SPEI6sept | | | | | 0.62 | -0.97 – 2.20 | | |
| (FD$_{HSM}$ × HSM) × SPEI6sept | | | | | 1.27 | -2.01 – 4.55 | | |
| REWmin | | | | | | | 0.99 | -1.04 – 3.01 |
| HSM × REWmin | | | | | | | **-4.17 **** | **-6.73 – -1.61** |
| FD$_{HSM}$ × REWmin | | | | | | | 2.81 | -3.27 – 8.88 |
| (FD$_{HSM}$ × HSM) × REWmin | | | | | | | 11.59 | -0.98 – 24.15 |
| **Random Effects** | | | | | | | | |
| $\sigma^2$ | 0.11 | | 0.11 | | 0.11 | | 0.11 | |
| $\tau_{00}$ | 0.02 Plot:Site | | 0.03 Plot:Site | | 0.03 Plot:Site | | 0.03 Plot:Site | |

| | | | | |
|---|---|---|---|---|
| | 0.03 Site | 0.02 Site | 0.02 Site | 0.03 Site |
| ICC | 0.34 | 0.32 | 0.3 | 0.33 |
| N | 69 Plot | 69 Plot | 69 Plot | 69 Plot |
| | 9 Site | 9 Site | 9 Site | 9 Site |
| Observations | 912 | 912 | 912 | 912 |
| Marginal $R^2$ / Conditional $R^2$ | 0.112 / 0.413 | 0.164 / 0.431 | 0.206 / 0.448 | 0.256 / 0.499 |
| AIC | 483.624 | 487.35 | 485.035 | 462.452 |

** p<0.05 ** p<0.01 *** p<0.001*

*Table note: Response variables are log-transformed growth response in terms of biomass increment. Individual models were fitted separately to test different approaches of the effect of the neighborhood tree diversity in terms of species richness (nSR, m1a) and functional-trait diversity ($FD_{HSM}$, m1b). Tree size is represented by $BA_{std\_sp\_site}$ (standardized via scaling per species and site) and neighborhood competition is represented by $Hegyi_{std\_site\_minmax}$ (standardized via min-max normalization per site). $HSM_{TLP}$ was standardized by scaling-centering across the functional trait dataset of species-specific values including all focal tree and neighbor tree species. $FD_{HSM}$ is represented by the neighborhood functional diversity in terms of hydraulic safety and was standardized via min-max normalization across all sites to account for absolute differences between sites. Site-specific indices of drought intensity during the single-year drought were calculated in terms of annual SPEI12 December, SPEI6 September, and minimum monthly relative extractable water (REWmin) during the growing season. Values correspond with the estimates of regression coefficients and 95% confidence intervals (CI) for each predictor variable. Significant fixed effects with p<0.05 are highlighted in bold. Models highlighted in green are the most parsimonious (lower AIC) and with higher explanatory power (marginal $R^2$). Random effect variance is represented by the variance of the model residuals ($\sigma^2$), random intercept variance ($\tau_{00}$), intraclass correlation coefficient (ICC) and the number of groups/observations (N). Linear-mixed effects models were fitted using restricted maximum likelihood estimation (REML) with lme4 R-package (Bates et al. 2015). Table statistics were created using the sjPlot R-package (Lüdecke et al. 2024).*

Table S10. Comparison of mixed-effect model statistics for the growth responses (BIOMinc) during the first drought considering the modulating effects of competition (Hegyi) or tree size (BA) on the interaction between the drought tolerance (hydraulic safety margin $HSM_{TLP}$) of the focal-tree species and the neighborhood tree diversity (species richness nSR, or hydraulic trait diversity $FD_{HSM}$).

| **Response BIOMinc** [log] *Predictors* | **m1a (nSR * HSM)** *Estimates* | *CI* | **m1a (nSR * HSM * Hegyi)** *Estimates* | *CI* | **m1a (nSR * HSM * BA)** *Estimates* | *CI* |
|---|---|---|---|---|---|---|
| (Intercept) | 0 | -0.15 – 0.16 | 0.03 | -0.14 – 0.20 | 0.01 | -0.14 – 0.16 |
| BA | **0.04 ** ** | **0.02 – 0.07** | **0.04 ** ** | **0.02 – 0.07** | 0.08 * | 0.00 – 0.16 |
| Hegyi | 0.09 | -0.08 – 0.26 | -0.04 | -0.45 – 0.36 | 0.09 | -0.08 – 0.25 |
| HSM | **0.22 *** | **0.05 – 0.39** | **0.26 *** | **0.06 – 0.47** | **0.22 *** | **0.05 – 0.40** |
| nSR | 0.02 | -0.02 – 0.06 | 0.01 | -0.04 – 0.05 | 0.02 | -0.02 – 0.05 |
| nSR × HSM | 0.05 | -0.01 – 0.12 | 0.03 | -0.06 – 0.11 | 0.05 | -0.01 – 0.12 |
| HSM × Hegyi | | | -0.3 | -1.18 – 0.58 | | |
| nSR × Hegyi | | | 0.08 | -0.09 – 0.25 | | |
| (nSR × HSM) × Hegyi | | | 0.17 | -0.18 – 0.51 | | |
| HSM × BA | | | | | 0.02 | -0.12 – 0.16 |
| nSR × BA | | | | | -0.02 | -0.05 – 0.02 |
| (nSR × HSM) × BA | | | | | -0.01 | -0.07 – 0.05 |
| **Random Effects** | | | | | | |
| $\sigma^2$ | 0.11 | | 0.11 | | 0.11 | |
| $\tau_{00}$ | 0.02 Plot:Site<br>0.03 Site | | 0.02 Plot:Site<br>0.03 Site | | 0.02 Plot:Site<br>0.03 Site | |
| ICC | 0.33 | | 0.33 | | 0.33 | |
| N | 69 Plot<br>9 Site | | 69 Plot<br>9 Site | | 69 Plot<br>9 Site | |
| Observations | 912 | | 912 | | 912 | |
| Marginal $R^2$ / Conditional $R^2$ | 0.109 / 0.407 | | 0.109 / 0.404 | | 0.110 / 0.406 | |
| AIC | 490.1 | | 501.661 | | 512.045 | |

** p<0.05 ** p<0.01 *** p<0.001*

Table S10 (cont.)

| **Response BIOMinc** [log] *Predictors* | **m1b (FDis * HSM)** *Estimates* | *CI* | **m1b (FDis * HSM * Hegyi)** *Estimates* | *CI* | **m1b (FDis * HSM * BA)** *Estimates* | *CI* |
|---|---|---|---|---|---|---|
| (Intercept) | 0.05 | -0.08 – 0.19 | 0.06 | -0.08 – 0.20 | 0.05 | -0.08 – 0.19 |
| BA | **0.05 **** | **0.02 – 0.07** | **0.05 **** | **0.02 – 0.07** | **0.06 *** | **0.01 – 0.11** |
| Hegyi | 0.08 | -0.09 – 0.24 | -0.02 | -0.28 – 0.24 | 0.08 | -0.09 – 0.25 |
| HSM | **0.37 ***** | **0.24 – 0.49** | **0.37 ***** | **0.24 – 0.51** | **0.37 ***** | **0.24 – 0.49** |
| FDHSM | 0.01 | -0.22 – 0.24 | -0.12 | -0.41 – 0.17 | 0 | -0.24 – 0.23 |
| FDHSM × HSM | -0.24 | -0.72 – 0.25 | -0.44 | -1.06 – 0.17 | -0.29 | -0.81 – 0.23 |
| HSM × Hegyi | | | -0.15 | -0.68 – 0.37 | | |
| FDHSM × Hegyi | | | 1.23 | -0.24 – 2.69 | | |
| (FDHSM × HSM) × Hegyi | | | 2.28 | -1.39 – 5.95 | | |
| HSM × BA | | | | | 0.02 | -0.06 – 0.09 |
| FDHSM × BA | | | | | -0.11 | -0.32 – 0.09 |
| (FDHSM × HSM) × BA | | | | | -0.18 | -0.70 – 0.34 |
| **Random Effects** | | | | | | |
| $\sigma^2$ | 0.11 | | 0.11 | | 0.11 | |
| $\tau_{00}$ | 0.02 Plot:Site_code | | 0.02 Plot:Site_code | | 0.02 Plot:Site_code | |
| | 0.04 Site_code | | 0.04 Site_code | | 0.04 Site_code | |
| ICC | 0.39 | | 0.38 | | 0.39 | |
| N | 69 Plot | | 69 Plot | | 69 Plot | |
| | 9 Site_code | | 9 Site_code | | 9 Site_code | |
| Observations | 912 | | 912 | | 912 | |
| Marginal $R^2$ / Conditional $R^2$ | 0.105 / 0.452 | | 0.106 / 0.447 | | 0.105 / 0.452 | |
| AIC | 496.747 | | 498.67 | | 510.751 | |

** p<0.05   ** p<0.01   *** p<0.001*

*Table note: Response variables are log-transformed growth response in terms of biomass increment. Individual models were fitted separately to test different approaches of the effect of the neighborhood tree diversity in terms of species richness (nSR, m1a) and functional-trait diversity ($FD_{HSM}$, m1b). Tree size is represented by $BA_{std\_sp\_site}$ (standardized via scaling per species and site) and neighborhood competition is represented by $Hegyi_{std\_site\_minmax}$ (standardized via min-max normalization per site). $HSM_{TLP}$ was standardized by scaling-centering across the functional trait dataset of species-specific values including all focal tree and neighbor tree species. $FD_{HSM}$ is represented by the neighborhood functional diversity in terms of hydraulic safety and was standardized via min-max normalization across all sites to account for absolute differences between sites. Values correspond with the estimates of regression coefficients and 95% confidence intervals (CI) for each predictor variable. Significant fixed effects with p<0.05 are highlighted in bold. Random effect variance is represented by the variance of the model residuals ($\sigma^2$), random intercept variance ($\tau_{00}$), intraclass correlation coefficient (ICC) and the number of groups/observations (N). Linear-mixed effects models were fitted using restricted maximum* *likelihood estimation (REML) with lme4 R-package (Bates et al. 2015). Table statistics were created using the sjPlot R-package (Lüdecke et al. 2024).*

#### Growth responses during consecutive drought years

Table S11. Comparison of mixed-effect model statistics for the growth responses (BIOMinc) across consecutive drought years and for different tree-diversity experiments considering the interaction between the drought tolerance (hydraulic safety margin $HSM_{TLP}$) of the focal-tree species and the neighborhood tree diversity in terms of species richness (nSR).

**m2a Resp BIOMinc [log]**

| | F_G | | F_H | | F_Z | |
|---|---|---|---|---|---|---|
| *Predictors* | *Estimates* | *CI* | *Estimates* | *CI* | *Estimates* | *CI* |
| BA | **0.12** * | 0.00 – 0.23 | 0.09 | -0.01 – 0.18 | **0.17** *** | 0.08 – 0.26 |
| Hegyi | -0.05 | -0.71 – 0.61 | -0.42 | -0.89 – 0.04 | -0.03 | -0.98 – 0.92 |
| nSR | 0.14 | -0.06 – 0.35 | 0.13 | -0.04 – 0.31 | 0.12 | -0.01 – 0.25 |
| HSM | 0.2 | -0.72 – 1.12 | -0.38 | -1.12 – 0.36 | 0.08 | -0.34 – 0.49 |
| Resp yr [1] | 0 | -0.49 – 0.49 | -0.33 | -0.80 – 0.14 | -0.22 | -0.54 – 0.10 |
| Resp yr [2] | -0.02 | -0.52 – 0.47 | 0.22 | -0.25 – 0.69 | -0.16 | -0.48 – 0.16 |
| Resp yr [3] | -0.37 | -0.86 – 0.12 | -0.31 | -0.78 – 0.16 | -0.04 | -0.36 – 0.28 |
| Resp yr [4] | 0.16 | -0.33 – 0.65 | **0.74** ** | 0.27 – 1.21 | -0.09 | -0.41 – 0.24 |
| nSR × HSM | 0.11 | -0.27 – 0.49 | 0.24 | -0.05 – 0.53 | 0.13 | -0.04 – 0.30 |
| nSR × Resp yr [2] | 0.01 | -0.10 – 0.12 | -0.04 | -0.12 – 0.03 | -0.03 | -0.09 – 0.04 |
| nSR × Resp yr [3] | 0 | -0.11 – 0.12 | -0.03 | -0.10 – 0.04 | **-0.10** ** | -0.17 – -0.03 |
| nSR × Resp yr [4] | 0.05 | -0.07 – 0.16 | -0.05 | -0.12 – 0.02 | **-0.11** ** | -0.18 – -0.04 |
| HSM × Resp yr [2] | 0.26 | -0.07 – 0.60 | 0.22 | -0.09 – 0.53 | **-0.25** ** | -0.41 – -0.09 |
| HSM × Resp yr [3] | -0.08 | -0.41 – 0.26 | 0 | -0.32 – 0.31 | **-0.28** *** | -0.44 – -0.12 |
| HSM × Resp yr [4] | **1.01** *** | 0.67 – 1.34 | **0.82** *** | 0.50 – 1.13 | -0.14 | -0.30 – 0.02 |
| **Random Effects** | | | | | | |
| $\sigma^2$ | 0.12 | | 0.12 | | 0.13 | |
| $\tau_{00}$ | 0.22 $_{Tree\_ID}$ | | 0.19 $_{Tree\_ID}$ | | 0.24 $_{Tree\_ID}$ | |
| ICC | 0.64 | | 0.62 | | 0.65 | |
| N | 108 $_{Tree\_ID}$ | | 155 $_{Tree\_ID}$ | | 182 $_{Tree\_ID}$ | |
| Observations | 432 | | 620 | | 728 | |
| Marginal $R^2$ / Conditional $R^2$ | 0.214 / 0.713 | | 0.240 / 0.708 | | 0.122 / 0.693 | |
| AIC | 559.585 | | 830.233 | | 868.378 | |

* p<0.05 ** p<0.01 *** p<0.001

Table S11 (cont.)

**m2a Resp BIOMinc [log]**

| | K | | I_F | | S | |
|---|---|---|---|---|---|---|
| *Predictors* | *Estimates* | *CI* | *Estimates* | *CI* | *Estimates* | *CI* |
| BA | **0.09** * | 0.01 – 0.17 | **0.14** * | 0.01 – 0.26 | -0.01 | -0.10 – 0.08 |
| Hegyi | **0.62** * | 0.13 – 1.10 | -0.49 | -1.29 – 0.31 | -0.19 | -0.74 – 0.36 |
| nSR | -0.03 | -0.23 – 0.16 | -0.01 | -0.15 – 0.14 | 0.68 | -0.17 – 1.54 |
| HSM | 0.51 | -0.15 – 1.18 | 0.34 | -1.20 – 1.88 | **-3.48** * | -6.73 – -0.24 |
| Resp yr [1] | -0.3 | -0.72 – 0.12 | 0.13 | -0.31 – 0.57 | **-2.09** * | -3.99 – -0.18 |
| Resp yr [2] | **-0.61** ** | -1.03 – -0.19 | -0.17 | -0.61 – 0.27 | **-2.19** * | -4.09 – -0.28 |
| Resp yr [3] | **-1.39** *** | -1.81 – -0.97 | **-0.61** ** | -1.05 – -0.17 | -1.77 | -3.68 – 0.13 |
| Resp yr [4] | **-0.99** *** | -1.41 – -0.57 | -0.09 | -0.53 – 0.35 | -0.98 | -2.89 – 0.92 |
| nSR × HSM | -0.03 | -0.33 – 0.26 | 0.02 | -0.57 – 0.61 | 1.16 | -0.29 – 2.61 |
| nSR × Resp yr [2] | -0.06 | -0.14 – 0.03 | **0.19** *** | 0.12 – 0.26 | -0.03 | -0.09 – 0.03 |
| nSR × Resp yr [3] | -0.03 | -0.11 – 0.05 | **0.18** *** | 0.11 – 0.25 | -0.04 | -0.10 – 0.03 |
| nSR × Resp yr [4] | **-0.09** * | -0.18 – -0.01 | **0.26** *** | 0.19 – 0.33 | **-0.07** * | -0.13 – -0.00 |
| HSM × Resp yr [2] | **-1.13** *** | -1.42 – -0.84 | 0.67 | -0.16 – 1.49 | -0.23 | -1.36 – 0.90 |
| HSM × Resp yr [3] | **-2.04** *** | -2.33 – -1.75 | **1.62** *** | 0.80 – 2.45 | 0.42 | -0.71 – 1.55 |
| HSM × Resp yr [4] | **-1.73** *** | -2.02 – -1.44 | **1.50** *** | 0.67 – 2.32 | **1.67** ** | 0.54 – 2.80 |
| **Random Effects** | | | | | | |
| $\sigma^2$ | 0.13 | | 0.11 | | 0.04 | |
| $\tau_{00}$ | 0.14 $_{Tree\_ID}$ | | 0.16 $_{Tree\_ID}$ | | 0.10 $_{Tree\_ID}$ | |
| ICC | 0.52 | | 0.59 | | 0.69 | |
| N | 179 $_{Tree\_ID}$ | | 88 $_{Tree\_ID}$ | | 100 $_{Tree\_ID}$ | |
| Observations | 716 | | 352 | | 400 | |
| Marginal $R^2$ / Conditional $R^2$ | 0.254 / 0.640 | | 0.422 / 0.762 | | 0.058 / 0.712 | |
| AIC | 298.126 | | 707.172 | | 108.803 | |

* p<0.05 ** p<0.01 *** p<0.001

*Table note: Response variables are log-transformed growth response in terms of biomass increment. Individual models were fitted separately for each experiment. Tree size is represented by $BA_{std\_sp\_site}$ (standardized per species and site) and neighborhood competition is represented by $Hegyi_{std\_site\_minmax}$ (standardized via min-max normalization per site). $HSM_{TLP}$ was standardized by scaling-centering across the functional trait dataset of species-specific values including all focal tree and neighbor tree species. $FD_{HSM}$ is represented by the neighborhood functional diversity in terms of hydraulic safety and was standardized via min-max normalization across all sites to account for absolute differences between sites. Values correspond with the estimates of regression coefficients and 95% confidence intervals (CI) for each predictor variable. Significant fixed effects with p<0.05 are highlighted in bold, with positive significant effects highlighted in blue and negative significant effects in orange. Random effect variance is represented by the variance of the model residuals ($\sigma^2$), random intercept variance ($\tau_{00}$), intraclass correlation coefficient (ICC) and the number of observations as tree samples. Linear-mixed effects models were fitted using restricted maximum likelihood estimation (REML) with lme4 R-*

*package (Bates et al. 2015). Table statistics were created using the sjPlot R-package (Lüdecke et al. 2024).*

Table S12. Comparison of mixed-effect model statistics for the growth responses (BIOMinc) across consecutive drought years and for different tree-diversity experiments considering the interaction between the drought tolerance (hydraulic safety margin $HSM_{TLP}$) of the focal-tree species and the neighborhood tree diversity in terms of hydraulic trait diversity ($FD_{HSM}$).

**m2b Resp BIOMinc [log]**

| | **F_G** | | **F_H** | | **F_Z** | |
|---|---|---|---|---|---|---|
| *Predictors* | *Estimates* | *CI* | *Estimates* | *CI* | *Estimates* | *CI* |
| $BA_{std\_sp\_site}$ | 0.11 | -0.00 – 0.23 | 0.08 | -0.02 – 0.18 | **0.12** ** | 0.03 – 0.22 |
| $Hegyi_{std\_site\_minmax}$ | -0.06 | -0.73 – 0.62 | -0.34 | -0.79 – 0.12 | -0.27 | -1.19 – 0.64 |
| $FD_{HSM\_minmax}$ | 1.58 | -0.18 – 3.35 | 0.4 | -0.78 – 1.57 | 0.04 | -0.49 – 0.58 |
| HSM | 0.32 | -0.25 – 0.88 | -0.06 | -0.58 – 0.46 | **0.29** * | 0.02 – 0.56 |
| Resp yr [1] | 0.14 | -0.17 – 0.45 | -0.11 | -0.44 – 0.23 | 0.03 | -0.20 – 0.26 |
| Resp yr [2] | 0.18 | -0.13 – 0.49 | **0.42** * | 0.08 – 0.76 | 0.16 | -0.07 – 0.38 |
| Resp yr [3] | -0.13 | -0.44 – 0.18 | -0.02 | -0.36 – 0.32 | **0.29** * | 0.06 – 0.52 |
| Resp yr [4] | **0.45** ** | 0.14 – 0.76 | **0.80** *** | 0.46 – 1.13 | **0.27** * | 0.04 – 0.49 |
| $FD_{HSM}$ × HSM sc | 0.94 | -2.57 – 4.45 | 1.84 | -0.71 – 4.39 | 0.41 | -0.53 – 1.36 |
| $FD_{HSM}$ × Resp yr [2] | -0.52 | -1.49 – 0.45 | -0.42 | -1.04 – 0.21 | **-0.43** * | -0.85 – -0.00 |
| $FD_{HSM}$ × Resp yr [3] | **-1.03** * | -2.00 – -0.06 | -0.61 | -1.23 – 0.02 | **-1.07** *** | -1.49 – -0.65 |
| $FD_{HSM}$ × Resp yr [4] | -0.73 | -1.70 – 0.24 | 0.18 | -0.45 – 0.80 | **-1.26** *** | -1.68 – -0.84 |
| HSM × Resp yr [2] | 0.26 | -0.07 – 0.60 | 0.27 | -0.06 – 0.60 | -0.16 | -0.34 – 0.01 |
| HSM × Resp yr [3] | -0.07 | -0.40 – 0.27 | 0.1 | -0.23 – 0.43 | -0.06 | -0.24 – 0.11 |
| HSM × Resp yr [4] | **1.01** *** | 0.67 – 1.34 | **0.75** *** | 0.42 – 1.08 | 0.11 | -0.07 – 0.29 |
| **Random Effects** | | | | | | |
| $\sigma^2$ | 0.12 | | 0.12 | | 0.12 | |
| $\tau_{00}$ | 0.22 $_{Tree\_ID}$ | | 0.19 $_{Tree\_ID}$ | | 0.23 $_{Tree\_ID}$ | |
| ICC | 0.64 | | 0.62 | | 0.65 | |
| N | 108 $_{Tree\_ID}$ | | 155 $_{Tree\_ID}$ | | 182 $_{Tree\_ID}$ | |
| Observations | 432 | | 620 | | 728 | |
| Marginal $R^2$ / Conditional $R^2$ | 0.206 / 0.717 | | 0.247 / 0.711 | | 0.159 / 0.708 | |
| AIC | 536.048 | | 801.561 | | 815.519 | |

** p<0.05   ** p<0.01   *** p<0.001*

Table S12 (cont.)

**m2b Resp BIOMinc [log]**

| | **K** | | **I_F** | | **S** | |
|---|---|---|---|---|---|---|
| *Predictors* | *Estimates* | *CI* | *Estimates* | *CI* | *Estimates* | *CI* |
| $BA_{std\_sp\_site}$ | 0.08 | -0.00 – 0.16 | 0.12 | -0.00 – 0.25 | -0.01 | -0.11 – 0.09 |
| $Hegyi_{std\_site\_minmax}$ | **0.53** * | 0.05 – 1.00 | -0.69 | -1.49 – 0.12 | -0.24 | -0.75 – 0.28 |
| $FD_{HSM\ \_minmax}$ | 0.32 | -0.96 – 1.59 | 0.38 | -1.32 – 2.08 | 0.02 | -7.00 – 7.05 |
| HSM | 0.37 | -0.05 – 0.79 | 0.49 | -0.67 – 1.65 | -1.14 | -3.02 – 0.73 |
| Resp yr [1] | **-0.40** ** | -0.69 – -0.11 | 0.09 | -0.26 – 0.44 | -0.73 | -1.87 – 0.41 |
| Resp yr [2] | **-0.84** *** | -1.13 – -0.54 | 0.08 | -0.27 – 0.43 | -0.83 | -1.97 – 0.32 |
| Resp yr [3] | **-1.53** *** | -1.82 – -1.24 | **-0.35** * | -0.70 – -0.01 | -0.38 | -1.52 – 0.76 |
| Resp yr [4] | **-1.08** *** | -1.37 – -0.79 | 0.28 | -0.07 – 0.63 | 0.44 | -0.70 – 1.58 |
| $FD_{HSM}$ × HSM sc | 0.62 | -1.76 – 2.99 | -0.83 | -7.13 – 5.47 | -0.4 | -12.71 – 11.91 |
| $FD_{HSM}$ × Resp yr [2] | 0.11 | -0.68 – 0.91 | **1.44** *** | 0.70 – 2.18 | -0.25 | -0.64 – 0.13 |
| $FD_{HSM}$ × Resp yr [3] | -0.09 | -0.88 – 0.70 | **1.22** ** | 0.48 – 1.96 | **-0.45** * | -0.83 – -0.07 |
| $FD_{HSM}$ × Resp yr [4] | **-1.04** * | -1.84 – -0.25 | **2.02** *** | 1.28 – 2.76 | **-0.79** *** | -1.17 – -0.40 |
| HSM × Resp yr [2] | **-1.13** *** | -1.45 – -0.82 | 0.6 | -0.27 – 1.47 | -0.17 | -1.28 – 0.95 |
| HSM × Resp yr [3] | **-2.02** *** | -2.33 – -1.71 | **1.53** *** | 0.66 – 2.40 | 0.53 | -0.58 – 1.64 |
| HSM × Resp yr [4] | **-1.56** *** | -1.87 – -1.25 | **1.40** ** | 0.53 – 2.27 | **1.86** ** | 0.75 – 2.98 |
| **Random Effects** | | | | | | |
| $\sigma^2$ | 0.13 | | 0.12 | | 0.04 | |
| $\tau_{00}$ | 0.14 $_{Tree\_ID}$ | | 0.16 $_{Tree\_ID}$ | | 0.10 $_{Tree\_ID}$ | |
| ICC | 0.52 | | 0.58 | | 0.71 | |
| N | 179 $_{Tree\_ID}$ | | 88 $_{Tree\_ID}$ | | 100 $_{Tree\_ID}$ | |
| Observations | 716 | | 352 | | 400 | |
| Marginal $R^2$ / Conditional $R^2$ | 0.251 / 0.644 | | 0.385 / 0.739 | | 0.047 / 0.725 | |
| AIC | 272.764 | | 712.405 | | 79.88 | |

* p<0.05   ** p<0.01   *** p<0.001

*Table note: Response variables are log-transformed growth response in terms of biomass increment. Individual models were fitted separately for each experiment. Tree size is represented by $BA_{std\_sp\_site}$ (standardized per species and site) and neighborhood competition is represented by $Hegyi_{std\_site\_minmax}$ (standardized via min-max normalization per site). $HSM_{TLP}$ was standardized by scaling-centering across the functional trait dataset of species-specific values including all focal tree and neighbor tree species. $FD_{HSM}$ is represented by the neighborhood functional diversity in terms of hydraulic safety and was standardized via min-max normalization across all sites to account for absolute differences between sites. Values correspond with the estimates of regression coefficients and 95% confidence intervals (CI) for each predictor variable. Significant fixed effects with p<0.05 are highlighted in bold, with positive significant effects highlighted in blue and negative significant effects in orange. Random effect variance is represented by the variance of the model residuals ($\sigma^2$), random intercept variance ($\tau_{00}$), intraclass correlation coefficient (ICC) and the number of observations as tree samples (N). Linear-mixed effects models were fitted using restricted maximum likelihood estimation (REML) with lme4 R-*

*package (Bates et al. 2015). Table statistics were created using the sjPlot R-package (Lüdecke et al. 2024).*
